# Self-generated cues and learned landmarks support spatial learning in *Drosophila*

**DOI:** 10.1101/2024.09.28.615625

**Authors:** Yang Chen, Robert Alfredson, Xiaofan Zuo, Dorsa Motevalli, Sydney Fogleman, Ulrich Stern, Chung-Hui Yang

## Abstract

The ability to return to remembered goal locations is essential for animal survival. Although animals are known to use visual landmarks, how well they can exploit non-visual cues for spatial learning remains poorly understood. Here we show that *Drosophila* can combine learned olfactory cues with self-motion information to return to initially unmarked goals. In feature-poor arenas, flies assign positive value to self-deposited scents near the rewarded goal and combine these cues with self-motion cues to guide their return. This strategy depends on the mushroom body, an olfactory learning center that associates odors with reinforcement^1^, and PFN neurons, which encode egocentric translational velocity^2,3^. External olfactory landmarks away from the goal improve performance: flies assign them negative value, again through the mushroom body, and use them as aversive guideposts to constrain search area. Comparisons among wild-caught strains suggest that the capacities to learn in feature-poor arenas and to benefit from external olfactory landmarks are genetically separable. Complementary transcriptomic analyses reveal differences in learning-associated gene expression in feature-poor and landmark-enriched arenas. Our findings reveal that flies can construct and combine complementary spatial information to navigate without prominent visual cues and that the ability to learn in different sensory contexts is under partly distinct genetic control.

## Introduction

The ability to return to remembered goal locations without guidance from prominent visual landmarks is critical for animal survival, particularly for nocturnal animals and species with limited vision. Defining the behavioral, neural, and genetic mechanisms that support navigation under such conditions can provide insight into basic principles of spatial cognition. One strategy is path integration, in which animals integrate self-motion information to estimate their displacement from a remembered goal and thereby compute a home vector specifying the direction and distance back to it^4,5^. For example, desert ants have been shown to use step-based odometry to estimate distance traveled^6^. Evidence for path integration has also been reported in mammals such as rats^7^, mice^8^, bats^9^, and dogs^10^, and insects such as bees^11–14^, dung beetles^15–17^, and fruit flies^18–21^.

Olfactory cues provide another major source of spatial information when prominent visual cues are unavailable. Odors can aid successful navigation to goals in at least two ways. First, animals can locate a goal by following the odor plume if the goal emits innately attractive odors, such as those from a food source or attractive conspecifics^22–27^. Second, animals can use odors associated with fixed locations away from the goal to recalibrate the path integrator^28–30^. For example, mice can use odors delivered at specific locations along a linear path to estimate their position relative to the goal^30^. How animals combine self-motion information with olfactory landmarks during spatial learning, and the genetic mechanisms that control this capacity, remain poorly understood.

Previous work has shown that *Drosophila* can use visual landmarks for spatial learning^31,32^. Here we asked how *Drosophila* learn to return to a goal when prominent visual landmarks are unavailable. We developed a task in which flies receive an optogenetic reward upon entering an initially unmarked goal region at the center of a feature-poor arena. We found that flies learn to find self-deposited scents near the goal attractive and use these scents together with self-motion information to guide their returns. When external olfactory landmarks are introduced away from the goal, flies learn to treat them as aversive guideposts that constrain search and improve performance. Comparisons among F_1_ progeny of wild-derived strains suggest that the capacities to learn in feature-poor arenas and to exploit external olfactory landmarks are both genetically influenced and partly separable. Complementing these behavioral findings, RNA-seq revealed learning-associated transcriptional changes, some shared across arena types and others dependent on arena features.

Together, these findings reveal how a compact brain learns the spatial significance of non-visual external cues and integrates them with self-motion information to navigate without prominent visual landmarks, while providing a framework for dissecting the genetic and molecular mechanisms underlying spatial learning.

## Results

### Flies can learn the location of a spatial goal without relying on prominent visual cues

To test whether flies can learn a spatial goal location without relying on prominent visual cues, we designed a task where a fly can collect virtual rewards at a goal location in a feature-poor 10 × 16 mm arena (“flat chamber”) that lacks prominent visual landmarks (**Fig. 1a**). The task was administered using our custom-designed high-throughput SkinnerTrax^33^ system for behavior-dependent optogenetic stimulation of many flies independently and in parallel. Each apparatus houses 40 individual flies, each in a separate arena (**Fig. 1b**). We customized SkinnerTrax to place a circular goal at the center of each arena, a location not directly associated with specific spatial features such as arena walls (**Fig. 1a**). Each time the fly crossed over the software-defined border of the reward circle *from the outside*, a 250-ms pulse of light was delivered to optogenetically activate a group of neurons in the brain labeled by *0273-GAL4* (**Supplementary Video 1**). Previous studies have shown that flies robustly seek optogenetic activation of *0273-GAL4*-labeled neurons^34,35^; we therefore used this stimulation as a potent reinforcer in our task. The key populations of neurons labeled by *0273-GAL4* include the reward-mediating PAM class of dopaminergic (DA) neurons that project to the mushroom body (MB) and the *fruitless*-expressing P1 neurons known to signal strongly to PAM-DA neurons^36–38^ (**Extended Data Fig. 1**). We followed learning over two 1-h Training sessions, preceded by a 15-min Pre period and each followed by a 15-min Post period (**Fig. 1d**). Rewards were unavailable in either Pre or Post periods. Two features of our design are noteworthy. First, to collect additional rewards following receipt of a reward, the fly must exit the reward circle and reenter it from the outside, as staying inside will not trigger any rewards (**Fig. 1c**). This design should push flies to solve the task continuously. Second, we paired each experimental fly with a yoked control fly that received a reward at the same time as the experimental fly; consequently, the yoked control’s position had no influence over reward delivery (**Fig. 1d**). The yoked controls therefore provide a baseline for behavioral effects of reward receipt without spatial learning.

**Fig. 1.**
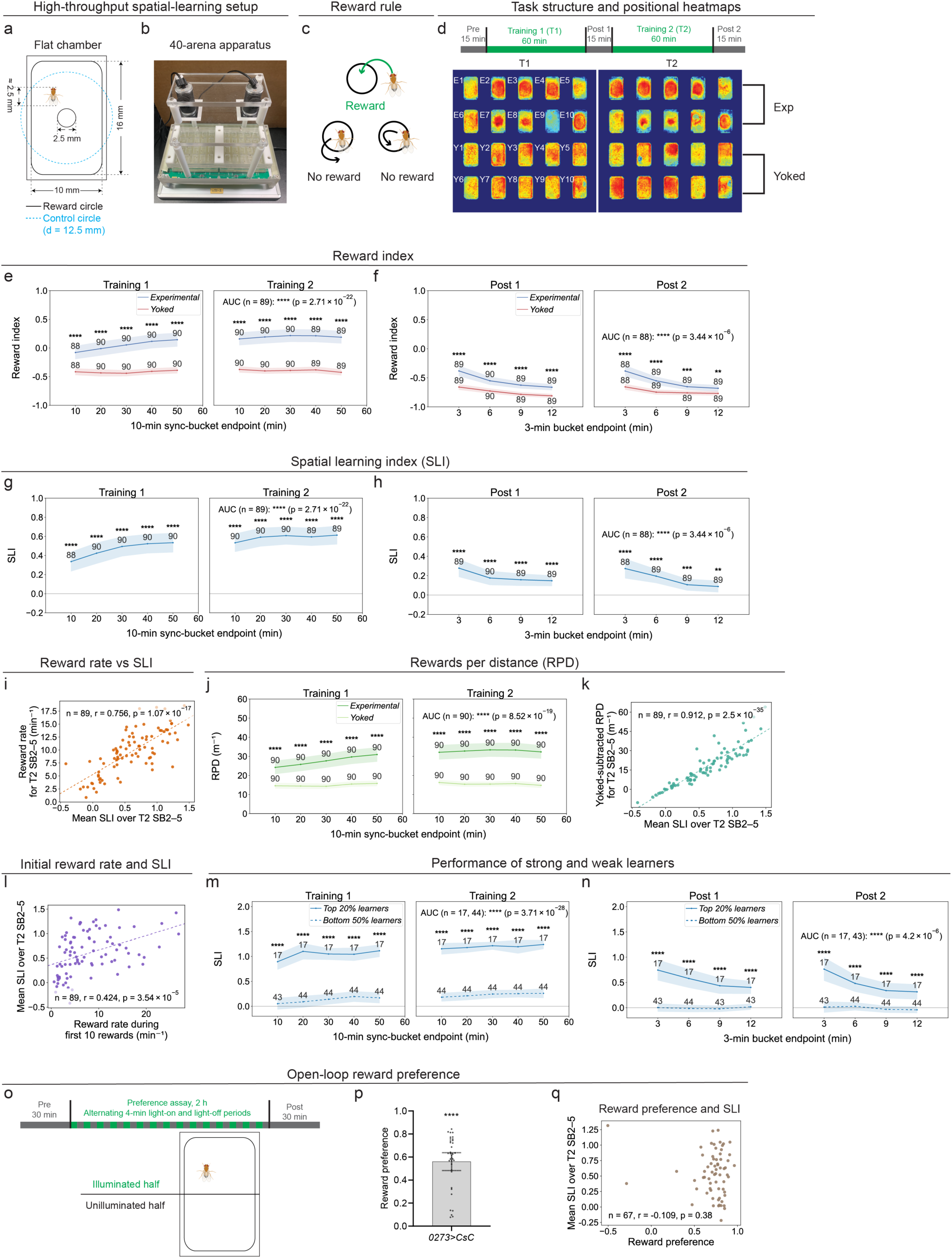
Flies can learn a spatial goal without prominent visual landmarks. a, Schematic of the flat arena (16 × 10 × 3 mm) for single flies. **b,** High-throughput SkinnerTrax setup for assaying 40 flies in parallel. **c,** During Training, an experimental fly triggers a 250-ms light pulse upon entering the reward circle from outside. **d,** Top, task timeline. Bottom, positional heatmaps of 10 experimental (E) and yoked (Y) pairs; matching numbers identify pairs receiving light pulses simultaneously. **e,f,** Reward index (RI) of experimental and yoked flies in 10-min sync buckets (SB) during Training (**e**) and 3-min buckets during Post (**f**), as defined in **Methods**. **g,h,** Spatial learning index (SLI; experimental RI minus paired yoked RI) during Training (**g**) and Post (**h**). **i,** Reward rate versus mean SLI, both for Training 2 (T2) SB2–5. **j,** Rewards per distance (RPD) of experimental and yoked flies during Training. **k,** Yoked-subtracted RPD versus mean SLI, both for T2 SB2–5. **l,** Mean SLI over T2 SB2–5 versus reward rate during the first 10 rewards. **m,n,** SLI of strong (top 20%) and weak (bottom 50%) learners during Training (**m**) and Post (**n**), classified by mean SLI over T2 SB2–5. **o,** Open-loop reward-preference assay for **p**: timeline (top), illumination pattern (bottom). **p,** Reward preference (*n* = 36). **q,** Mean SLI over T2 SB2–5 versus reward preference after training (shorter assay; **Methods**). Flies were *0273>CsChrimson*; **g–n** use the cohort in **e,f**. Time courses and the bar in **p** show means; shading and error bars indicate 95% confidence intervals. Points in **i,k,l,p,q** represent individual flies; numbers denote sample sizes. Two-sided tests: paired *t*-tests (**e,f,j**), one-sample *t*-tests against zero (**g,h,p**), Welch’s *t*-tests (**m,n**), and Pearson correlations (**i,k,l,q**); area-under-the-curve (AUC) comparisons use corresponding tests (**Methods**). Per-bucket *p* values are unadjusted. \**p* < 0.05; \*\**p* < 0.01; \*\*\**p* < 0.001; \*\*\*\**p* < 0.0001.

To quantify spatial learning beyond the positional heatmaps generated by SkinnerTrax (**Fig. 1d**), we first used the reward index (RI), which measures flies’ preference for entering the reward circle relative to a larger control circle (**Fig. 1a, e**). The calculation and rationale for this metric are described in **Methods**. The RI of experimental flies increased as training progressed and declined in the Post periods, whereas the RI of yoked controls stayed relatively constant during training (**Fig. 1e, f**). The higher RI of yoked controls during the Training sessions than during the Post periods shows that RI can capture behavioral effects of reward receipt without spatial learning. To account for these non-learning effects, we calculated the RI *difference* between each experimental fly and its paired yoked control, which we termed the spatial learning index (SLI) (**Fig. 1g**, **Methods**). The SLI increased as training progressed and declined in the Post periods (**Fig. 1g, h**, **Extended Data Fig. 2c-e**). To further evaluate SLI as a measure of learning performance, we correlated the SLI of individual flies with additional performance measures. A fly’s SLI correlated well with the rate at which it gathered rewards (**Fig. 1i**, **Extended Data Fig. 3a**). SLI correlated even more strongly with rewards per distance (RPD), which measures rewards collected per distance traveled, and with yoked-subtracted RPD (**Fig. 1j, k**, **Extended Data Fig. 3e**, **and Methods;** see **Supplementary Tables 1and 2** for reward rates and yoked-subtracted RPD across genotypes and experimental conditions, respectively). Importantly, SLI did not detectably correlate with reward preference in an open-loop assay (**Fig. 1o-q**), suggesting that differences in SLI are not simply explained by differences in reward-seeking motivation. We also detected no correlation between SLI and Pre-period speed (**Extended Data Fig. 3d**), consistent with RI and SLI being designed to minimize sensitivity to differences in walking speed (**Methods**).

**Table 1.** List of Putative Observed Compounds identified by GC-MS.

| **Retention Time (min)** | **Compound ID** | **Formula** | **Exp Tracking Mass (ug)** | **Yoked Tracking Mass (ug)** |
|---|---|---|---|---|
| 19.1 | Tricosene isomer | C<sub>23</sub>H<sub>46</sub> | 0.28 | 0.13 |
| 19.2 | Tricosene isomer | C<sub>23</sub>H<sub>46</sub> | 3.11 | 1.53 |
| 19.4 | Tricosene isomer | C<sub>23</sub>H<sub>46</sub> | 0.20 | 0.10 |
| 19.5 | Tricosene isomer | C<sub>23</sub>H<sub>46</sub> | 0.78 | 0.44 |
| 20.6 | Cyclotetracosane | C<sub>24</sub>H<sub>48</sub> | 0.18 | 0.09 |
| 21.9 | Pentacosene isomer | C<sub>25</sub>H<sub>50</sub> | 0.12 | 0.08 |
| 22.0 | Pentacosene isomer | C<sub>25</sub>H<sub>50</sub> | 0.65 | 0.46 |
| 22.3 | Pentacosene isomer | C<sub>25</sub>H<sub>50</sub> | 0.08 | 0.05 |

**Table 2.** DEGs of exp vs. yoked flies in the flat chamber, *w^1118^* background.

| Gene | padj | pvalue | log2FoldChange | baseMean |
|---|---|---|---|---|
| *DnaJ-1* | 1.42025E-14 | 1.26E-18 | -1.0439 | 6682.895 |
| *Hsp70Bc* | 4.8914E-12 | 9.75E-16 | -3.94163 | 350.6932 |
| *Hsp70Ba* | 4.8914E-12 | 1.3E-15 | -4.22101 | 200.3317 |
| *Hsp70Bb* | 1.27653E-11 | 4.51E-15 | -3.89844 | 1280 |
| *pirk* | 2.06411E-11 | 9.12E-15 | -1.93135 | 622.4805 |
| *Hsp70Bbb* | 2.53285E-11 | 1.34E-14 | -4.03736 | 602.9822 |
| *Hsp70Aa* | 1.63012E-10 | 1.01E-13 | -4.18315 | 462.6835 |
| *Hsp27* | 2.37949E-10 | 1.68E-13 | -2.36874 | 374.4291 |
| *Hsp70Ab* | 4.40336E-10 | 3.5E-13 | -4.0476 | 1490.373 |
| *Hsp26* | 7.01872E-10 | 6.2E-13 | -3.119 | 1159.298 |
| *Hsp68* | 8.41304E-10 | 8.18E-13 | -2.34835 | 2437.231 |
| *Hsp23* | 2.42624E-08 | 2.57E-11 | -1.3924 | 662.2332 |
| *CG13305* | 3.33108E-06 | 3.83E-09 | -0.74576 | 141.5543 |
| *CR45330* | 3.44361E-05 | 4.26E-08 | -1.06664 | 29.82355 |
| *CG12262* | 5.85308E-05 | 7.76E-08 | 1.314994 | 22.79654 |
| *CG31176* | 0.000412448 | 5.83E-07 | -0.28443 | 294.2922 |
| *CG33229* | 0.00070045 | 1.05E-06 | -0.37418 | 1272.426 |
| *stv* | 0.001517042 | 2.41E-06 | -0.55614 | 4691.085 |
| *Cpr66D* | 0.003299178 | 5.54E-06 | -0.80566 | 62.91581 |
| *CR45938* | 0.013722479 | 2.43E-05 | -1.92449 | 5.105216 |
| *CG6441* | 0.018605362 | 3.45E-05 | -0.59963 | 51.12128 |
| *Peb* | 0.021619805 | 4.2E-05 | -9.88456 | 78.49728 |
| *armi* | 0.029862309 | 6.33E-05 | -0.29145 | 238.8973 |
| *CG15531* | 0.029862309 | 6.29E-05 | -2.55043 | 5.822389 |
| *mid* | 0.045768138 | 0.000101 | -0.30417 | 221.4282 |

Note that **Tables 2-5** show only genes with padj < 0.05 and are sorted by padj.

We next used SLI to examine several features of how the flies learned. First, blind (*norpA* mutant) flies exhibited clear positive SLI over the course of training (**Extended Data Fig. 2d, e**), suggesting the task can be solved without vision. Second, the extent of a fly’s arena exploration during the Pre period did not detectably correlate with its subsequent learning performance (**Extended Data Fig. 3b**). Third, a fly’s SLI early in Training 1 predicted its SLI over the course of Training 2 (**Extended Data Fig. 3c**). In fact, how quickly a fly collected its first ten rewards already showed some predictive power for how well it learned during Training 2 (**Fig. 1l**). Consistent with these correlations, flies classified by their Training 2 performance as strong (top 20%) or weak (bottom 50%) learners showed an early and significant separation in SLI (**Fig. 1m, n**).

Together, these results show that flies can learn to return to a reward-baited goal in an arena lacking prominent visual landmarks, and that blind flies can solve the task. Individual differences in performance emerged early and predicted later performance, a pattern consistent with contributions from early reward history and/or pre-existing individual differences among flies. By contrast, Pre-period arena exploration, open-loop reward preference, and Pre-period walking speed showed no detectable association with subsequent learning performance.

### Self-motion cues play a role in supporting spatial learning in the absence of prominent visual cues

To identify the cues flies might use for returning to the unmarked goal in the absence of prominent visual landmarks, we first tested the potential involvement of self-motion information. Previous studies have suggested that flies can perform *egocentric* path integration, a strategy where animals rely solely on self-motion cues to calculate a home vector, which specifies the relative direction and distance to a goal, and then use the home vector to return to the goal^19–21^. PFN neurons are prime candidates for providing critical input to the computation of such a vector. They have been shown to track a fly’s egocentric translational velocity and comprise four subgroups – PFNd-R, PFNd-L, PFNv-R, and PFNv-L – each defined by its distinct self-motion selectivity^2,3^. For example, PFNd-R neurons are maximally tuned to forward movement in the direction ∼31° to the right of the fly’s body axis. The population activities of the four PFN subgroups are combined, via vector addition, to compute a fly’s allocentric translational velocity^2,3^. Moreover, perturbing PFNd neurons has been shown to impair the spatial accuracy of local search^3^.

To test the role of PFN neurons, we first chronically silenced the activities of PFNd neurons by expressing the Kir2.1 channel in them and found that learning decreased significantly (**Fig. 2a, b**). A learning deficit was also apparent when we transiently manipulated PFN neurons at reward delivery, using the same 250-ms pulse that delivered the optogenetic reward, by transiently silencing PFNd neurons with GtACR1 or by transiently activating PFNd or PFNv neurons with CsChrimson (**Fig. 2c-f**). Note that simultaneously activating the entire population of either PFNd or PFNv neurons is expected to disrupt their function, as such activation should compromise their spatial selectivity. The learning deficits in these experiments were unlikely to reflect altered reward seeking or walking speed (**Extended Data Fig. 4a-h**). It is also noteworthy that, by design, the SLI tends to suppress potential non-learning behavioral effects of various manipulations. This is a direct consequence of the SLI’s being calculated as the RI *difference* between each experimental fly and its paired yoked control, both of which underwent the same manipulation and hence likely share such effects.

**Fig. 2.**
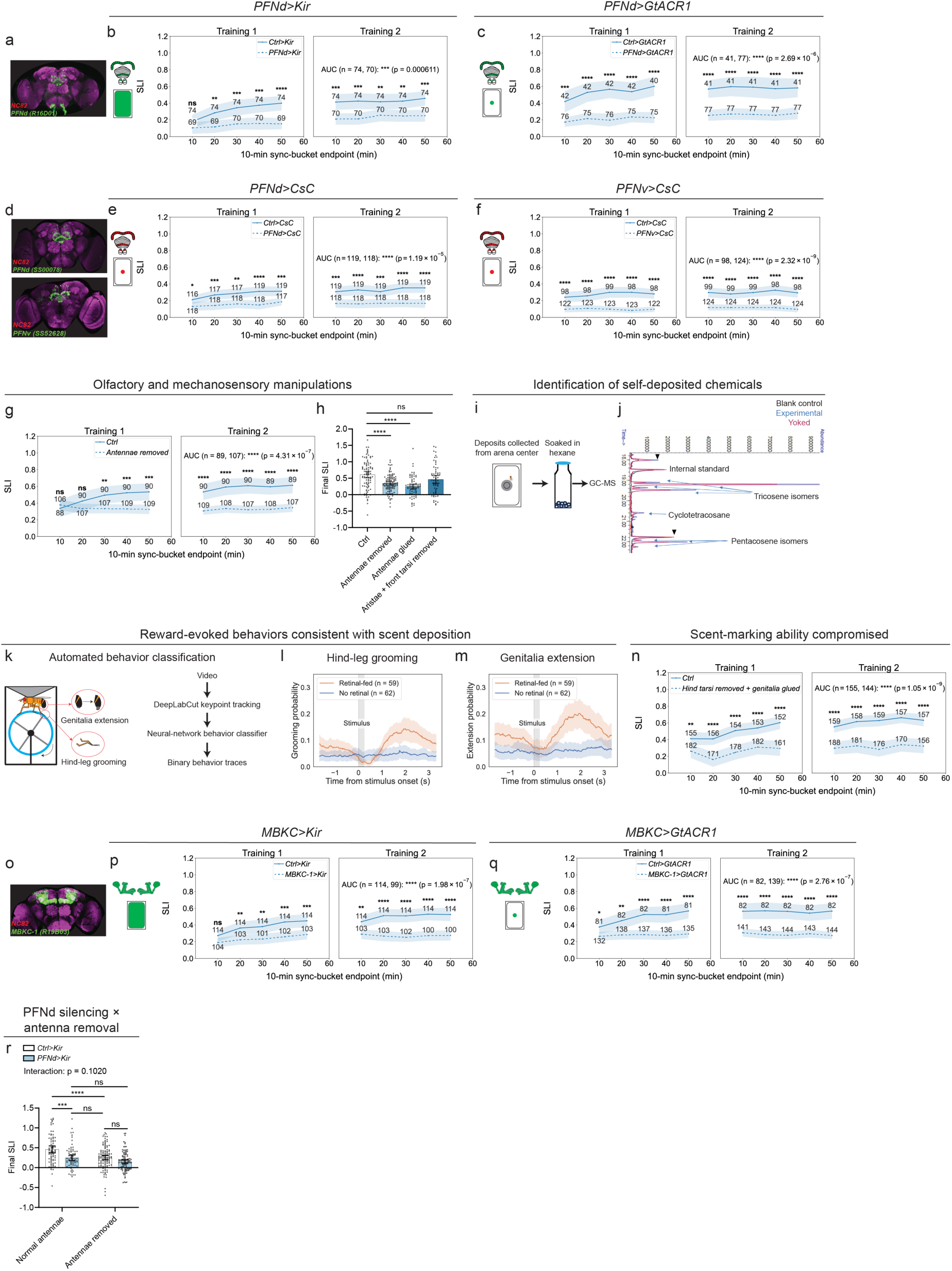
Flies rely on self-motion and olfactory cues to learn the spatial task. **a,** Expression pattern of the PFNd driver *R16D01-LexA*^3^. **b,c,** Spatial learning index (SLI) with PFNd silencing using Kir2.1 (Kir; **b**) or GtACR1 (**c**). Filled arena schematics indicate chronic manipulations; central discs indicate manipulations during reward delivery. **d,** Expression patterns of the split-GAL4 drivers for PFNd (SS00078; top) and PFNv (SS52628; bottom)^3,44^. **e,f,** SLI with PFNd (**e**) or PFNv (**f**) activated using CsChrimson (CsC). **g,** SLI after antenna removal; controls in **g,h** are from Fig. 1g. **h,** Final SLI (Training 2, sync bucket 5) of controls (*n* = 89) and flies with antennae removed (107), antennae glued (71), or aristae and front tarsi removed (61). **i,** Collection of arena-center deposits on transparent film (gray disc). **j,** Gas chromatography–mass spectrometry (GC–MS) chromatograms of experimental, yoked, and empty-arena control samples; putatively identified hydrocarbons are labeled. Triangles mark peaks in the blank control. **k,** Pose-based behavioral classification in thorax-fixed flies walking on a wheel. **l,m,** Stimulus-aligned probabilities of hind-leg grooming (**l**) and genitalia extension (**m**) in retinal-fed flies and no-retinal controls. Gray bars mark 250-ms light pulses used to activate 0273 neurons (onset, time 0). **n,** SLI with hind tarsi removed and genitalia glued. **o,** *R19B03-LexA*^45^ expression in mushroom body Kenyon cells (KCs). **p,q,** SLI with KC silencing using Kir (**p**) or GtACR1 (**q**). **r,** Final SLI for PFNd silencing × antenna removal (*n* = 74, 70, 110 and 106, left to right). Intact-antenna groups are from **b**. Behavioral experiments used the *0273>CsChrimson* background; controls for neuronal manipulations carried corresponding empty drivers (full genotypes, **Methods**). SLI time courses compare manipulations with corresponding controls during Training 1 and Training 2. Behavioral plots show means with 95% confidence intervals (shading or error bars). Points in **h,r** represent individual flies; numbers denote sample sizes. Two-sided Welch’s *t*-tests compare per-bucket and area-under-the-curve (AUC) values (**b,c,e–g,n,p,q**; **Methods**). Per-bucket *p* values are unadjusted. Welch’s ANOVA with Games-Howell comparisons (**h**); two-way ANOVA with Šídák’s comparisons (**r**). ns, not significant; \**p* < 0.05; \*\**p* < 0.01; \*\*\**p* < 0.001; \*\*\*\**p* < 0.0001.

We next tested whether EPG neurons, the “compass neurons,” may also play a role. EPG neurons represent another major class of spatially selective neurons whose population activities signal allocentric heading direction – the heading direction of a fly relative to a stable external landmark – and provide a known input to PFN neurons^2,3,39^. Interestingly, chronically silencing EPG neurons had no detectable effect on learning (**Extended Data Fig. 5a, b**). Similarly, neither transiently silencing nor activating EPG neurons during reward delivery had any detectable effect (**Extended Data Fig. 5c-e**). (As in the case of PFNs, activating the entire EPG population would be expected to disrupt its spatial selectivity.) Thus, we detected no requirement for the allocentric heading-direction signal represented by EPG neurons for flies to locate the goal in our task. Similarly, PFR activity, which tracks a signal closely related to allocentric translational direction^2^, also appeared dispensable (**Extended Data Fig. 6**).

Together, our results suggest that PFN-dependent egocentric self-motion cues support learning in our task, whereas we detected no requirement for EPG or PFR neurons, suggesting that the allocentric heading and translational signals represented by these populations may be dispensable under these conditions.

### Self-deposited odors also play a role in supporting spatial learning in the absence of prominent visual cues

Next, we asked whether flies could use sensory cues in addition to self-motion information to help solve our task, initially focusing on the antennae, which detect both mechanosensory and odor cues. We found that removing the antennae or covering them with UV-cured glue decreased learning significantly (**Fig. 2g, h**). Removing just the aristae – long hair-like structures that project from the tips of the antennae and are critical for detecting mechanosensory but not odor cues – together with the front tarsi, the mechanosensor-rich distal segments of the front legs, did not detectably impact learning (**Fig. 2h**). These results suggest that odors, rather than mechanosensory cues, might be the relevant antennae-detected signals supporting learning. To test more directly whether olfactory cues contributed to learning, we tested *Orco* mutants and found that SLI was reduced (**Extended Data Fig. 7**). The effect of *Orco* mutation on learning appeared smaller than that of antennae removal, possibly because *Orco*, an obligatory co-receptor for many odorant receptors, is expressed in only ∼78% of olfactory neurons^40^. The learning deficits of flies with impaired olfaction were, as before, unlikely to reflect altered reward seeking or walking speed (**Extended Data Fig. 8a, b, g, h**).

What then might be the source of the odors supporting learning? Given that the arenas were thoroughly cleaned prior to each experiment, we speculated that the putative odors might come from deposits made by the flies themselves at or around the goal. To test this, we first collected samples from the center of many arena floors after the flies had completed the task and sent the samples for gas chromatography-mass spectrometry (GC-MS) analysis (**Fig. 2i**, **Methods**). Indeed, several cuticular hydrocarbons, putatively identified as tricosene isomers, cyclotetracosane, and pentacosene isomers, were detected in these samples (**Table 1**, **Fig. 2j**); of note, tricosene and pentacosene are known fly pheromones^41,42^. Although we did not intentionally provide odor cues, these results indicate that the arenas were not necessarily odor-free during training. Next, using high-resolution recording combined with automated behavior classification, we found that optogenetic rewards increased hindleg grooming and genitalia extension, two behaviors that could facilitate transfer of scented material from the fly to the arena surface (**Fig. 2k-m**, **Methods**). Consistently, learning was reduced when we compromised flies’ capacity for scent transfer by removing their hind tarsi (the distal, segmented portion of the hind legs) and covering their genitalia with UV-cured glue (**Fig. 2n**). Next, the probability of making a sharp turn – a move that should bring flies closer to the goal – increased with increasing distance from the goal (**Extended Data Fig. 9a**). Importantly, this distance-dependent increase was reduced when we compromised flies’ olfaction or made them less capable of depositing scented material (**Extended Data Fig. 9b, c**), consistent with self-deposited olfactory cues contributing to this return behavior. Lastly, because previous work has shown that male *Drosophila* can deposit the cuticular hydrocarbon 9-tricosene in response to appetitive odors^24^, we tested *Or7a* mutants, which cannot detect this pheromone^24^. These mutants showed no detectable learning deficit, arguing against an essential role for 9-tricosene-Or7a signaling in our task (**Extended Data Fig. 23**).

We next assessed whether the self-deposited scents were strongly attractive intrinsically or became so through repeated association with reward. Several pieces of evidence suggest the latter. First, naïve flies showed no difference in occupancy near the goal between washed arenas and arenas that had just been used to train other flies, whereas trained flies spent significantly more time near the goal in the latter (**Extended Data Fig. 10**). Second, chronically silencing Kenyon cells (KCs), the odor-representing principal neurons of the mushroom body (MB), decreased learning significantly (**Fig. 2o, p**, **Extended Data Fig. 11a, b**). The MB is an olfactory association center in the fly brain where neutral odors represented by distinct KC ensembles can acquire positive or negative valence through association with reward or punishment^1,43^. Third, transiently silencing KCs at reward delivery also significantly decreased learning (**Fig. 2q**, **Extended Data Fig. 11c**), consistent with the notion that flies use KCs to associate scents from self-deposits with reward at the goal location. As in flies with impaired olfaction, the distance-dependent increase in the probability of making a sharp turn was reduced in KC-silenced flies (**Extended Data Fig. 9d, e**), and their learning deficits were unlikely to reflect altered reward seeking or walking speed (**Extended Data Fig. 8d-f, j-l**).

Lastly, we examined the joint contributions of PFN-dependent self-motion processing and olfactory input from self-deposited scents by assessing the impact of disrupting them separately versus together. Silencing PFNd neurons and removing the antennae each reduced SLI to similar levels (**Fig. 2r**). Combining the two manipulations lowered SLI only modestly further than either manipulation alone, a pattern that could reflect partly *interdependent* contributions of the two cue systems in guiding flies’ return to goal (**Fig. 2r**).

Together, these results suggest that, in feature-poor flat chambers, flies assign positive value to self-deposited scents near the goal and combine these learned olfactory cues with self-motion information to guide their return to an initially unmarked location.

### Flies are able to use external olfactory landmarks to improve learning

Given that flies appeared capable of assigning positive value to self-deposited scents associated with the goal and using them to help locate it, we wondered whether they could also use salient external odor landmarks *away* from the goal to improve performance. To address this, we constructed a set of apparatuses that allowed us to place strips of agarose in built-in troughs at two opposite edges of each arena, away from the goal (“agarose chamber”) (**Fig. 3a**, **Supplementary Video 2**). Flies performed significantly better in the agarose than in the flat chamber (**Fig. 3b, c**, **Extended Data Fig. 12a**). As in the flat chamber, Training 2 performance in the agarose chamber correlated with early Training 1 performance but not detectably with the extent of arena exploration during the Pre period (**Extended Data Fig. 12c, e**), and flies classified by their Training 2 performance as strong (top 20%) or weak (bottom 50%) learners showed an early and significant separation in SLI (**Extended Data Fig. 12g**).

**Fig. 3.**
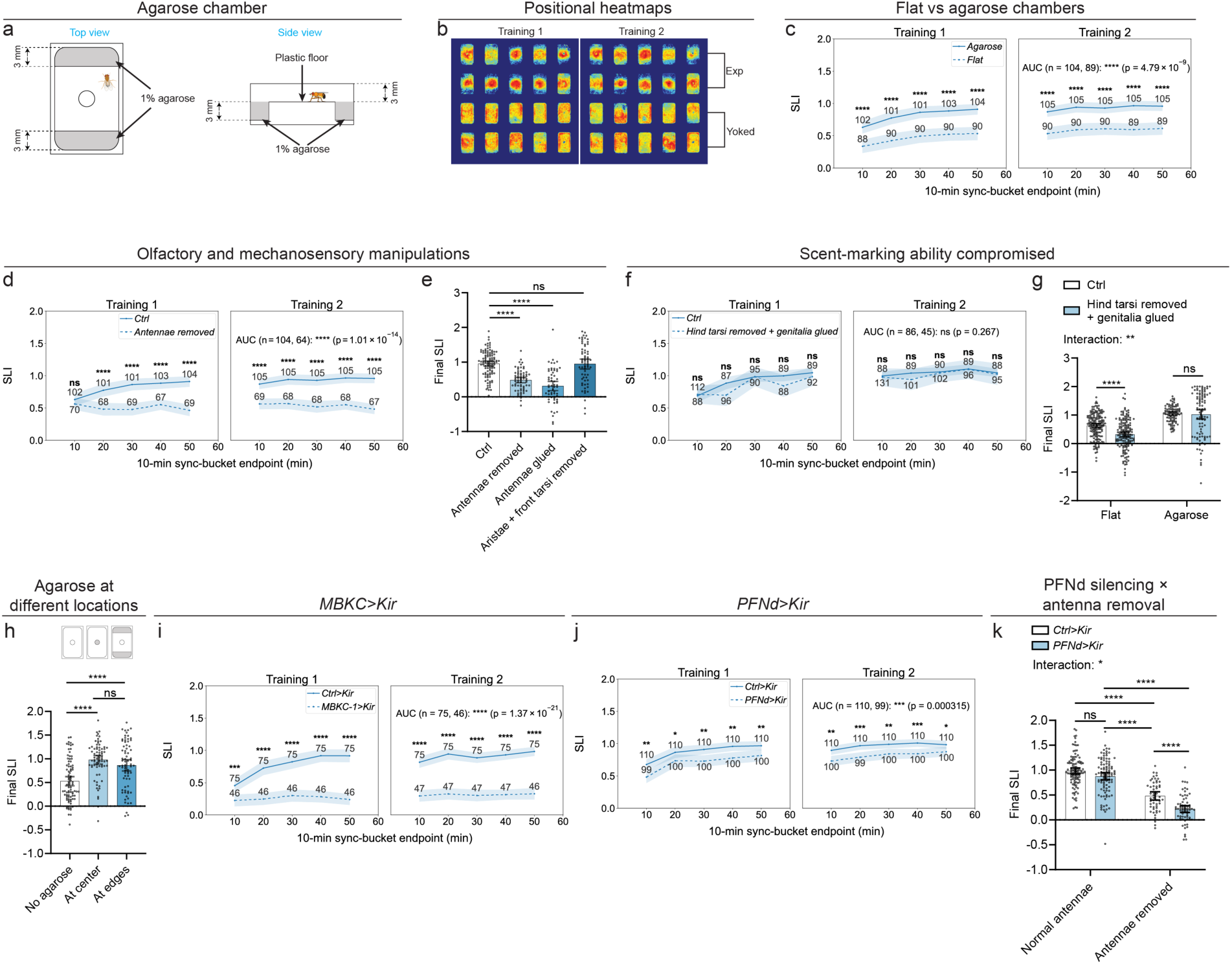
Flies use external olfactory landmarks to improve spatial learning. **a,** Schematic of the agarose chamber with two 1% agarose strips away from the goal. **b,** Positional heatmaps of 10 experimental–yoked pairs during Training 1 and Training 2. **c,** Spatial learning index (SLI; **Methods**) in flat versus agarose chambers; flat-chamber flies are from Fig. 1g. **d,** SLI after antenna removal. **e,** Final SLI (Training 2, sync bucket 5) of controls (*n* = 105) and flies with antennae removed (67), antennae glued (58), or aristae and front tarsi removed (63). Controls in **d,e** are the agarose group in **c**. **f,g,** Combined hind-tarsus removal and genitalia gluing: SLI during Training (**f**) and Final SLI in flat and agarose chambers (**g,** *n* = 157, 156, 88, 95, left to right). Flat-chamber groups are from Fig. 2n; agarose groups from **f**. **h,** Final SLI with no agarose (*n* = 85), agarose at the goal (73), or agarose at the edges (72). **i,j,** SLI with Kenyon cell (KC; **i**) or PFNd (**j**) silencing using Kir2.1 (Kir). **k,** Final SLI for PFNd silencing × antenna removal (*n* = 110, 100, 52, and 67, left to right); intact-antenna groups are from **j**. Assays used agarose chambers unless indicated otherwise. Flies carried *0273>CsChrimson* (full genotypes, **Methods**). SLI time courses compare conditions with corresponding controls in 10-min sync buckets during Training 1 and Training 2. Time courses and bars show means; shading and error bars indicate 95% confidence intervals. Points in **e,g,h,k** represent individual flies; numbers denote sample sizes. Two-sided Welch’s *t*-tests compare per-bucket and area-under-the-curve (AUC) values (**c,d,f,i,j**; **Methods**). Per-bucket *p* values are unadjusted. Welch’s ANOVA with Games-Howell comparisons (**e,h**); two-way ANOVA with Šídák’s comparisons (**g,k**). ns, not significant; \**p* < 0.05; \*\**p* < 0.01; \*\*\**p* < 0.001; \*\*\*\**p* < 0.0001.

Since agarose likely provided both olfactory and mechanosensory cues, we next assessed which type of cue flies primarily exploited. We found that removing both aristae and front tarsi, two mechanosensor-rich appendages that can contact agarose directly, did not detectably affect learning (**Fig. 3e**). In contrast, disrupting flies’ odor-sensing ability significantly reduced learning (**Fig. 3d, e**, **Extended Data Fig. 7**). These results suggest that flies primarily exploited olfactory rather than mechanosensory cues associated with the agarose to improve learning.

How might flies use the agarose-associated odors to improve learning? We first asked whether the benefit of agarose depended on use of self-deposited scents. We therefore compromised flies’ ability to deposit scents by simultaneously removing their hind tarsi and covering their genitalia with glue. Although this manipulation reduced learning in the flat chamber, it did not detectably reduce learning in the agarose chamber (**Fig. 3f, g**), indicating that flies can benefit from agarose even when their ability to deposit scents is compromised. We next asked whether flies avoided agarose innately or instead learned to treat agarose-associated odors as signals of *reward absence*. Several observations favored the latter interpretation. First, flies showed only very mild avoidance of agarose during the Pre period (**Extended Data Fig. 13**). Second, placing a small drop of agarose only at the reward location significantly improved learning relative to the flat-chamber condition (**Fig. 3h**), showing that agarose could also improve learning when directly associated with reward. Third, from the Pre period to the last sync bucket of Training 2, the percentage of time flies spent in agarose areas decreased more than the corresponding percentage in matched agarose-equivalent areas of the flat chamber (**Extended Data Fig. 13**). Finally, inhibiting KCs strongly compromised the agarose-induced improvement in performance (**Fig. 3i**, **Extended Data Fig. 11d-f**), consistent with a requirement for MB-dependent associative learning. Together, these findings favor the interpretation that flies learned to avoid agarose because it predicted reward absence, rather than simply strongly avoiding it innately. This interpretation is consistent with a previous finding that odors presented consistently *after* reward can acquire aversive value^46^, potentially because such post-reward odors signal “opportunity loss.”

We then asked whether self-motion cues remained important when external odor landmarks were available. Silencing PFNd neurons in the agarose chamber reduced learning, although the deficit was relatively mild (**Fig. 3j**). This deficit was substantially greater in olfaction-defective than in olfaction-intact flies (**Fig. 3k**), suggesting that self-motion cues become more important when olfactory input is compromised. These results suggest that self-motion and olfactory landmark cues both support learning in the agarose chamber, with each becoming more important when access to the other is reduced. Notably, despite their lower learning performance in the agarose chamber, PFNd-silenced flies with intact antennae showed an agarose-induced *improvement* in learning that did not differ significantly from that in controls (**Extended Data Fig. 12h**). By contrast, antenna removal markedly reduced the agarose-induced improvement observed in control flies (**Extended Data Fig. 12h**).

Together, our results from the agarose chamber show that flies can improve learning by exploiting external olfactory landmarks and support a model in which this improvement is mediated primarily by MB-dependent assignment of negative value to these landmarks. In this model, the landmarks serve as aversive guideposts that constrain search to a smaller region, within which self-deposited scents and self-motion information can guide navigation. The resulting reduction in spatial demand may explain why PFNd silencing produced only a modest learning deficit and why compromising scent *deposition* produced no detectable deficit.

### Flies exploit the same cue systems but adjust their reliance when task difficulty increases

The preceding experiments identified the cue systems that support learning in feature-poor and improvement in landmark-enriched chambers, but these arenas were relatively small (**Fig. 1a**). We next asked whether the same cue systems could support learning at a larger spatial scale and whether increasing arena size would alter flies’ relative reliance on self-deposited scents, self-motion information, and external olfactory landmarks. We therefore challenged flies to learn the reward location in a 39 x 39 mm arena (“large flat chamber”), with approximately 10 times the area of the small flat chamber (**Fig. 4a**). Although smaller than those used in previous local-search studies^19,20^, this arena provided more space for excursions while retaining sufficient throughput to systematically test multiple neuronal manipulations and their yoked controls. We retained the same temporal task structure used for the small chambers (**Fig. 1d**), but approximately doubled the reward-circle radius in Training 1 to increase the likelihood of initial reward encounters and reduced it to the small-chamber radius in Training 2 (**Fig. 4a**). Flies learned in the feature-poor large flat chamber, as indicated by their positional heatmaps, the shortening of their between-rewards trajectories over training, and positive SLI (**Fig. 4b-d**). The reduction in SLI from Training 1 to Training 2 reflects the smaller reward circle, which was more difficult to find. As in the small flat chamber, SLI correlated well with yoked-subtracted RPD (**Extended Data Fig. 14a-c**), early performance predicted later performance (**Extended Data Fig. 14f**), and strong and weak learners separated early (**Extended Data Fig. 14h**). One difference emerged with increasing arena size: whereas Pre-period exploration was not detectably associated with subsequent learning in the small flat chamber, it showed a modest positive correlation with SLI in the large flat chamber (**Extended Data Figs. 3b, 14d**), suggesting that prior exploration may become more useful as spatial scale increases.

**Fig. 4.**
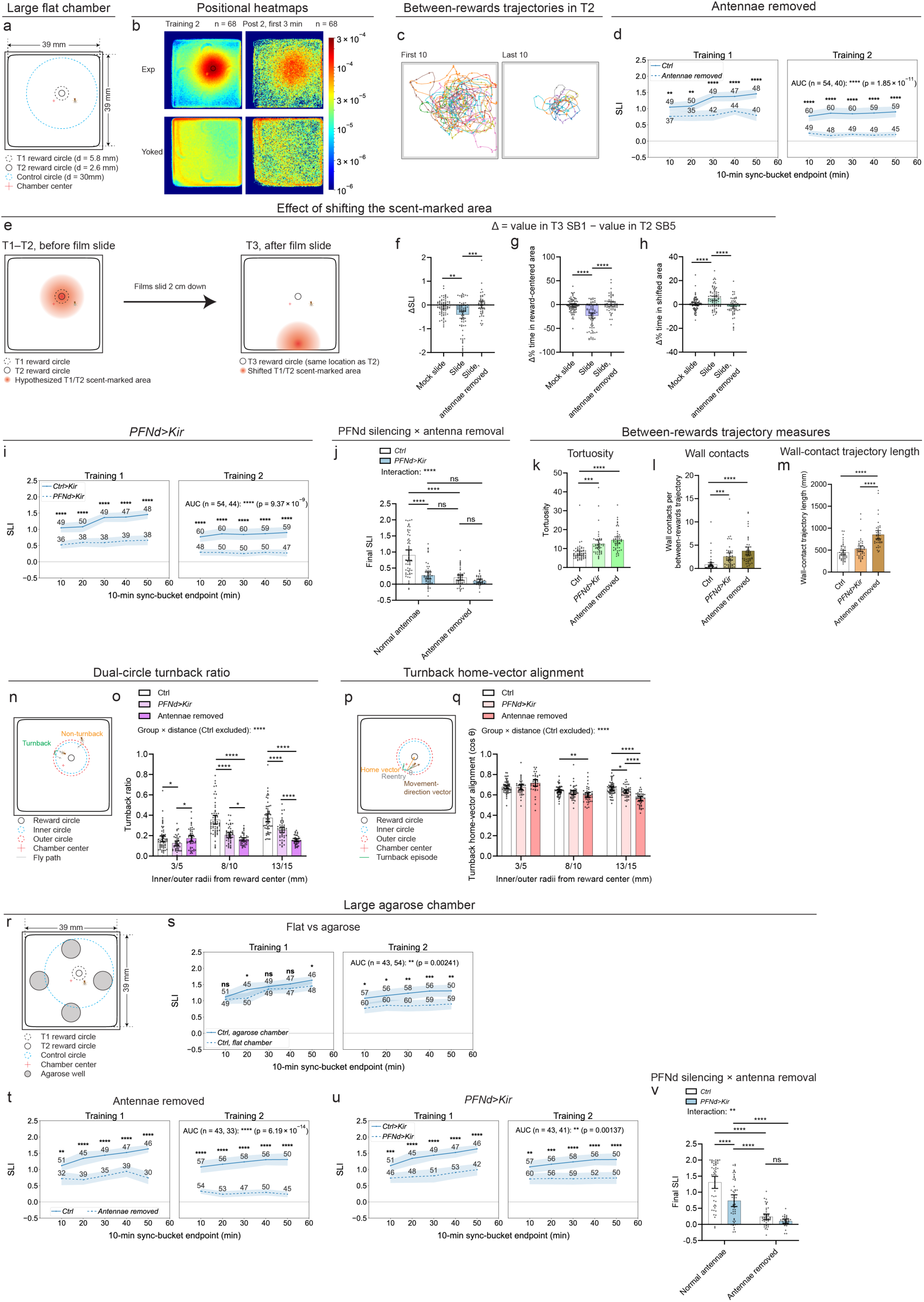
Flies use self-motion and olfactory cues for spatial learning in larger arenas. **a,** Schematic of the large flat chamber (39 × 39 mm), with different reward-circle sizes in Training 1 (T1) and Training 2 (T2). **b,** Mean positional heatmaps of experimental and yoked flies during T2 and the first 3 min of Post 2. **c,** First (left) and last (right) 10 between-rewards trajectories of one fly in T2. **d,** Spatial learning index (SLI) after antenna removal. **e,** Films lining the chamber were shifted 20 mm after T2, before Training 3 (T3), without moving the reward location; mock-slide films were returned to their original positions. **f-h,** Change (T3 SB1 − T2 SB5; SB, sync bucket) in SLI (**f**) and percentage of time in reward-centered (**g**) or shifted (**h**) areas (diameter, 20 mm). Group order: mock-slide, slide, antennae-removed slide; *n* = 71, 70, 45 (**f**) and 83, 78, 57 (**g,h**). **i,** SLI with PFNd silencing using Kir2.1 (Kir). **j,** Final SLI (T2 SB5) for PFNd silencing × antenna removal (*n* = 59, 47, 45, 33, left to right). **k–m,** Tortuosity (**k**), wall contacts per between-rewards trajectory (**l**), and wall-contact trajectory length (**m**). Group order in **k–m,o,q**: Ctrl, *PFNd>Kir*, antennae removed; *n* = 58, 46, 45 (**k,l**) and 50, 46, 42 (**m**). **n,o,** Dual-circle turnback schematic (**n**) and ratios (**o**): fraction of excursions beyond the inner circle that return before crossing the outer circle. Inner/outer radii from reward center: 3/5, 8/10, 13/15 mm; *n* = 59, 46, 45; 59, 47, 45; 58, 47, 45, respectively. **p,q,** Turnback home-vector alignment: schematic (**p**) and cos θ (**q**), where θ is the angle between movement direction and the vector toward the reward center at successful reentry; *n* = 55, 38, 36; 58, 45, 45; 57, 46, 45 for the same radius pairs. Measures in **k–m,o,q** use T2 SB2–5 from the corresponding cohorts in **j** (**Methods**). **r,** Large agarose chamber with four off-goal agarose patches (diameter, 8 mm); other dimensions as in **a**. **s,** SLI in flat versus agarose chambers. **t,u,** SLI after antenna removal (**t**) or PFNd silencing (**u**) in agarose chambers. **v,** Final SLI for PFNd silencing × antenna removal in agarose chambers (*n* = 50, 50, 45, 29, left to right). Controls are shared across **d,i,s** (flat) and **s–u** (agarose); **j,v** reuse the corresponding control and single-manipulation groups. Assays used large flat chambers unless indicated otherwise. Flies carried *0273>CsChrimson* (full genotypes, **Methods**). SLI time courses show T1 and T2 in 10-min sync buckets. Time courses and bars show means; shading and error bars indicate 95% confidence intervals. Points in bar plots represent individual flies; numbers denote sample sizes. Two-sided Welch’s *t*-tests compare per-bucket and area-under-the-curve (AUC) values (**d,i,s–u**; **Methods**); per-bucket *p* values are unadjusted. Welch’s ANOVA with Games-Howell comparisons (**f-h,k-m**); two-way ANOVA with Šídák comparisons (**j,v**). Mixed-effects models account for repeated distances within flies, with Tukey comparisons among groups at each distance (**o,q**); displayed group × distance interactions exclude controls. ns, not significant; \**p* < 0.05; \*\**p* < 0.01; \*\*\**p* < 0.001; \*\*\*\**p* < 0.0001.

We found that flies still relied on olfaction for learning in the large flat chamber, as antennae-removed flies showed strongly reduced SLI (**Fig. 4d, j**, **Extended Data Fig. 15d**). The larger arena also allowed more detailed analysis of between-rewards trajectories with substantially less influence from the sidewalls. We therefore quantified several features of these trajectories using measures inspired by previous work^19,20,47^. Compromising olfaction shifted the center of mass (COM) of the trajectories farther from the reward center (**Extended Data Fig. 15b, e**), lengthened the return leg from the trajectory’s farthest point back to the reward (**Extended Data Fig. 15c, f**), and increased trajectory tortuosity (**Fig. 4k**). Together, these changes indicate that impaired olfaction made returns to the goal less spatially focused and less efficient.

To test more directly whether reward-associated self-deposited cues contributed to learning in the large flat chamber, we adapted a substrate-shifting approach from previous work^20,48^. We incorporated transparent sliding films into the chamber. After Training 2, we slid the films by 2 cm while leaving the software-defined reward location unchanged, thereby displacing material deposited near the reward location during Trainings 1 and 2 before the start of Training 3 (**Fig. 4e**, **Methods**). Immediately after the slide, SLI decreased relative to the end of Training 2, and flies spent less time in the reward-centered area and more time in the shifted area (**Fig. 4f–h**). Neither mock-slide control flies, whose films were returned to their original positions before Training 3, nor antennae-removed flies showed a comparable behavioral shift (**Fig. 4f–h**), indicating that the shift depended on substrate displacement and intact olfaction. As in the small flat chamber, silencing Kenyon cells in the mushroom body also reduced learning and altered between-rewards trajectories in the large chamber (**Extended Data Figs. 14i, 15j-l**).

As in the small flat chamber, silencing EPG neurons had no detectable effect on performance in the large flat chamber, whereas PFN-dependent self-motion processing remained important (**Fig. 4i**, **Extended Data Fig. 14j**). PFNd silencing reduced SLI to a similar extent as antenna removal (**Fig. 4j**). It also increased trajectory tortuosity (**Fig. 4k**), shifted the COM of between-rewards trajectories farther from the reward center, and lengthened their return legs (**Extended Data Fig. 15g–i**). Further, the two manipulations showed a significant interaction: PFNd silencing strongly reduced SLI in flies with intact antennae but had little additional effect after antenna removal (**Fig. 4j**). This non-additive pattern is again consistent with partly *interdependent* contributions of olfactory and self-motion information to guiding returns to the goal.

Despite their similar effects on overall learning (**Fig. 4d, i-l**), antenna removal and PFNd silencing produced different trajectory phenotypes. For example, antennae-removed flies had longer wall-contact trajectory lengths than PFNd-silenced flies (**Fig. 4m**), suggesting that the relative contributions of olfaction and self-motion might vary with distance from the goal. To test this possibility, we quantified the dual-circle turnback ratio: the fraction of outward excursions beyond an inner radius in which the fly returned across that radius before reaching a larger outer radius (**Fig. 4n**, **Methods**). The effects of antenna removal and PFNd silencing differed across distances (**Fig. 4o**). At the farthest tested radius pair (13/15 mm), antenna removal reduced the turnback ratio more than PFNd silencing; at the nearest radius pair (3/5 mm), PFNd silencing instead reduced the turnback ratio more than antenna removal (**Fig. 4o**). For successful turnbacks, we also measured how closely the fly’s direction of movement at reentry aligned with the home vector toward the reward center (**Fig. 4p**). Here, antenna removal did not detectably alter alignment at 3/5 mm but reduced it at 8/10 and 13/15 mm, with the strongest reduction at the farthest tested distance (**Fig. 4q**). Together, these analyses reveal distance-dependent differences in how flies organize and direct their returns. In particular, the results suggest that PFNd-dependent self-motion information contributes relatively more to local returns, whereas olfactory cues become relatively more important during excursions farther from the goal.

Next, we asked whether, at a larger spatial scale, flies could still use external odor landmarks to improve learning. We placed four agarose patches away from the goal to create a “large agarose chamber” (**Fig. 4r**, **Supplementary Video 3**) and found that flies learned better than in the large flat chamber (**Fig. 4s**). As in the small agarose chamber, removing the antennae, silencing Kenyon cells in the mushroom body, or silencing PFNd neurons reduced learning, whereas silencing EPG neurons had no detectable effect; antenna removal produced a larger deficit than PFNd silencing (**Fig. 4t-v**, **Extended Data Fig. 16i, j**). To test whether flies learned to avoid the agarose landmarks, we quantified the agarose avoidance ratio: the fraction of approaches entering a 5-mm-radius zone around an agarose patch that ended without the fly’s entering the 4-mm-radius agarose region (**Extended Data Fig. 17a**, **Methods**). During training, avoidance increased in the large agarose chamber relative to corresponding virtual regions in the large flat chamber (**Extended Data Fig. 17b**). This differential avoidance was lost after antenna removal or Kenyon-cell silencing but persisted after PFNd or EPG silencing (**Extended Data Fig. 17c-f**), supporting the idea that flies learned through an olfactory, MB-dependent mechanism to treat agarose as an aversive landmark.

Finally, we asked whether increasing spatial scale altered flies’ reliance on self-motion cues when external olfactory landmarks were available. PFNd-silenced flies learned less well than controls at both scales (**Figs. 3j and 4u**, **v**), supporting a contribution of PFNd neurons to learning in landmark-enriched arenas. However, following antenna removal, PFNd silencing further reduced SLI in the small agarose chamber but produced no detectable additional deficit in the large agarose chamber (**Figs. 3k and 4v**). Thus, a contribution of PFNd neurons in antennae-removed flies was evident in the small arena but could not be detected in the large arena. Notably, despite their lower learning performance, PFNd-silenced flies with intact antennae retained substantial agarose-induced *improvement* at both scales, whereas antenna removal markedly reduced this improvement in control flies (**Extended Data Figs. 12h and 16k**).

Together, these results show that the cue systems supporting learning in the small chambers continue to operate at a larger spatial scale. In the feature-poor large flat chamber, MB-dependent use of self-deposited scents and PFN-dependent self-motion information jointly support returns to the goal, with partly interdependent contributions. The larger arena further suggested a distance-dependent division of labor: PFN-dependent self-motion information contributed relatively more to local returns, whereas olfactory cues became relatively more important during farther excursions. In the landmark-enriched large agarose chamber, flies again improved performance by using MB-dependent olfactory learning to treat agarose as an aversive landmark. After antenna removal, however, the pattern differed between chamber sizes: while PFNd silencing further reduced SLI in the small agarose chamber, it produced no detectable additional deficit in the large agarose chamber. This suggests that PFN-dependent self-motion information may have lower capacity to compensate for impaired olfaction at the larger spatial scale.

### Feature-poor learning and landmark-dependent improvement are under genetic control and show shared and distinct transcriptional signatures

Finally, we leveraged the throughput of our small-chamber assay to investigate how genetic background affects flies’ ability to learn and to explore molecular signatures associated with learning. We first asked whether the capacities to learn in feature-poor environments and to benefit from external olfactory landmarks vary with genetic background, and whether these capacities are partly separable. To test this, we assessed the F_1_ progeny of crosses between females from 32 wild-derived *Drosophila* lines and *w*¹¹¹⁸ males carrying optogenetic reinforcement transgenes (**Fig. 5a**). Despite sharing the same paternal genetic background and being raised and tested under the same conditions, progeny of different maternal lines differed significantly in learning performance in both flat and agarose chambers (**Fig. 5a**). Importantly, we also found a significant line × chamber interaction, indicating that the benefit from agarose varied among lines (**Fig. 5a**). For example, progeny of Line 7 (Taiwan) learned poorly in the flat chamber but performed similarly to the standard laboratory strain in the agarose chamber. In contrast, progeny of Line 45 (India) learned well in the flat chamber but showed no detectable improvement with agarose. Together, these results support a strong genetic contribution to spatial learning ability and suggest that learning in feature-poor environments and exploiting external olfactory landmarks are under partly distinct genetic influences.

**Fig. 5.**
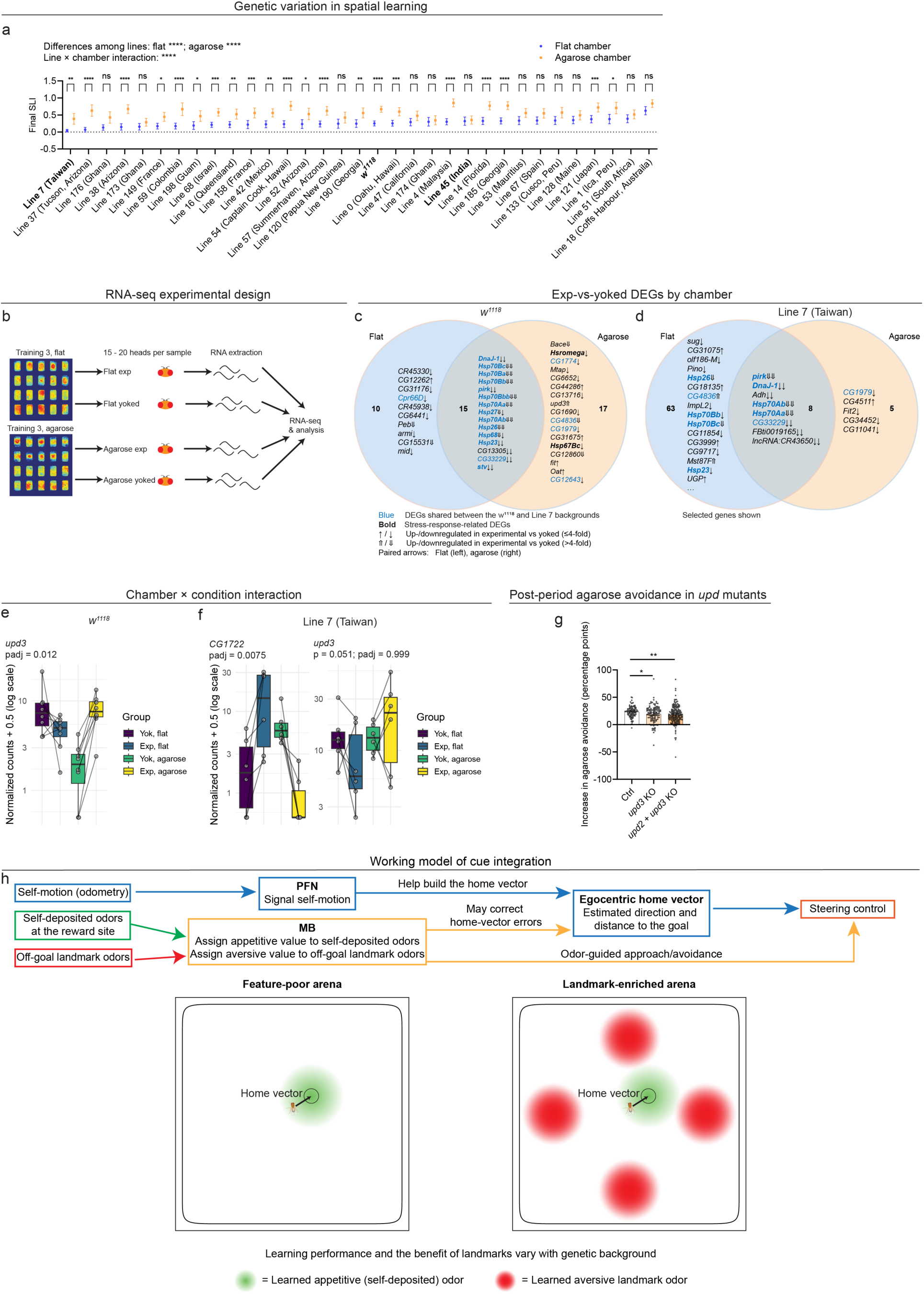
Genetic and molecular evidence for flexible spatial learning strategies depending on landmark availability. **a,** Final spatial learning index (SLI; Training 2, sync bucket 5) in flat and agarose chambers for male F_1_ progeny of females from 32 wild-derived lines or *w*¹¹¹⁸ crossed with males carrying reinforcement transgenes (**Methods**). Sample sizes (flat/agarose): Line 7, 56/43; 37, 45/47; 176, 43/39; 38, 51/58; 173, 35/42; 149, 67/67; 59, 39/43; 198, 65/49; 68, 65/61; 16, 55/59; 158, 59/64; 42, 50/56; 54, 57/47; 52, 58/42; 57, 69/58; 120, 55/46; 190, 58/58; *w*¹¹¹⁸, 92/109; 0, 67/63; 47, 56/59; 174, 65/44; 4, 70/61; 45, 54/53; 14, 63/63; 185, 70/66; 53, 56/57; 67, 61/54; 133, 52/58; 128, 54/60; 121, 67/58; 1, 46/47; 51, 60/59; 18, 57/71. **b,** RNA-seq design: 15–20 heads per sample, collected after Training 3; eight experimental–yoked sample pairs per chamber for *w*¹¹¹⁸ and six for Line 7. **c,d,** Experimental-versus-yoked differentially expressed genes (DEGs; adjusted *p* < 0.05) in flat and agarose chambers for *w*¹¹¹⁸ (**c**) and Line 7 (**d**). Selected flat-only genes are shown in **d**; complete lists, **Tables 2–5**. Within each region, genes are ordered by adjusted *p* value (most significant first), using flat-chamber values for genes shared between chambers. **e,f,** DESeq2-normalized counts + 0.5 (log scale) of *upd3* in *w*¹¹¹⁸ (**e**), and *CG1722* and *upd3* in Line 7 (**f**). Points represent pooled-head samples; lines join experimental–yoked pairs. Boxes show medians and interquartile ranges; whiskers extend to the most extreme values within 1.5 times the interquartile range of the box edges. Boxplot statistics are calculated on the log₁₀ scale. **g,** Increase in agarose avoidance from Pre to Post 3 (last 10 min of each period) in controls (*n* = 93), *upd3* knockout (KO; 113), and *upd2* + *upd3* double KO flies (176). **h,** Working model of joint guidance by self-motion and learned olfactory cues. Green, self-deposited odors with learned appetitive value; red, off-goal landmark odors with learned aversive value; MB, mushroom body. In **a,g**, means and 95% confidence intervals are shown; points in **g** represent individual flies. Welch’s ANOVA compares lines within each chamber; two-way ANOVA tests line × chamber interaction, with Šídák comparisons between chambers for each line (**a**). DESeq2 Wald tests for chamber × condition interactions, with Benjamini–Hochberg-adjusted *p* values (**e,f**; **Methods**); two-sided Welch’s *t*-tests with Holm–Šídák correction for the two mutant-versus-control comparisons (**g**). ns, not significant; \**p* < 0.05; \*\**p* < 0.01; \*\*\**p* < 0.001; \*\*\*\**p* < 0.0001.

**Table 3.** DEGs of exp vs. yoked flies in the agarose chamber, *w^1118^* background.

| **Gene** | **padj** | **pvalue** | **log2FoldChange** | **baseMean** |
|---|---|---|---|---|
| *DnaJ-1* | 1.76446E-11 | 1.5594E-15 | -0.945090831 | 6682.89538 |
| *pirk* | 1.01295E-08 | 1.79045E-12 | -1.765963853 | 622.4804769 |
| *Hsp70Bc* | 5.77612E-06 | 1.53145E-09 | -2.962490753 | 350.6932071 |
| *Hsp26* | 1.2424E-05 | 4.39206E-09 | -2.545200658 | 1159.297935 |
| *Hsp70Bb* | 5.77545E-05 | 2.55212E-08 | -2.767395385 | 1279.999904 |
| *Bace* | 7.8624E-05 | 4.16919E-08 | -6.699536933 | 113.1478825 |
| *Hsp23* | 8.67172E-05 | 6.89752E-08 | -1.128550613 | 662.2332408 |
| *Hsp70Bbb* | 8.67172E-05 | 6.48185E-08 | -2.828926881 | 602.9822009 |
| *Hsp70Ba* | 8.67172E-05 | 5.99623E-08 | -2.843582605 | 200.3316547 |
| *Hsp70Aa* | 0.000100131 | 8.84944E-08 | -3.002471695 | 462.6835101 |
| *Hsp70Ab* | 0.000202409 | 1.96774E-07 | -2.893242022 | 1490.372849 |
| *Hsp27* | 0.000341506 | 3.6218E-07 | -1.641415892 | 374.429149 |
| *Hsromega* | 0.000749524 | 9.27382E-07 | -1.089176587 | 5752.578233 |
| *CG13305* | 0.000749524 | 9.16302E-07 | -0.629984693 | 141.5542801 |
| *Hsp68* | 0.00097901 | 1.29785E-06 | -1.588531248 | 2437.230876 |
| *CG1774* | 0.001931848 | 2.89165E-06 | -0.349644188 | 1006.281782 |
| *Mtap* | 0.001931848 | 2.90247E-06 | -0.406799711 | 869.5789162 |
| *CG6652* | 0.002640433 | 4.20042E-06 | -0.76395653 | 33.00803686 |
| *CG44286* | 0.003164224 | 5.31332E-06 | 0.756613612 | 37.38348626 |
| *CG13716* | 0.005944023 | 1.05064E-05 | -1.937425464 | 5.826944357 |
| *stv* | 0.006077082 | 1.12787E-05 | -0.518123666 | 4691.085418 |
| *upd3* | 0.009225113 | 1.79366E-05 | 2.289811894 | 5.329893188 |
| *CG33229* | 0.012459881 | 2.53272E-05 | -0.323119753 | 1272.425727 |
| *CG1690* | 0.014660271 | 3.10956E-05 | -0.773050017 | 87.57476181 |
| *CG4836* | 0.019342334 | 4.2736E-05 | -3.150167407 | 44.67969917 |
| *CG1979* | 0.023244875 | 5.34129E-05 | -0.453798991 | 839.7916888 |
| *CG31675* | 0.023438268 | 5.59287E-05 | 0.368706446 | 95.93719538 |
| *Hsp67Bc* | 0.024171662 | 5.9815E-05 | -0.549190698 | 107.1898748 |
| *CG12860* | 0.046332279 | 0.000118748 | -4.905549597 | 9.174980313 |
| *fit* | 0.046852705 | 0.000124223 | 0.809225312 | 280.2422355 |
| *Oat* | 0.047762998 | 0.000134105 | 0.239130579 | 1467.021575 |
| *CG12643* | 0.047762998 | 0.000135079 | -0.524138918 | 406.2134448 |

**Table 4.** DEGs of exp vs. yoked flies in the flat chamber, Line 7 (Taiwan) background.

| Gene | padj | pvalue | log2FoldChange | baseMean |
|---|---|---|---|---|
| *pirk* | 9.44071E-22 | 6.95E-26 | -3.17007 | 550.7845 |
| *DnaJ-1* | 7.56584E-09 | 1.78E-12 | -0.81948 | 9580.06 |
| *Adh* | 7.56584E-09 | 2.23E-12 | -0.45666 | 747.9772 |
| *Hsp70Ab* | 7.56584E-09 | 1.65E-12 | -4.13846 | 600.2862 |
| *sug* | 2.15082E-08 | 7.92E-12 | -1.41241 | 1060.596 |
| *Hsp70Aa* | 7.41006E-08 | 3.27E-11 | -3.95545 | 217.4752 |
| *CG31075* | 1.93805E-06 | 9.99E-10 | 0.479182 | 6982.644 |
| *olf186-M* | 1.42101E-05 | 8.37E-09 | -0.41861 | 1091.948 |
| *Pino* | 4.17471E-05 | 2.77E-08 | -0.37209 | 10128.79 |
| *Hsp26* | 4.2621E-05 | 3.14E-08 | -2.24437 | 1038.762 |
| *CG18135* | 0.000112703 | 9.13E-08 | 0.370595 | 18327.06 |
| *CG4836* | 0.000272976 | 2.41E-07 | 3.717254 | 67.80728 |
| *ImpL2* | 0.000584758 | 5.82E-07 | -0.36497 | 1308.926 |
| *Hsp70Bb* | 0.000584758 | 6.46E-07 | -1.22115 | 317.7202 |
| *Hsp70Bc* | 0.000584758 | 6.32E-07 | -2.08324 | 148.4164 |
| *CG11854* | 0.000661828 | 7.79E-07 | -0.72749 | 130.1304 |
| *CG3999* | 0.000729071 | 9.12E-07 | 0.728106 | 876.1658 |
| *CG33229* | 0.000733199 | 9.71E-07 | -0.35083 | 2472.022 |
| *CG9717* | 0.001117098 | 1.56E-06 | -0.32014 | 1504.963 |
| *Mst87F* | 0.001503491 | 2.21E-06 | 3.275213 | 11.65601 |
| *Hsp23* | 0.002724835 | 4.21E-06 | -1.0638 | 1006.365 |
| *UGP* | 0.004193023 | 6.79E-06 | 0.276202 | 11931.2 |
| *CG14718* | 0.005137651 | 8.7E-06 | 3.483017 | 10.59746 |
| *FBti0019165* | 0.005688525 | 1.03E-05 | -0.69403 | 146.5141 |
| *CG42564* | 0.005688525 | 1.05E-05 | 0.910375 | 62.2713 |
| *Hsp68* | 0.006110748 | 1.2E-05 | -1.40498 | 1915.734 |
| *CG2861* | 0.006110748 | 1.21E-05 | 2.892651 | 42.73441 |
| *Sirup* | 0.006852472 | 1.46E-05 | 0.296954 | 6543.626 |
| *CG12643* | 0.006852472 | 1.45E-05 | -0.37926 | 464.4633 |
| *Lsd-1* | 0.006999862 | 1.55E-05 | 0.287078 | 8629.292 |
| *Hsp70Bbb* | 0.007637185 | 1.74E-05 | -2.73142 | 55.85069 |
| *Pen* | 0.010078543 | 2.37E-05 | 3.010233 | 52.79438 |
| *Got2* | 0.011511503 | 2.97E-05 | 0.264123 | 23380.76 |
| *Hsp70Ba* | 0.011511503 | 2.89E-05 | -2.15153 | 44.80768 |
| *CG32548* | 0.011511503 | 2.95E-05 | 1.508373 | 36.15083 |
| *lncRNA:CR46006* | 0.013944154 | 3.7E-05 | -0.48985 | 277.9012 |
| *stv* | 0.015048479 | 4.1E-05 | -0.33681 | 6796.087 |
| *CG2065* | 0.015815534 | 4.42E-05 | -0.47262 | 416.7839 |
| *TrxT* | 0.0170883 | 4.91E-05 | 2.615509 | 25.65211 |
| *Galk* | 0.017139486 | 5.08E-05 | 0.310231 | 12773.81 |
| *lncRNA:CR43650* | 0.017139486 | 5.17E-05 | -0.74153 | 516.9493 |
| *CG7886* | 0.018202881 | 5.63E-05 | 3.237613 | 30.12014 |
| *SLC5A11* | 0.018332659 | 5.8E-05 | 0.24403 | 2242.847 |
| *h* | 0.021723704 | 7.04E-05 | -0.34027 | 2127.366 |
| *Caf1-180* | 0.023307365 | 7.72E-05 | 0.430281 | 249.9539 |
| *CG15109* | 0.023902863 | 8.09E-05 | 3.927603 | 15.93088 |
| *CG16782* | 0.024416176 | 8.45E-05 | 3.204454 | 10.78587 |
| *lectin-28C* | 0.025337176 | 8.95E-05 | 0.398242 | 1638.007 |
| *CG3330* | 0.026058396 | 9.4E-05 | 4.624395 | 11.52126 |
| *Hsp83* | 0.027431508 | 0.000101 | -0.40312 | 50730.14 |
| *CG10859* | 0.02808696 | 0.000105 | 6.911894 | 10.74627 |
| *SkpB* | 0.035184039 | 0.000135 | 1.271102 | 27.1962 |
| *dsx* | 0.036028641 | 0.000141 | -0.21567 | 1433.027 |
| *Cyp4d21* | 0.037117153 | 0.000148 | -0.27476 | 7080.196 |
| *Hsp60B* | 0.038397359 | 0.000155 | 4.145023 | 13.60788 |
| *hll* | 0.038439197 | 0.00016 | 0.816605 | 659.2921 |
| *CG32591* | 0.038439197 | 0.000161 | 1.072695 | 63.98575 |
| *CG1722* | 0.038468739 | 0.000164 | 2.981386 | 6.245153 |
| *Hsp27* | 0.039366411 | 0.000173 | -1.23944 | 384.1476 |
| *Cpr66D* | 0.039366411 | 0.000174 | -0.82898 | 140.2097 |
| *xmas* | 0.040938701 | 0.000184 | 0.233757 | 795.3803 |
| *CG1774* | 0.043310181 | 0.000198 | -0.37066 | 1346.436 |
| *CG30357* | 0.044346608 | 0.000207 | -0.50281 | 211.5034 |
| *CG32119* | 0.044346608 | 0.000209 | 5.339671 | 10.30743 |
| *Mtpalpha* | 0.044369252 | 0.000213 | 0.174876 | 12619.7 |
| *foxo* | 0.044369252 | 0.000216 | -0.14483 | 3839.98 |
| *SPH93* | 0.044369252 | 0.000219 | -5.1167 | 252.852 |
| *CG10512* | 0.04477271 | 0.000231 | -0.25358 | 2528.157 |
| *CG5089* | 0.04477271 | 0.000226 | 3.281889 | 19.03742 |
| *FBti0018978* | 0.04477271 | 0.000231 | 1.443573 | 16.81991 |
| *lncRNA:CR32661* | 0.048524063 | 0.000254 | 2.747884 | 11.09442 |

**Table 5.** DEGs of exp vs. yoked flies in the agarose chamber, Line 7 (Taiwan) background.

| **Gene** | **padj** | **pvalue** | **log2FoldChange** | **baseMean** |
|---|---|---|---|---|
| *pirk* | 2.21687E-12 | 2.02E-16 | -2.48998 | 550.7845 |
| *Adh* | 4.75467E-08 | 8.68E-12 | -0.4585 | 747.9772 |
| *DnaJ-1* | 3.03997E-06 | 8.33E-10 | -0.71408 | 9580.06 |
| *CG1979* | 3.86082E-05 | 1.41E-08 | -0.86245 | 1029.236 |
| *lncRNA:CR43650* | 0.00516889 | 2.36E-06 | -0.86398 | 516.9493 |
| *CG33229* | 0.011304095 | 6.19E-06 | -0.32544 | 2472.022 |
| *Hsp70Aa* | 0.012071761 | 7.72E-06 | -2.68543 | 217.4752 |
| *CG4511* | 0.016590943 | 1.21E-05 | 0.233771 | 1501.898 |
| *Hsp70Ab* | 0.021808512 | 1.79E-05 | -2.51681 | 600.2862 |
| *FBti0019165* | 0.023877728 | 2.18E-05 | -0.68935 | 146.5141 |
| *Fit2* | 0.034640598 | 3.52E-05 | -0.34772 | 370.27 |
| *CG34452* | 0.034640598 | 3.8E-05 | -1.19309 | 50.77851 |
| *CG11041* | 0.048179265 | 5.72E-05 | -0.92013 | 62.5982 |

To explore whether learning in flat and agarose chambers was associated with distinct transcriptional responses, we asked whether gene-expression differences between experimental flies and their yoked controls varied with chamber type. We performed bulk RNA-seq on heads collected immediately after the task from experimental and yoked flies in both chambers, using two genetic backgrounds, *w^11^*^18^ and Line 7 (Taiwan) (**Fig. 5b**, **Methods**). A key advantage of our yoked-control design is that each yoked fly received the exact same optogenetic rewards at the same times as its paired experimental fly, but independently of its position. This matching largely controls for transcriptional responses to reward stimulation itself, potentially providing high specificity for changes associated with the learnable spatial contingency. Such differences need not reflect learning itself, however; they could also arise from responses to the absence of a learnable contingency in yoked flies or other behavioral and physiological consequences of the two conditions.

Several findings emerged from our RNA-seq analyses of *w^1118^*and Line 7. First, the experimental-versus-yoked comparisons identified relatively few differentially expressed genes (DEGs) in either chamber, consistent with the potential specificity afforded by our yoked-control design (**Tables 2–5**, **Fig. 5c, d**, **Extended Data Fig. 18-21**). Second, a striking pattern emerged: many of the most statistically significant DEGs were stress-associated and more highly expressed in yoked than in experimental flies. Indeed, all DEGs shared between the flat and agarose chambers showed this direction of expression change, and 14 of the 15 shared DEGs in *w^1118^* were annotated as stress-associated (**Fig. 5c, d**, **Extended Data Fig. 18**). Third, chamber × condition interaction tests identified genes for which the experimental-versus-yoked expression difference depended on chamber type, including *upd3* in *w^1118^* and *CG1722* in Line 7 (**Fig. 5e, f**). The identification of *upd3*, which encodes a cytokine ligand for the JAK/STAT signaling pathway^49^, was particularly interesting because previous work demonstrated that signaling through the related ligand Upd and other JAK/STAT pathway components is required in the mushroom body for aversive olfactory long-term memory^50^. We therefore examined *upd3* mutants and mutants lacking both *upd3* and its close homologue *upd2*. Although the double-mutant flies showed no detectable reduction in SLI during Training sessions **(Extended Data Fig. 22**), both mutant groups showed a smaller increase in agarose avoidance from the Pre to Post period compared to controls (**Fig. 5g**), consistent with a contribution of Upd-family signaling to learned agarose avoidance. Together, the relatively small DEG sets and the identification of a functionally relevant candidate suggest that RNA-seq with yoked condition controls is a promising approach for identifying genes associated with spatial learning.

Together, these results show that genetic background strongly influences spatial learning performance, including the benefit flies derive from external olfactory landmarks. The significant line × chamber interaction suggests that learning in feature-poor environments and exploiting external olfactory landmarks are subject to partly distinct genetic influences. This finding is consistent with our earlier evidence that, although learning in the two chamber types relies on overlapping cue systems, exploiting external olfactory landmarks involves additional MB-dependent learning of their negative value (**Figs. 2-4**). Complementing these behavioral findings, RNA-seq revealed experimental-versus-yoked transcriptional differences that were shared across chambers as well as differences that depended on chamber type. The latter included *upd3* in *w^1118^* and *CG1722* in Line 7 (**Fig. 5c-f**). Follow-up mutant analyses provided functional evidence that *upd3* contributes to the increase in agarose avoidance from the Pre to Post period (**Fig. 5g**).

## Discussion

Our findings suggest that flies can learn the spatial significance of both self-generated and external olfactory cues and use this information together with self-motion cues to return to initially unmarked goals. Thus, navigation without prominent visual landmarks can draw on cues that animals themselves leave behind, as well as cues already present in the environment. In feature-poor small arenas, flies learn to find self-deposited scents near the goal attractive through MB-dependent learning and combine these odor cues with PFN-dependent self-motion information to guide their returns. When external agarose landmarks are introduced away from the goal, flies learn to treat them as aversive through MB-dependent learning and use them as guideposts to constrain search and improve performance. The same cue systems support learning in arenas approximately tenfold larger in area, although their relative contributions may depend on arena size and landmark arrangement. Variation in learning among F_1_ progeny of wild-derived strains suggests partly distinct genetic influences on learning in feature-poor arenas and the ability to benefit from external landmarks. Complementing these behavioral findings, RNA-seq reveals shared and chamber-dependent transcriptional differences between experimental and yoked flies, and functional analysis implicates *upd3* in contributing to learned avoidance of agarose.

A central and unexpected finding of this work is that self-deposited scents can serve as learned navigational cues. Our evidence includes the detection of fly-derived compounds on arena floors after training, reduced learning following manipulations expected to impair scent transfer, and reward-evoked behaviors consistent with deposition. Most directly, shifting the chamber’s transparent films biased subsequent search toward the displaced area in an olfaction-dependent manner. This use of self-generated cues is reminiscent of the construction of visual landmarks by desert ants^51^. We acknowledge that such deposition need not be deliberate: repeated visits to the reward site could concentrate deposits whose scents flies learn to find attractive. Male flies are known to actively deposit 9-tricosene in response to appetitive food odors^24^, but the absence of a detectable learning deficit in *Or7a* mutants argues against an essential role for 9-tricosene–Or7a signaling (**Extended Data Fig. 23**). Identifying the behaviorally active odorants and their cognate receptors will be an important next step; GC–MS provides candidates but may have missed relevant components below detection thresholds.

How might olfactory and self-motion cues work to support learning? Building on our results and prior work on path integration during local search^19–21^, we propose the following working model. In the feature-poor arenas, flies may use PFN-dependent self-motion information, potentially together with other signals, to construct a home vector – an estimate of the goal’s direction and distance relative to the fly – and learned olfactory cues could help correct this estimate (**Fig. 5h**). For example, movement-dependent changes in learned olfactory signals could provide feedback for updating the home vector. Although PFNd neurons have recently been shown to support sustained course correction^52^, learning deficits after brief inhibition at the goal argue against a strong role of ongoing steering impairment. Our perturbation experiments support contributions from both cue systems to spatial learning, consistent with their proposed joint role in guiding flies’ return to the goal. Residual learning when either system is compromised also suggests some capacity for independent navigation through dead reckoning or learned odor-guided simple approach and avoidance. Comparisons across two arena sizes further suggest that their relative contributions vary with spatial demands. In large feature-poor chambers, PFN-dependent information appears relatively more important for local returns, whereas olfactory cues become relatively more important during excursions farther from the goal. This pattern might reflect the accumulation of errors in self-motion-based estimates during longer excursions, increasing the value of olfactory cues. External olfactory landmarks improve learning in both small and large chambers, but their placement and spatial extent relative to the goal may influence how flies use them. In the small agarose chamber, learned odor avoidance may primarily constrain search to a restricted region; in the large agarose chamber, the landmarks may additionally provide positional information for correcting the putative error-prone home vector as animals move further away from goal.

How might the conceptual working model we proposed be implemented at the circuit level? Learning is known to alter how the odor-representing Kenyon cells drive the ensemble activity of mushroom body output neurons (MBONs) in response to learned odors. But beyond their roles in promoting simple approach and avoidance responses to learned odors, MBONs also provide input to the fan-shaped body (FB) circuitry, which also receives PFN input, offering a plausible route for learned olfactory signals to interact with self-motion information^53^. Whether this circuitry actually corrects errors in the home vector, and whether and how the specific reinforcer we used here resets the putative home vector to zero at the goal and contributes to error correction away from goal as suggested by a recent study^54^ are important questions to be investigated. We note that the lack of a detected requirement for EPG neurons is consistent with the proposed model. Although EPG’s allocentric heading signals support mapping of egocentric movements onto an allocentric frame^2,3^, the home vector in our proposed model is egocentric and could, in principle, be computed without allocentric heading signals. Updating this vector would nevertheless require information about the fly’s turns (angular self-motion). Turning-related activity in P-EN neurons innervating the noduli makes them candidates for supplying this information, although their contribution to navigation in our task remains unresolved^2^.

An intriguing finding from our RNA-seq analyses is that many stress-associated genes were more highly expressed in yoked than in experimental flies across both genetic backgrounds and chamber types, despite both groups receiving identical rewards at identical times. This consistent pattern raises the possibility that receiving rewards without a learnable spatial contingency produces a distinct stress-related transcriptional state. Whether this signature reflects the lack of control itself or associated behavioral and physiological differences remains unresolved. Understanding its basis could reveal how control over reward delivery shapes molecular responses to experience. Our F1 survey opens a complementary avenue for investigating how genetic variation tunes navigational circuits. Dissecting this variation could generate testable hypotheses about how specialized navigational abilities evolve across species.

## Methods

### Fly husbandry

Flies were raised at 25 °C on standard molasses food. We controlled the density of the breeding crosses and supplied them with yeast paste (6 g dry yeast in 10 mL of 0.5% propionic acid) to ensure the health of the progeny. Only 10- to 14-day-old males were used for our spatial learning tasks, to limit variability associated with female mating and reproductive state. They were collected on the day of eclosion and kept in regular food vials for 6-10 days at a density of 12-14 flies per vial. They were then transferred to food vials containing 400 μM all-trans-retinal and maintained in darkness for 2 days, followed by transfer to fresh retinal-containing food vials and approximately 2 days on a 12-h light/12-h dark cycle before being assayed.

### Fly genotypes by figure panel

Each cell in the left column lists figure panels that use data from the same group of flies. Different cells may share the same genotype but represent different groups of flies. For example, **Figs. 2l** and **2m** show two different behaviors measured in the same flies. The sources of the fly stocks are provided in **Supplementary Table 3**.

| Figure panel(s) | Genotype |
|---|---|
| **Fig. 1d-n, 2g, h (**Ctrl**), 3c** (Flat), **Extended Data Fig. 3a-e, 8g** (Ctrl), **9b** (Ctrl), **13a** (flat)**, c** (flat) | *UAS-CsChrimson /y; +/+; 0273-GAL4/+* |
| **Fig. 1p, Extended Data Fig. 8a** (Ctrl) |  |
| **Fig. 1q** |  |
| **Fig. 2n, 3g** (flat), **Extended Data Fig. 8i, 9c** |  |
| **Fig. 2g, h** (antennae removed), **Extended Data Fig. 8g** (antennae removed)**, 9b** (antennae removed) |  |
| **Fig. 2h** (other groups), **Extended Data Fig. 8g** (other groups) |  |
| **Fig. 2j** |  |
| **Fig. 3b, c** (agarose), **d, e** (Ctrl), **Extended Data Fig. 12a-g, 13a** (agarose)**, c** (agarose) |  |
| **Fig. 3d, e** (other groups) |  |
| **Fig. 3f, g** (agarose) |  |
| **Fig. 3h** |  |
| **Fig. 4f-h** |  |
| **Extended Data Fig. 2a-c** |  |
| **Extended Data Fig. 8a** (antennae removed) |  |
| **Extended Data Fig. 8a** (other groups) |  |
| **Extended Data Fig. 8c** |  |
| **Extended Data Fig. 10** |  |
| **Fig. 2l, m** |  |
| **Fig. 2b, r** (normal antennae)**, Extended Data Fig. 4e, 8k, 11b, 12h** (flat) | *UAS-CsChrimson /y; Empty-LexA/Otd-FLP; 0273-GAL4/LexAop-(FRT)-tdTomato-STOP-(FRT)-eGFP-Kir*<br>(*Ctrl>Kir*) |
| **Fig. 2r** (antennae removed)**, Extended Data Fig. 12h** (antennae removed, flat) |  |
| **Fig. 2p, Extended Data Fig. 8j, 9d** |  |
| **Fig. 3i** |  |
| **Fig. 3j, k** (normal antennae)**, Extended Data Fig. 12h** (agarose) |  |
| **Fig. 3k** (antennae removed)**, Extended Data Fig. 12h** (antennae removed, agarose) |  |
| **Fig. 4b-d, i-j** (normal antennae)**, k-q, s** (flat), **Extended Data Fig. 14b-j, 15e-f, h, i, k, l, n, o, 16k** (Ctrl, flat), **17b** (flat) |  |
| **Fig. 4d, j** (antennae removed)**, k-q** (antennae removed), **Extended Data Fig. 15d-f, 16k** (antennae removed, flat), **17c** (flat) |  |
| **Fig. 4s-v** (normal antennae), **Extended Data Fig. 16b-j, k** (Ctrl, agarose)**, 17b** (agarose) |  |
| **Fig. 4s-v** (antennae removed)**, Extended Data Fig. 16k** (antennae removed, agarose)**, 17c** (agarose) |  |
| **Extended Data Fig. 4a** |  |
| **Extended Data Fig. 5b, 6b** |  |
| **Extended Data Fig. 8d** |  |
| **Extended Data Fig. 8e** |  |
| **Extended Data Fig. 11d** |  |
| **Extended Data Fig. 14i, 15k, l** |  |
| **Extended Data Fig. 14j, 15n, o** |  |
| **Fig. 2b, r** (normal antennae)**, 3k, Extended Data Fig. 4e,12h** (flat) | *UAS-CsChrimson /y; R16D01-LexA/Otd-FLP; 0273-GAL4/LexAop-(FRT)-tdTomato-STOP-(FRT)-eGFP-Kir*<br>(*PFNd>Kir*) |
| **Fig. 2r** (antennae removed)**, Extended Data Fig. 12h** (antennae removed, flat) |  |
| **Fig. 3j, k** (normal antennae)**, Extended Data Fig. 12h** (agarose) |  |
| **Fig. 3k** (antennae removed)**, Extended Data Fig. 12h** (antennae removed, agarose) |  |
| **Fig. 4i-j** (normal antennae), **k-q, Extended Data Fig. 15g-l, 16k** (flat)**, 17e** (flat) |  |
| **Fig. 4j** (antennae removed)**, Extended Data Fig. 16k** (antennae removed, flat) |  |
| **Fig. 4u, v** (normal antennae)**, Extended Data Fig. 16k** (agarose)**, 17e** (agarose) |  |
| **Fig. 4v** (antennae removed)**, Extended Data Fig. 16k** (antennae removed, agarose) |  |
| **Extended Data Fig. 4a** |  |
| **Extended Data Fig. 5b** | *UAS-CsChrimson /y; R60D05-LexA/Otd-FLP; 0273-GAL4/LexAop-(FRT)-tdTomato-STOP-(FRT)-eGFP-Kir*<br>(*EPG>Kir*) |
| **Extended Data Fig. 14j, 15m-o, 17f** (flat) |  |
| **Extended Data Fig. 16j, 17f** (agarose) |  |
| **Extended Data Fig. 6b** | *UAS-CsChrimson /y; R37G12-LexA/Otd-FLP; 0273-GAL4/LexAop-(FRT)-tdTomato-STOP-(FRT)-eGFP-Kir*<br>(*PFR>Kir*). |
| **Fig. 2p, Extended Data Fig. 8j, 9d** | *UAS-CsChrimson/y; R19B03-LexA/Otd-FLP; 0273-GAL4//LexAop-(FRT)-tdTomato-STOP-(FRT)-eGFP-Kir*<br>(*MBKC-1>Kir*) |
| **Fig. 3i** |  |
| **Extended Data Fig. 8d** |  |
| **Extended Data Fig. 14i, 15j-l, 17d** (flat) |  |
| **Extended Data Fig. 16i, 17d** (agarose) |  |
| **Extended Data Fig. 11b, 8k** | *UAS-CsChrimson/y; R26E07-LexA/Otd-FLP; 0273-GAL4/LexAop-(FRT)-tdTomato-STOP-(FRT)-eGFP-Kir*<br>(*MBKC-2>Kir*) |
| **Extended Data Fig. 11d** |  |
| **Extended Data Fig. 8e** |  |
| **Fig. 2c, Extended Data Fig. 4f** | *w; Empty-LexA/lexAop-GtACR1; 0273-GAL4/UAS-CsChrimson*<br>(*Ctrl>GtACR1*) |
| **Extended Data Fig. 5c** |  |
| **Extended Data Fig. 6c** |  |
| **Fig. 2q, Extended Data Fig. 11c, 8l, 9e** |  |
| **Extended Data Fig. 11e, f** |  |
| **Extended Data Fig. 8f** |  |
| **Extended Data Fig. 4b** |  |
| **Fig. 2c, Extended Data Fig. 4f** | *w; R16D01-LexA/lexAop-GtACR1; 0273-GAL4/UAS-CsChrimson*<br>(*PFNd>GtACR1*) |
| **Extended Data Fig. 4b** |  |
| **Extended Data Fig. 5c** | *w; R60D05-LexA/lexAop-GtACR1; 0273-GAL4/UAS-CsChrimson*<br>(*EPG>GtACR1*) |
| **Extended Data Fig. 6c** | *w; R37G12-LexA/lexAop-GtACR1; 0273-GAL4/UAS-CsChrimson*<br>(*PFR>GtACR1*) |
| **Fig. 2q, Extended Data Fig. 8l, 9e** | *w; R19B03-LexA/lexAop-GtACR1; 0273-GAL4/UAS-CsChrimson*<br>(*MBKC-1>GtACR1*) |
| **Extended Data Fig. 8f** |  |
| **Extended Data Fig. 11f** |  |
| **Extended Data Fig. 8l, 11c** | *w; R26E07-LexA/lexAop-GtACR1; 0273-GAL4/UAS-CsChrimson*<br>(*MBKC-2>GtACR1*) |
| **Extended Data Fig. 8f** |  |
| **Extended Data Fig. 11e** |  |
| **Fig. 2e, Extended Data Fig. 4g, 5e** | *w; UAS-CsChrimson/Empty-GAL4-AD; 0273-GAL4/Empty-GAL4-DBD*<br>(*Ctrl>CsC*) |
| **Extended Data Fig. 4c** |  |
| **Fig. 2f, Extended Data Fig. 4h** |  |
| **Extended Data Fig. 4d** |  |
| **Extended Data Fig. 6e** |  |
| **Fig. 2e, Extended Data Fig. 4g** | *w; UAS-CsChrimson/SS00078; 0273-GAL4/SS00078*<br>(*PFNd>CsC*) |
| **Extended Data Fig. 4c** |  |
| **Fig. 2f, Extended Data Fig. 4h** | *w; UAS-CsChrimson/SS52628; 0273-GAL4/SS52628*<br>(*PFNv>CsC*) |
| **Extended Data Fig. 4d** |  |
| **Extended Data Fig. 5e** | *w; UAS-CsChrimson/SS00090; 0273-GAL4/SS00090*<br>(*EPG>CsC*) |
| **Extended Data Fig. 6e** | *w; UAS-CsChrimson/SS54549; 0273-GAL4/SS54549*<br>(*PFR>CsC*) |
| **Extended Data Fig. 7, 8h** | *w; UAS-CsChrimson/+; 0273-GAL4/Orco23130*<br>(*Orco2/+*) |
| **Extended Data Fig. 8b** |  |
| **Extended Data Fig. 7, 8h** | *w; UAS-CsChrimson/+; 0273-GAL4, Orco23129/Orco23130*<br>(*Orco1/Orco2*) |
| **Extended Data Fig. 8b** |  |
| **Extended Data Fig. 23** | *w; UAS-CsChrimson/+; 0273-GAL4/+*<br>(*Ctrl*) |
| **Extended Data Fig. 23** | *Or7a/y; UAS-CsChrimson/+; 0273-GAL4/+*<br>(*Or7a mutant*) |
| **Extended Data Fig. 2d, e** | *norpA/y; +; UAS-CsChrimson/0273-GAL4* |
| **Fig. 5a-d, f, Extended Data Fig. 18, 19, 21** | *14021-0231.**07**; UAS-(FRT.STOP)CsChrimson-tdTomato, Otd-FLP/14021-0231.**07**; 0273-GAL4/14021-0231.**07***<br>(Line 7) |
| **Fig. 5g, Extended Data Fig. 22** | w; UAS-(FRT.STOP)CsChrimson-tdTomato, Otd-FLP /+; 0273-GAL4/+<br>(Ctrl) |
| **Extended Data Fig. 1d** |  |
| **Fig. 5a-e, Extended Data Fig. 18-20** (*w<sup>1118</sup>* background flies) |  |
| **Fig. 5g** | *w, upd3[Delta]; UAS-(FRT.STOP)CsChrimson-tdTomato, Otd-FLP/+; 0273-GAL4/+*<br>(*upd3 KO*) |
| **Fig. 5g, Extended Data Fig. 22** | *w, upd2[Delta] upd3[Delta]; UAS-(FRT.STOP)CsChrimson-tdTomato, Otd-FLP /+; 0273-GAL4/+*<br>(*upd2 + upd3 KO*) |
| **Extended Data Fig. 1e** | *w; UAS-GtACR1/+; 0273-GAL4/+* |
| **Extended Data Fig. 1a** | *w; +; UAS-mCD8GFP/0273-GAL4* |
| **Extended Data Fig. 1b, c** | *w; 71G01-LexA/UAS-(FRT.STOP)CsChrimson-tdTomato; 0273-GAL4/lexAop-FLP* |
| **Fig. 2a** | *w; R16D01-LexA/lexAop-GtACR1.d.EYFP; 0273-GAL4* |
| **Fig. 2d** | *w; +/SS00078; UAS-eGFP-Kir/SS00078* |
| **Fig. 2d** | *w; +/SS52628; UAS-eGFP-Kir/SS52628* |
| **Fig. 2o** | *w; R19B03-LexA/lexAop-GtACR1.d.EYFP; 0273-GAL4* |
| **Extended Data Fig. 5a** | *w; R60D05-LexA/lexAop-GtACR1.d.EYFP; 0273-GAL4* |
| **Extended Data Fig. 5d** | *w; +/SS00090; UAS-eGFP-Kir/SS00090* |
| **Extended Data Fig. 6a** | *w; R37G12-LexA/lexAop-GtACR1.d.EYFP; 0273-GAL4* |
| **Extended Data Fig. 6d** | *w; +/SS54549; UAS-eGFP-Kir/SS54549* |
| **Extended Data Fig. 11a** | *w; R26E07-LexA/lexAop-GtACR1.d.EYFP; 0273-GAL4* |

Wild-derived *Drosophila melanogaster* strains were designated “Line X,” where X represents the last one to three digits of the corresponding stock number from the Cornell Drosophila Stock Center (e.g., Line 7 corresponds to stock 14021-0231.07). The SLI data shown in **Fig. 5a** were obtained from male **F<sub>1</sub>** progeny of crosses between females from the respective wild-derived lines and males of genotype *w*; *UAS-(FRT.STOP)CsChrimson-tdTomato, Otd-FLP/CyO; 0273-GAL4/TM2*.

### Non-genetic manipulations of flies

#### Removal of third antennal segments, aristae, and front tarsi

Depending on the experimental condition, we removed the third antennal segments, the aristae, or both the aristae and front tarsi with forceps one day before the assay to allow the flies to recover and adapt.

#### Gluing the antennae

Both antennae were covered with a small amount of UV-curable glue (Bondic) under a dissecting microscope, and the glue was cured with 5 s of exposure to 365-nm UV light (Darkbeam UV Flashlight). This manipulation was performed one day before the assay to allow the flies to recover and adapt.

#### Gluing the genitalia

The genitalia were covered with a small amount of UV-curable glue (Bondic) under a dissecting microscope, and the glue was cured with 5 s of exposure to 365-nm UV light (Darkbeam UV Flashlight). To minimize unintended activation of *0273-GAL4*-labeled neurons by the dissecting microscope illuminator (Nikon NI-150), we kept its intensity setting below 30% and limited the total illumination time to less than 3 min. Control flies were anesthetized and exposed to the same microscope illumination and UV light for the same durations but were not glued. Because the glue on the genitalia sometimes detached, we applied it 2-3 h before the assay and inspected the flies afterward to confirm that the glue was still present.

#### Removal of hind tarsi

Hind tarsi were removed with forceps 2-3 h before the assay, matching the timing of the genitalia-gluing procedure, because the two manipulations shared the same control group.

### Implementation of the spatial learning task

#### Hardware

We used two types of multi-arena apparatuses, containing 40 small arenas or eight large arenas, for studying spatial learning. Each apparatus was a removable chamber containing multiple individual arenas, whereas a setup comprised the apparatus, LED board, cameras and their supporting stand, and infrared backlight. The same stand could accommodate either apparatus type and its corresponding LED board.

#### High-throughput small-chamber apparatuses

Each small-chamber apparatus contained 40 individual arenas, each housing one fly. We built six setups, enabling us to assay up to 240 flies in parallel. In the flat chamber, each arena was rectangular (16 × 10 × 3 mm) and lacked prominent internal features (**Fig. 1a**). The agarose chamber had the same arena dimensions but included two troughs (3 × 10 × 3 mm) at opposite ends, filled with 1% agarose (**Fig. 3a**).

#### Moderate-throughput large-chamber apparatuses

Each apparatus contained eight individual arenas. In the large flat chamber, each arena was square (39 × 39 × 5 mm) and lacked prominent internal features (**Fig. 4a**). The large agarose chamber had the same arena dimensions but included four circular wells (8 mm in diameter) filled with 1% agarose (**Fig. 4r**).

#### Optogenetic stimulation

Light pulses were delivered by custom LED controller boards. In the small-chamber apparatuses, each arena was illuminated by two 5-mm green Cree LEDs (C503B-GAN-CB0F0791; 527 nm; 15° viewing angle). In the large-chamber apparatuses, each arena was illuminated by nine 5-mm red Cree LEDs (C503B-RAN-B0-RA-00; 624 nm; 15° viewing angle). For all genotypes and both LED board types, we used a 25% controller intensity setting for the 250-ms light pulses. This corresponded to approximately 16.4 μW/mm² at the arena floor in the small chambers and 9.03 μW/mm² in the large chambers. The original LED board schematic and layout are available in the SkinnerTrax GitHub repository.

#### Cameras

Each setup used two cameras (Microsoft LifeCam Cinema) with their internal IR-blocking filters removed and external IR-pass filters (LEE Filters 87) added.

#### Infrared backlight

An 850-nm light pad was placed underneath the arenas and LED controller board to provide background illumination for tracking. The light pads were custom-made by Junlong Display Factory, which replaced the white LEDs in their Artograph A920 light pads with 850-nm LEDs.

#### Software

SkinnerTrax^33^ was used to administer the learning task independently and in parallel to each fly in the 40- or 8-arena apparatuses. SkinnerTrax’s experimental records, which served as input for subsequent analysis, included the flies’ center positions and fitted ellipses for each frame, as well as the times when each fly was stimulated. The code for SkinnerTrax is available at https://github.com/ulrichstern/SkinnerTrax.

#### Handling of flies for the spatial learning task

Flies were anesthetized with CO₂ for loading into the chambers (and for the non-genetic manipulations described above). Circadian-entrained male flies (see Fly husbandry) were loaded into flat or agarose chambers, one fly per arena, within 2.5 h of light onset. The chambers were then placed into the tracking setup and shielded from ambient light with cardboard boxes.

For comparisons between groups differing in genotype, non-genetic manipulation, or chamber type, we generally reared flies, treated them with retinal, and circadian-entrained them in parallel. The groups were also assayed on the same days, typically over several days, to minimize differences in learning performance arising from differences in handling or day-to-day experimental conditions.

#### Task protocol

Experiments began with a 15-min Pre period followed by three 1-h Training sessions. Each Training session was followed by a 15-min Post period, except that RNA-seq samples were collected immediately after Training 3. Rewards were available only during Training sessions. Unless otherwise stated, behavioral analyses used Training 1 and Training 2 and their corresponding Post periods. Training 3 was analyzed in the film-sliding experiment; Post 3 measurements were used for the chamber-washing and agarose-avoidance analyses.

### Analysis of the spatial learning task

All behavioral analysis code used in this paper is available on GitHub. The main repository is https://github.com/rcalfredson/nvsl-analysis; code for reward-evoked behavior detection and associated analyses is available in two additional repositories: https://github.com/rcalfredson/kinelearn and https://github.com/rcalfredson/kinelearn-stimviz.

#### Real-time heatmaps

SkinnerTrax accumulated detected fly-center positions during each Training session in 2 × 2-pixel spatial bins. Each detected position incremented its spatial bin, and no walking-speed threshold was applied. Maps included experimental and yoked flies when present.

#### Detection of circular boundary crossings

Analyses involving entry or exit events across software-defined (“virtual”) circular boundaries were based on fly-center positions. To prevent noise from causing spurious entry or exit events, we used borders with small yet non-zero width, and registered events only when the border was fully crossed. For reward- and control-circle entry events, frames with missing tracking inherited the circle-classification state from the last tracked frame. Other circular-boundary analyses used interpolated fly-center positions according to our general missing-tracking policy.

#### Calculation of the reward index (RI)

The reward index (RI), which we introduced in prior work^35^, is defined as follows:

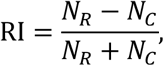

where *N_R_* and *N_C_* denote the numbers of reward-circle and control-circle entries, respectively. An entry into either circle is counted when the fly crosses the corresponding circular boundary from outside to inside. The reward and control circles are shown in **Figs. 1a** and **4a**, with control circle diameters of 12.5 mm in the small and 30 mm in the large chambers.

In our previous study^35^, the reward and control circles were equal in size and placed at spatially symmetric locations. In the present study, we use a larger control circle concentric with the reward circle. This design distinguishes repeated local returns to the reward circle from broader excursions away from the goal. It is particularly appropriate for the small chambers, where the reward circle is centered to minimize its association with fixed spatial features, such as walls and corners, leaving no spatially equivalent location for an equal-sized control circle. A fly that efficiently returns to the reward circle can repeatedly enter it without crossing the outer control-circle boundary, resulting in an RI approaching 1.

To align learning performance across flies, we synchronized the analysis intervals to the first reward received by each experimental fly in each Training session, rather than to the start of the session. The first reward defines time 0 for the experimental fly and its paired yoked control. We refer to each subsequent 10-min interval as a sync bucket (SB). For example, SB5 covers the interval from 40 to 50 min after the first reward. This synchronization excludes the initial interval during which the experimental fly had not yet received a reward in that Training session.

As described previously^35^, RI was designed to minimize sensitivity to differences in walking speed. Because RI measures the relative frequency of entries into two circles rather than their absolute numbers, a slower fly can achieve the same RI as a faster fly if their relative entry frequencies are equal. To limit sampling variability in RI estimates, we excluded intervals in which the total number of reward- and control-circle entries was less than 10. This criterion also excluded intervals in which flies moved so little that they rarely crossed either circle boundary. Consequently, flies with no qualifying intervals contributed no data to the corresponding mean RI values.

#### Calculation of the spatial learning index (SLI)

To quantify spatial learning while accounting for non-learning effects of reward exposure, we calculated the spatial learning index (SLI) as the difference in RI between each experimental fly and its paired yoked control:

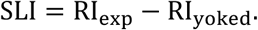

Because yoked controls received the same rewards at the same times as their paired experimental flies, but independently of their positions, they provide a baseline for behavioral changes caused by reward exposure without spatial learning. For example, repeated rewards might reduce sidewall walking and increase exploration of the chamber center, raising RI even without learning the reward location. Consistent with such effects, the RI of yoked controls was higher during Training than during the Post periods (**Figs. 1e**, **f)**, and their positional heatmaps showed increased center occupancy during Training relative to the Pre period (**Extended Data Fig. 13a**). Subtracting the yoked RI therefore reduces these non-learning contributions, including baseline spatial biases associated with the unequal reward and control circles. The same reasoning applies to genetic and sensory manipulations: effects on locomotion or spatial exploration that are shared by experimental and yoked flies are reduced by subtraction, making them less likely to be mistaken for effects on spatial learning.

Defining P(*R*) = *N_R_*/(*N_R_* + *N_C_*) as the fraction of circle entries occurring at the reward circle, RI can be rewritten as RI = 2P(*R*) − 1. Consequently,

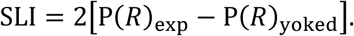

Thus, SLI is linear in the paired difference in reward-circle entry fractions, and yoked subtraction introduces no additional nonlinearity.

SLI can be calculated for any specified time interval. We refer to SLI for T2 SB5 as Final SLI; other analysis intervals are specified where applicable.

#### Calculation of rewards per distance (RPD)

As an alternative measure of spatial learning performance, we calculated rewards per distance (RPD) as

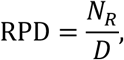

where *N_R_* denotes the number of reward-circle entries and *D* the cumulative distance traveled during the corresponding interval. RPD measures reward-acquisition efficiency, with distance traveled serving as a proxy for energy expenditure. By normalizing reward-circle entries by distance rather than time, RPD minimizes sensitivity to differences in walking speed.

To account for non-learning effects of reward exposure, we estimated the non-learning contribution to the experimental fly’s reward-circle entries as *N_R,yoked_*(*D*_exp_/*D*_yoked_), applying the yoked fly’s entry rate per unit distance to the experimental fly’s traveled distance. Subtracting this estimate from the experimental fly’s entry count and normalizing by its traveled distance gives

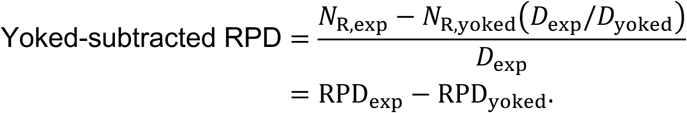

Thus, yoked-subtracted RPD combines distance normalization with the benefits of yoked subtraction described for SLI above. Consistent with its suitability as an alternative learning measure, yoked-subtracted RPD correlated well with SLI (**Fig. 1k**; **Extended Data Figs. 12b, 14c, and 16c**).

RPD was calculated separately for each sync bucket. For analyses spanning multiple sync buckets (e.g., T2 SB2–5), reward-circle entries and distances were summed over the selected buckets before division, rather than averaging the individual bucket-level RPD values. Yoked-subtracted RPD was then calculated from the corresponding pooled RPD values.

#### Calculation of reward preference

We used open-loop reward preference assays to assess flies’ preference for optogenetically delivered rewards. Two variants of the assay were used. To assess the effects of experimental manipulations on reward preference, we used a protocol consisting of a 30-minute Pre period, a 2-hour preference assay, and a 30-minute Post period. For analyses relating reward preference to SLI, a shorter protocol was conducted following the corresponding closed-loop spatial training experiment. This protocol consisted of a 15-minute Pre period followed by a 60-minute preference assay.

During the preference assay, only the upper half of the chamber was illuminated in a repeating cycle of 4 minutes of light on (7% LED intensity, which corresponds to about 4.63 μW/mm^2^ at the arena floors) followed by 4 minutes of light off.

For each fly, we pooled the time spent in the illuminated and unilluminated halves across all light-on intervals and calculated reward preference as follows:

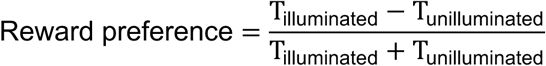

#### Classification of learner subgroups

Strong and weak learners were classified according to mean SLI over T2 SB2–5. Strong learners comprised the highest-ranked 20% of flies with a defined mean SLI, whereas weak learners comprised the lowest-ranked 50%. Mean SLI was calculated for flies with defined SLI values in at least three of the four sync buckets.

Fast learners were classified according to SLI for T1 SB1. Fast learners comprised the highest-ranked 20% of flies with a defined SLI in this bucket.

Percentile rankings were calculated separately within each experimental group. For each classification, the number of selected flies was determined by rounding down the specified fraction of flies with defined SLI values, with a minimum of one fly. Ties at selection boundaries were resolved according to the ordering of flies in the input data.

For comparisons of fast and strong learners, the two classifications were performed independently, allowing flies to belong to both categories. Flies with undefined SLI for one classification could still qualify for the other. In the corresponding scatter plots, only flies with defined SLI values on both axes were displayed, and Pearson correlations were calculated across all such flies, irrespective of subgroup membership.

#### Post-hoc heatmaps

Tracked fly-center positions were aligned in chamber-template coordinates and accumulated in bins approximately 2 × 2 template pixels in size. Missing internal positions were linearly interpolated, and only frames classified as walking using a 2 mm/s threshold contributed. Each histogram was divided by the total number of frames in its analysis window, and the resulting histograms were averaged separately across experimental and yoked flies, excluding flies with no walking-associated positions contributing to their histograms. The resulting values therefore represented the fraction of all frames in the analysis window during which the fly was walking within each spatial bin. Maps used the final 10 min of the Pre period, T2 SB5, the entirety of Training 2, or the first 3 min of the Post 2 period, as specified for each figure.

#### Exclusion of flies with poor tracking

Tracking quality was evaluated separately for each fly. Recordings with more than 99% missing fly-center positions were excluded from both spatial-learning and open-loop analyses. For spatial-learning recordings, tracking loss was calculated from the first successfully tracked frame to the end of the recording. Recordings were excluded if the proportion of missing frames exceeded 10% in small chambers or 60% in large chambers. The higher threshold for large chambers accommodated frequent tracking loss when flies walked on the sidewalls, away from the reward circle. We also excluded recordings with repeated suspicious tracking jumps, indicating that tracking could alternate between the fly and erroneous positions, potentially affecting reward delivery controlled by real-time tracking. When an experimental fly failed the tracking-quality criteria, its data were excluded; when only its yoked control failed, measurements requiring both flies were excluded, whereas independent measurements of the experimental fly could be retained.

#### Handling of missing tracking in occupancy, distance, and speed calculations

Internal spans of lost center-position tracking were linearly interpolated between the last position before the gap and the first position after it. Unless otherwise specified, successfully interpolated center positions were treated equivalently to originally tracked positions in frame-wise position-based measurements and behavioral classifications.

For cumulative-distance measurements, frame-to-frame displacements were calculated from the resulting position trace; an internal interpolated gap therefore contributed the straight-line distance between its bounding tracked positions. Isolated large displacements remained in cumulative path length, whereas trajectories containing repeated suspicious back-and-forth tracking jumps were excluded by the trajectory-level tracking-quality criteria described above.

Speed and ellipse-edge contact detection were exceptions to the default interpolation policy. A speed value assigned to frame *i*, representing displacement from frame *i − 1* to frame *i*, was retained only when both endpoint frames were originally tracked, because interpolation distributes the shortest-path displacement across a missing interval and can therefore systematically underestimate instantaneous speed. Likewise, contact detectors that required the fitted ellipse edge, including wall-contact detection, required intact tracking and ellipse measurements because ellipse orientation and axis lengths were not interpolated. Frames with lost tracking in these detectors were treated as unknown rather than carrying forward the preceding contact state, and contact episodes adjoining an unknown span were treated as censored and excluded; fully observed contact episodes were retained normally.

### Additional behavioral analyses

For the analyses of mean wall contacts per between-rewards trajectory, tortuosity, wall-contact trajectory length, dual-circle turnback ratio, and turnback home-vector alignment, the initial cohort comprised experimental flies with a defined Final SLI. Each metric was then subject to its own eligibility criteria, including minimum numbers of qualifying trajectories or episodes, as specified below.

#### Wall contact detection

Wall-contact events were identified frame by frame from the fly’s fitted ellipse. For each frame, the point on the ellipse nearest to any chamber wall was determined. Contact was then classified using a dual-boundary, or hysteresis, criterion: an event began when this point entered the region within 0.7 mm of the nearest wall and continued until it moved back beyond a boundary 1.0 mm from the nearest wall. A single contact state was maintained across the four walls, so contact remained continuous if the nearest wall changed during an event. The boundary offsets were selected from four candidate pairs (0.4/0.75 mm, 0.6/0.9 mm, 0.7/1.0 mm, and 0.9/1.2 mm) based on which provided the best agreement between algorithmically detected and manually labeled events. The first detected wall-contact episode in each recording was excluded because tracking could begin after the fly had already established contact with the wall, making the onset and duration of the initial episode uncertain.

#### Calculation of sharp-turn probability at different distances from the reward center

We quantified sharp turns during brief excursions beyond virtual circular boundaries centered on the reward-circle center. Separate analyses were performed using circle radii of 2, 3, 4, and 5 mm. For the analyses shown, sharp-turn probabilities were calculated during T2 SB5.

a. *Detection of outside-boundary events.* At each radius, circular crossings were detected as described above, using a one-pixel-wide border band outside the circle radius. An outside-boundary event began when the center of the fitted ellipse crossed outward beyond the outer edge of this band and ended when it crossed inward past the inner edge. Events were represented as half-open frame intervals, such that the first frame classified as outside was included, and the first frame reclassified as inside was excluded. Each radius was analyzed independently. ***b.*** *Detection of sharp turns.* Outside-boundary events lasting no more than 0.75 s were evaluated for sharp changes in movement direction. For each event, we examined the fly’s velocity angle—the direction of its movement—across the complete boundary-crossing trajectory, including the segment entering the outside region, all segments beginning outside the circle, and the segment crossing back inside. Consequently, turns could be detected even when only one frame lay outside the circle. Each velocity-angle sample was paired with the speed calculated for the same movement segment and was included only when that speed was at least 2 mm/s. Samples associated with slower speeds were skipped, after which angular changes were calculated between successive eligible velocity angles. Each angular change was circularly wrapped to the interval [-π, π), preserving its sign, and the signed changes were summed across the event. Thus, changes in a consistent clockwise or counterclockwise direction accumulated, whereas reversals in direction partially canceled. An event was classified as a sharp turn when the magnitude of the summed change in velocity angle was at least 90°. ***c.*** *Probability calculation.* Sharp-turn probability was calculated separately at each radius as the number of outside-boundary events classified as sharp turns divided by the total number of outside-boundary events whose starts fell within the analysis period. The denominator included all detected outside-boundary events beginning within the analysis period, including those exceeding the 0.75-s duration threshold for sharp-turn classification. ***d.*** *Exclusion criteria.* At each radius, a probability was retained only when the fly completed at least 10 outside-boundary events during the selected analysis period. For plotting, exclusions were pooled across radii: if either member of a paired recording lacked a valid probability at any radius, both members were excluded at every radius. Consequently, the same flies and sample sizes were used at all distances.

#### Calculation of proportion of time within reward-centered area

We defined a virtual circle with a radius of 2.5 mm centered on the reward-circle center. On each frame, the center of the fitted body ellipse was classified as inside the region when its radial distance from the reward-circle center was less than 2.5 mm. Because this calculation measured framewise occupancy rather than entries or exits, it did not use the finite-width border bands employed for circular boundary-crossing events. The proportion of time within the reward-centered area was calculated as the number of frames classified as inside divided by the total number of frames in the analyzed interval. For the analysis shown, the proportion was calculated across the entire Post 3 period for trained flies and the corresponding time-matched observation period for naïve flies, which underwent the same experimental schedule without reward training.

#### Calculation of Pre-period speed

Mean locomotion speed was calculated for experimental flies during the 10-minute interval preceding the LED-on event marking the end of the Pre period, shortly before training began (**Extended Data Figs. 3d , 4e–h and 8g–l**). Framewise speed was calculated from the displacement between consecutive tracked positions and converted to mm/s. Measurements spanning originally missing tracking were excluded, as described above. Stationary and slow-moving frames were retained; no walking-speed threshold was applied. At least 100 qualifying frame-to-frame speed measurements were required to report a value. A valid RI or SLI was not required.

#### Calculation of reward rate

Reward rate quantified the frequency of calculated rewards for the experimental fly in rewards per minute. For analyses spanning a fixed set of sync buckets, the total number of calculated rewards across the selected buckets was divided by the total duration of those buckets. For pooled reward-rate calculations, all selected sync buckets were required to be complete and available. A reward rate of zero was retained when no calculated rewards occurred within a valid analysis window.

In the reward-rate-versus-SLI analysis, reward rate was calculated for both T1 SB1 and across T2 SB2–5. To quantify initial repeat reward-acquisition rate, we measured the elapsed time from the first through the tenth calculated reward during Training 1 and divided the nine intervening reward intervals by this duration in minutes. Flies that did not reach ten calculated rewards during the selected period were excluded from the initial-rate analysis.

#### Calculation of Pre-period chamber-floor exploration

The rectangular chamber-floor region associated with each fly was divided into 1 × 1-mm spatial bins, including partial bins along the edges. For each position during the entire Pre period, the bin containing the center of the fitted body ellipse was marked as visited. Successfully interpolated center positions were included, whereas positions with coordinates outside the chamber-floor bounds were excluded. The fraction of the floor explored was calculated as the number of bins visited divided by the total number of bins. For the analysis relating Pre-period exploration to SLI, the value for the experimental fly was used.

#### Calculation of COM distance from reward center

A between-rewards trajectory extended from the frame of one recorded reward-delivery event through the frame preceding the next recorded reward-delivery event. For each trajectory from an experimental fly, the center-of-mass (COM) position was calculated by independently averaging the horizontal (x) and vertical (y) coordinates of the fitted-ellipse center across frames. COM distance was calculated relative to the center of the reward circle.

Between-rewards trajectories containing sidewall contact were included. Each trajectory was assigned to the 10-minute sync bucket with which it had the greatest temporal overlap, with exact ties assigned to the earlier bucket. For each fly and bucket, COM distance was calculated as the mean of the individual trajectory-level COM distances. At least five between-rewards trajectories were required to report a value.

#### Calculation of return-leg length

Return-leg length was calculated separately for each between-rewards trajectory. For each trajectory, the fly’s radial distance from the reward-circle center was calculated at each frame, and the frame at which the maximum distance was reached was identified. If multiple frames had the same maximum radial distance, the earliest such frame was selected. The return leg extended from this frame through the frame of the next recorded reward-delivery event, including the final frame-to-frame segment leading to that event. Return-leg length was calculated as the cumulative frame-to-frame path length over this interval and converted to millimeters; thus, it measured the complete return path rather than the straight-line displacement toward the reward center.

Each qualifying trajectory was assigned to the 10-minute sync bucket with which it had the greatest temporal overlap, with exact ties assigned to the earlier bucket. For each fly and sync bucket, return-leg lengths were averaged across trajectories, with at least five qualifying trajectories required to report a value.

#### Calculation of wall contacts per between-rewards trajectory

For each between-rewards trajectory, we counted wall-contact events whose onset occurred within that trajectory. Each event was counted once, according to the trajectory containing its onset. Only between-rewards trajectories with both bounding reward events within the selected analysis window were included. For the analysis shown, between-rewards trajectories bounded by reward events within T2 SB2–5 were pooled across that period. For each fly, the mean number of wall contacts per between-rewards trajectory was calculated as the total number of wall-contact events divided by the number of included trajectories. At least five between-rewards trajectories were required to report a value.

#### Calculation of the dual-circle turnback ratio

Turnbacks were evaluated using pairs of virtual circles concentric with the reward circle. The inner and outer radii were defined as absolute distances from the reward-circle center, and each radius pair was analyzed independently. For the analyses shown, inner/outer radius pairs of 3/5, 8/10, and 13/15 mm were used. Circular crossings were detected as described above, using a 0.1-mm border band outside the inner-circle radius and no border band at the outer-circle boundary.

A turnback episode began when the fly crossed outward beyond the inner circle. The episode ended at the first of two outcomes: crossing inward past the inner circle, classified as a turnback, or crossing outward past the outer circle, classified as a non-turnback. Episodes that reached the end of training without either outcome were considered right-censored and excluded from both the numerator and denominator.

For each fly and radius pair, the turnback ratio was calculated as the number of turnback episodes divided by the total number of episodes. Episodes were pooled across T2 SB2–5, and an episode contributed only when its complete span fell inside that pooled interval. At least five resolved episodes and a full SB5 of T2 were required to report a ratio for a given radius pair. Episodes involving wall contact were retained.

#### Calculation of turnback home-vector alignment

We quantified how closely the fly’s direction of movement was aligned with the direction of the reward-circle center when it completed a successful dual-circle turnback episode, as defined above. The 3/5, 8/10, and 13/15 mm inner/outer-radius pairs were analyzed independently. For each successful turnback, the re-entry point was estimated as the intersection of the re-entry trajectory segment with the nominal inner-circle boundary. The home vector extended from this intersection point toward the reward-circle center.

The fly’s movement direction at re-entry was estimated from its tracked positions in four frames surrounding the crossing. If e denotes the re-entry frame—the first frame classified as inside after the inward crossing—the mean position at frames *e – 2* and *e – 1* was calculated, as was the mean position at frames *e* and *e + 1*. The displacement from the former mean position to the latter defined the movement-direction vector. At most one of the four position samples could be a position interpolated across lost tracking.

Alignment was calculated as the cosine similarity between the movement-direction and home vectors: the dot product of the two vectors divided by the product of their magnitudes. Episodes for which either vector had zero magnitude or the alignment was otherwise undefined were excluded from the calculation. Alignment values ranged from 1 for movement directly toward the reward-circle center, through 0 for perpendicular movement, to -1 for movement directly away from the center. Values were averaged across successful re-entry episodes to obtain one mean alignment value per fly and radius pair.

Episodes were pooled across T2 SB2–5, and an episode contributed only when its complete span fell inside that pooled interval. At least five scorable episodes were required per fly for each radius pair. Episodes involving wall contact were retained.

Because alignment was calculated only for successful turnbacks, it quantified differences in the direction of movement at re-entry, ranging from more direct to more oblique approaches toward the reward center, and should not be interpreted as a measure of orientation toward the reward center across all excursions.

#### Calculation of the agarose avoidance ratio

Agarose avoidance was quantified using pairs of concentric circular boundaries around each of the four agarose regions in the large chamber. The inner circle coincided with the 4-mm-radius boundary of the agarose region, and the outer circle had a radius of 5 mm. Corresponding virtual circular regions were analyzed in flat-chamber controls. Circular crossings were detected as described above, using hysteresis with a 0.1-mm finite-width border applied only to the outer circles.

An approach episode began when the center of the fly’s fitted body ellipse entered any outer circle and ended when it subsequently exited all outer circles. The episode was classified as a contact if the fly entered any inner circle during the episode and as an avoidance otherwise. Episodes were detected using the combined outer-circle regions, preventing simultaneous approaches to different regions from being counted separately. Episodes lacking an observed outer-circle entry or exit were excluded.

For each fly and analysis period, the agarose avoidance ratio was calculated as the number of avoidance episodes divided by the total number of approach episodes. Episodes were assigned to analysis periods according to their outer-circle entry frames. Values were calculated independently for each 10-minute sync bucket during Training 1 and Training 2; the final 10 min of the Pre period were also analyzed. At least five approach episodes were required to report a ratio for a fly in a given period.

#### Calculation of the percentage of time spent within agarose or agarose-equivalent areas

In the small chamber, the agarose areas were two rectangular regions spanning the width of the chamber floor and extending 3 mm inward from its top and bottom edges. For flat-chamber controls, corresponding virtual regions with the same positions were treated as agarose-equivalent areas. On each frame, the center of the fitted ellipse was classified according to whether it fell within either rectangular region; region membership was evaluated independently on each frame without an entry–exit hysteresis threshold. The percentage of time spent within these areas was calculated as 100 times the number of frames classified as inside divided by the total number of frames in the interval. For the analysis shown, percentages were calculated during the final 10 min of the Pre period and T2 SB5.

#### Calculation of increase in agarose avoidance

Using the agarose-area definitions and percentage-of-time calculation described above, percentages were calculated during the final 10 min of the Pre period and the final 10 min of the Post 3 period. The increase in agarose avoidance was calculated by subtracting the Post 3 percentage from the Pre-period percentage, yielding a difference expressed in percentage points. Thus, positive values indicated an increase in agarose avoidance.

#### Calculation of tortuosity

Tortuosity was calculated separately for each between-rewards trajectory. For each trajectory, the cumulative frame-to-frame path length, *L*, including the final frame-to-frame segment leading to the next recorded reward-delivery event, and the maximum radial distance of the fitted-ellipse center from the reward-circle center, *R_max_*, were calculated over the same between-rewards trajectory. Tortuosity was defined as *L / R_max_*. Larger values indicated a more circuitous trajectory relative to its maximum radial extent.

For the analysis shown, the complete between-rewards trajectory was used, and trajectories involving wall contact were retained. A single value was obtained for each fly by averaging the trajectory-level tortuosity values across T2 SB2–5. At least five qualifying between-rewards trajectories were required to report a fly-level mean.

#### Calculation of wall-contact trajectory length

A between-rewards trajectory was classified as a wall-contact trajectory when at least one wall-contact event began between its two bounding reward events. Its length was calculated as the cumulative frame-to-frame path length between those reward events and converted to millimeters. A trajectory containing multiple wall-contact events contributed one trajectory-length value.

For the analysis shown, wall-contact trajectories bounded by rewards within T2 SB2–5 were pooled across that period. For each fly, mean wall-contact trajectory length was calculated by averaging the individual trajectory lengths. At least five wall-contact trajectories and a full SB5 of T2 were required to report a fly-level mean.

### Courtship-related behavioral assays involving *0273-GAL4*-labeled neurons

#### Unilateral wing extension

We placed one male *0273>CsChrimson* fly in each arena of the small flat chamber. Beginning 80 s after the start of the assay, green LED illumination (527 nm, 16.4 μW/mm^2^) was alternately turned on and off at 60-s intervals using the SkinnerTrax LED board. Unilateral wing extension was subsequently scored from the recordings using Behavioral Observation Research Interactive Software (BORIS).

#### Copulation

We placed one male *0273>GtACR1* fly and two virgin *w*¹¹¹⁸ females in each arena of a deep agarose chamber, normally used for egg-laying assays. We used 1% agarose containing 1% apple cider vinegar to promote copulation. All board LEDs were turned on continuously for 3 h, illuminating the chamber with green light (527 nm) with the level intensity set at 6.4 μW/mm^2^. We recorded the time when copulation began for retinal-fed and no-retinal control groups.

### Odor sample collection and GC-MS

To collect fly-deposited substances, we placed a 3.5-mm-diameter piece of transparent film at the center of each arena before the task and collected the films immediately after training. We pooled 18 films from arenas containing experimental flies, 18 from yoked-control arenas, and 18 from empty arenas (blank controls) into three separate vials, each containing 100 µL of hexane. The samples were sent for gas chromatography-mass spectrometry (GC-MS) analysis at Alera Labs (https://aleralabs.com/).

At Alera Labs, the extracts were evaporated and redissolved in 100 µL of hexane containing 5 µg/mL methyl nonadecanoate as an internal standard. A 3-µL aliquot of each sample was analyzed using an Rxi-5MS column (30 m length, 0.25 mm inner diameter, 0.25 µm film thickness) with an inlet temperature of 280 °C and a detector temperature of 300 °C. Helium was used as the carrier gas with a 10:1 split ratio. The oven temperature program was increased from 120 °C to 300 °C at 6 °C/min and held at 300 °C for 5 min. The mass spectrometer operated in electron ionization (EI) mode with an m/z scan range of 35-550. Compounds were putatively identified based on their highest-probability spectral matches.

### High-resolution recording of reward-evoked behaviors

We built a setup to record thorax-fixed flies walking on a wheel at 60 frames/s under infrared illumination. A typical trial consisted of a 1-min period without stimulation, followed by 2 min during which 24 red-LED pulses (250 ms, 14 μW/mm² measured at the fly’s position) were delivered at 0.2 Hz (5-s intervals). This stimulation frequency approximated the average reward rate of experimental flies near the end of the learning task in the flat chamber.

#### Automated behavior classification and stimulus-aligned analysis

Fly pose was estimated using DeepLabCut. The model tracked 13 keypoints: the head, thorax, abdomen, abdomen tip, genitalia tip, left and right front-, mid-, and hind-leg tips, and left and right wingtips. The DeepLabCut model was trained using manually labeled frames from 40 videos sampled from a dataset of 121 videos comprising 59 retinal-fed and 62 non-retinal-fed flies.

The resulting keypoint trajectories were used to train separate bidirectional long short-term memory (BiLSTM) classifiers for hind-leg grooming and genitalia extension. Ground-truth behavior bouts were manually annotated in BORIS and exported for model training and evaluation. Model inputs comprised thorax-centered keypoint coordinates normalized by the head-thorax distance; framewise velocity and acceleration features; selected inter-keypoint distances and three-point angles; and the DeepLabCut likelihood associated with each keypoint. Absolute image coordinates were excluded. Inputs were divided into overlapping 60-frame windows with a 10-frame stride. Each classifier contained two bidirectional LSTM layers with 128 and 64 units, respectively, followed by time-distributed dense layers with 32 and 64 units and a sigmoid output giving a behavior probability for each frame. Models were trained separately for each behavior using focal loss, a batch size of 8, and a learning rate of 0.001. Model checkpoints were retained when validation F1 improved. Multiple classifiers were trained using different video-level training and validation splits, with approximately 90% and 10% of the development videos assigned to training and validation, respectively. One classifier per behavior was selected based on validation F1.

The selected classifiers were applied to all 121 videos. Predictions from overlapping windows were averaged to obtain one probability per frame. Behavior bouts were then identified using a probability threshold of 0.6. Bouts were retained when they contained at least 16 above-threshold frames, and interruptions of up to three consecutive below-threshold frames were bridged within a bout.

Classifier performance was assessed on an evaluation set of 16 videos (15 retinal-fed and one non-retinal-fed) using manually annotated behavior bouts as ground truth. These videos were excluded from training and validation-based checkpoint selection, although they were used for tuning the focal-loss α parameter. For bout-level evaluation, a predicted bout was considered correctly detected when it overlapped an annotated bout by at least 20% of the predicted bout’s duration. Precision and recall were calculated from the numbers of correctly detected bouts, unmatched predicted bouts, and unmatched annotated bouts, with counts pooled across evaluation videos. The selected hind-leg grooming classifier achieved a precision of 0.875, recall of 0.804, and F1 score of 0.838; the corresponding values for genitalia extension were 0.914, 0.864, and 0.888, respectively.

For each fly, the binary behavior traces were aligned to the final 10 stimulus events using a window extending one-third of this interval before stimulus onset and two-thirds after onset (1.67 s before to 3.33 s after; stimulus onset defined as time 0). At each aligned frame, behavior probability was calculated as the fraction of these 10 stimulus presentations during which a behavior bout was active. The resulting fly-level probability traces were then averaged across flies within each experimental group. Error bands show 95% confidence intervals for the group mean, calculated using Student’s *t* distribution.

### Large-chamber film-sliding experiment

Before loading the flies, we covered the floor and underside of the lid of each large arena with separate sheets of transparent film. After Training 2, we slid both films by 20 mm (approximately half the arena length), ensuring that no fly was walking on either film during the movement. For mock-slide controls, the films were moved and then returned to their original positions. Flies remained in their arenas throughout, and Training 3 was then conducted.

#### Calculation of changes following film sliding

To quantify changes in SLI and spatial occupancy, we compared T2 SB5 (before sliding) with T3 SB1 (after sliding) in the same flies. Changes in SLI were calculated as ΔSLI = SLI_T3 SB1_ − SLI_T2 SB5_. Both SLI values were required to be defined for a fly to contribute to this analysis. Changes in spatial occupancy were calculated for two circular regions (20 mm in diameter): one centered on the reward location and another centered 20 mm below it, corresponding to the shifted location of the previously rewarded film area. Occupancy was the percentage of frames in which the experimental fly’s center was inside the region. Changes in occupancy were calculated as the percentage during T3 SB1 minus that during T2 SB5, expressed in percentage points. A fly was included when both occupancy measurements were defined.

### RNA sequencing of fly heads after the spatial learning task

#### Sample collection

We dissected fly heads from experimental flies and their paired yoked controls immediately after Training 3. Each experimental-yoked sample pair contained the same number of heads (15-20 per sample). The samples were placed in TRIzol reagent immediately after dissection and stored at -80 °C overnight. The following day, the samples were ground and homogenized using a QIAshredder, and RNA was extracted using an RNeasy Mini Kit (QIAGEN). RNA concentration and integrity were measured using a Qubit fluorometer (Invitrogen) and Bioanalyzer (Agilent), respectively. For each chamber type (flat and agarose), eight experimental-yoked sample pairs from the *w*¹¹¹⁸ background and six pairs from Line 7 were selected for RNA-seq based on RNA quality (**Fig. 5b**).

#### Sequencing and gene counting

Novogene performed RNA sequencing using Illumina NovaSeq 6000 sequencers to generate 150-bp paired-end reads and processed the sequencing data to obtain raw gene counts. After removing adapter sequences and low-quality reads, the remaining reads were aligned to the *D. melanogaster* reference genome (BDGP6) using HISAT22.0.5. Transcripts were assembled using StringTie 1.3.3b, and raw counts were generated using featureCounts 1.5.0-p3.

### Internal data analysis

We analyzed the two genetic backgrounds separately for differential gene expression analysis using DESeq2 1.46.0 in R. To reduce noise while retaining potentially relevant lowly expressed genes, we required at least five raw counts in at least eight of the 32 *w*¹¹¹⁸ samples or six of the 24 Line 7 samples, matching the number of samples per chamber × condition group. For *w*¹¹¹⁸, we used the DESeq2 design formula ∼ run + chamber + cond + chamber:cond, accounting for the two sequencing runs. For Line 7, we used ∼ chamber + cond + chamber:cond. Here, cond denotes the experimental versus yoked condition. DESeq2 Wald tests were used for differential-expression contrasts and chamber × condition interactions. P values were adjusted across genes using the Benjamini–Hochberg procedure, separately for each comparison. Genes with adjusted *P* < 0.05 were considered differentially expressed. Differential-expression results were used to generate volcano plots and Venn diagrams, while DESeq2-normalized counts were used for gene-expression plots. We did not directly compare gene expression between the two genetic backgrounds because different gene-annotation (GTF) versions were used for the original *w*¹¹¹⁸ and subsequent Line 7 datasets.

### Statistics

Statistical analyses were performed using GraphPad Prism 11 and custom code; RNA-seq analyses were performed using DESeq2 in R, as described above. Unless otherwise specified, each biological replicate in behavioral experiments represents an individual fly, and sample sizes are indicated in the figures or figure legends. We generally used parametric tests: unpaired *t*-tests with Welch’s correction for comparisons between independent groups, paired *t*-tests for paired measurements, and one-sample *t*-tests for comparisons against zero. Comparisons involving multiple groups or experimental factors were performed using one-way Welch ANOVA or two- or three-way ANOVA, with multiple-comparisons tests specified in the figure legends. Repeated measurements at different distances within flies were analyzed using mixed-effects models. All *t*-tests and correlation tests were two-sided. Unless otherwise specified, plotted means are accompanied by 95% confidence intervals (CI).

#### Statistical analysis of line plots

Unless otherwise specified, line plots show the arithmetic mean across flies at each plotted point, with shaded regions representing two-sided 95% confidence intervals of the mean calculated using Student’s t distribution. Undefined per-fly values were omitted independently at each plotted point. This convention was used for measurements plotted across sync buckets as well as for other continuous or ordered analysis parameters.

For analyses across sync buckets, two independent groups were compared separately at each bucket using two-sided Welch’s t-tests. Per-bucket p values were not adjusted for multiple comparisons. Paired experimental and yoked-control flies were compared using two-sided paired t-tests, with a pair included only when both values were defined at that bucket. For single-group difference measures, values were compared with zero using two-sided one-sample t-tests.

Where an overall comparison across a series of plotted time points was reported, a signed area under the curve (AUC) was calculated separately for each fly by trapezoidal integration. For analyses across Training sync buckets, integration included all five sync buckets (SB1-SB5). AUC was calculated only when all values included in the integration were defined; consequently, AUC comparisons could include fewer flies than individual plotted points. For two independent groups, AUC values were compared using a two-sided Welch’s t-test. For paired experimental-versus-yoked data, the within-pair difference in AUC was tested against zero using a two-sided one-sample t-test, equivalent to a paired t-test of the two AUC series. Signed AUCs for single-group difference measures were likewise tested against zero using two-sided one-sample t-tests.

#### Correlation analysis of fly-level measurements

Associations between fly-level measurements were evaluated using Pearson’s product-moment correlation. Flies lacking a defined value for either measurement were excluded from the corresponding analysis. The number of included flies, Pearson correlation coefficient r, and two-sided p value were reported. When p < 0.05, an ordinary least-squares linear trend line was fitted to the plotted observations and displayed.

For measurements summarized across multiple sync buckets, the aggregation procedure depended on how the metric was defined. For metrics defined as the mean of bucket-level values, the mean was calculated across SB2–5 of the selected Training and required at least three of the four bucket-level values to be defined; undefined bucket values were omitted from the mean. For metrics calculated from pooled data, the underlying data were combined across the selected interval before the measurement was calculated, and the measurement-specific eligibility threshold was applied to the pooled data.

#### Data and code availability

The raw data and resource information that support the findings of this study are available from the corresponding author upon request.Code for administering the spatial-learning task is available in the SkinnerTrax repository at https://github.com/ulrichstern/SkinnerTrax Code for analyzing spatial-learning behavior is available at https://github.com/rcalfredson/nvsl-analysis.

Code for reward-evoked behavior detection and associated analyses is available at https://github.com/rcalfredson/kinelearn, https://github.com/rcalfredson/kinelearn-stimviz

## Acknowledgements

We would like to thank the Bloomington Stock Center for providing fly strains, the Duke Physics Shop for fabricating the behavior apparatuses, and Alera Labs (https://aleralabs.com/) for GC-MS analysis. We would also like to thank Drs. Calakos, Nowicki, Volkan, and Dong, members of the Yang Lab, and Chengcheng Du from the Volkan Lab at Duke University for helpful comments and suggestions. This work was supported by an invertebrate neuroscience research grant awarded to C.-H.Y.

## Author contributions

Y.C., U.S., and C.-H.Y. designed the project. Y.C. performed most of the experiments and analyzed the behavioral data; X.Z. conducted additional behavioral experiments. Y.C. and U.S. independently analyzed the RNA-seq data. R.A. developed the behavioral data-analysis code, building on earlier code written by U.S. U.S. built the behavior-dependent stimulation system. Y.C., R.A., U.S., and C.-H.Y. interpreted the data. D.M. dissected fly brains for imaging and prepared RNA samples together with Y.C., X.Z., and S.F. Y.C. wrote the initial draft of the manuscript, which was substantially revised by R.A., X.Z., U.S., and C.-H.Y.

---

**Extended Data Fig. 1.**
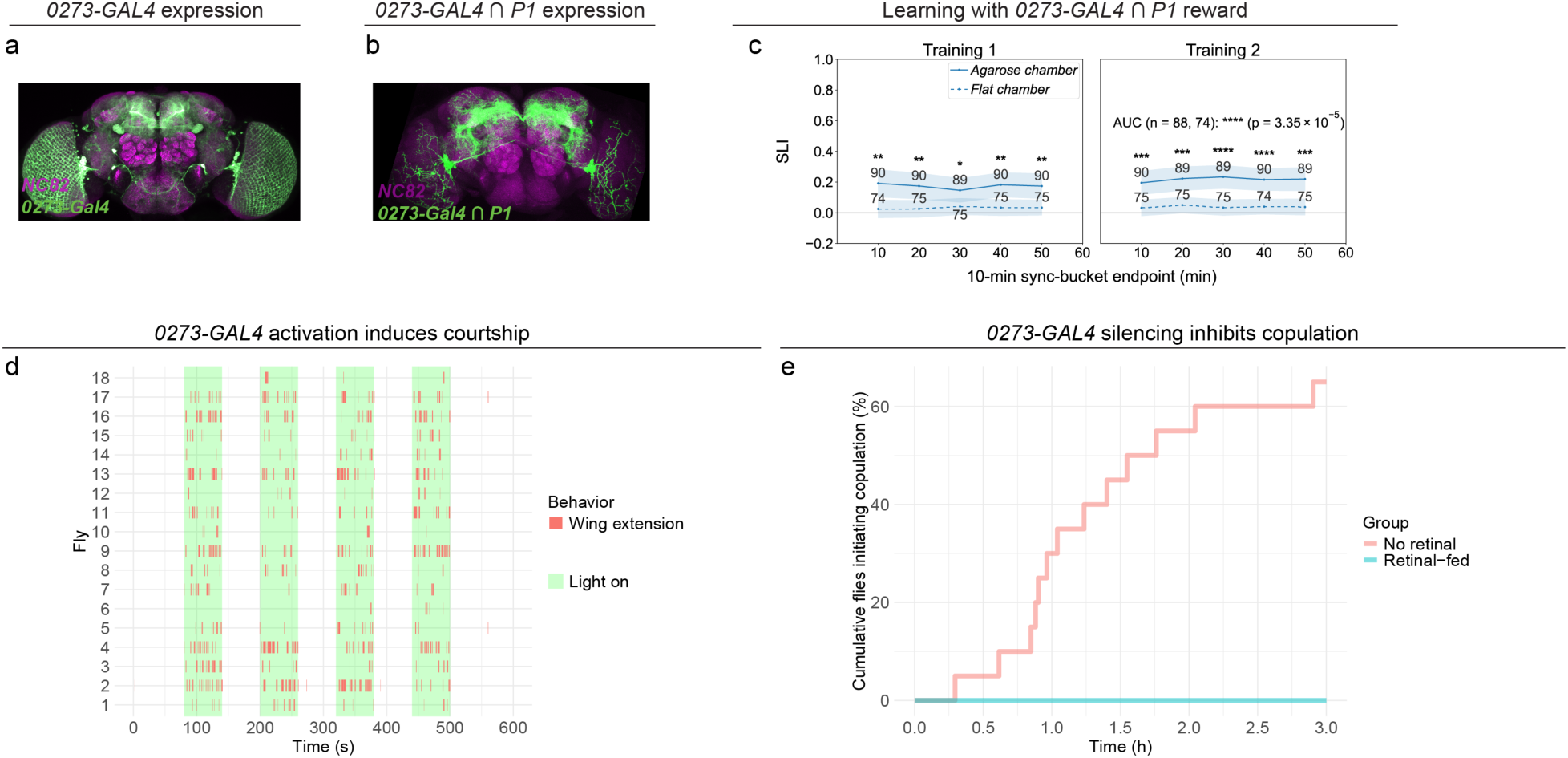
*0273-*GAL4 labels P1 *fruitless*-expressing neurons whose activation can also serve as a reinforcer. **a,** Expression pattern of *0273-GAL4*. **b,** Neurons labeled by both the P1 driver (*71G01-LexA*) and *0273-GAL4*. **c,** SLI of flies receiving optogenetic activation of neurons labeled by both the P1 driver and *0273-GAL4* as a reward in flat versus agarose chambers. **d,** Unilateral wing extension during optogenetic activation of *0273>CsChrimson* flies. **e,** Cumulative percentage of *0273>GtACR1* flies initiating copulation over time in the copulation assay (see **Methods**), with retinal (n = 20) or without retinal (n = 19; GtACR1 nonfunctional). In **c**, line plots show means and shaded 95% confidence intervals; numbers indicate sample sizes for each sync bucket. Throughout the paper, *: *p* < 0.05; **: *p* < 0.01; ***: *p* < 0.001; ****: *p* < 0.0001.

**Extended Data Fig. 2.**
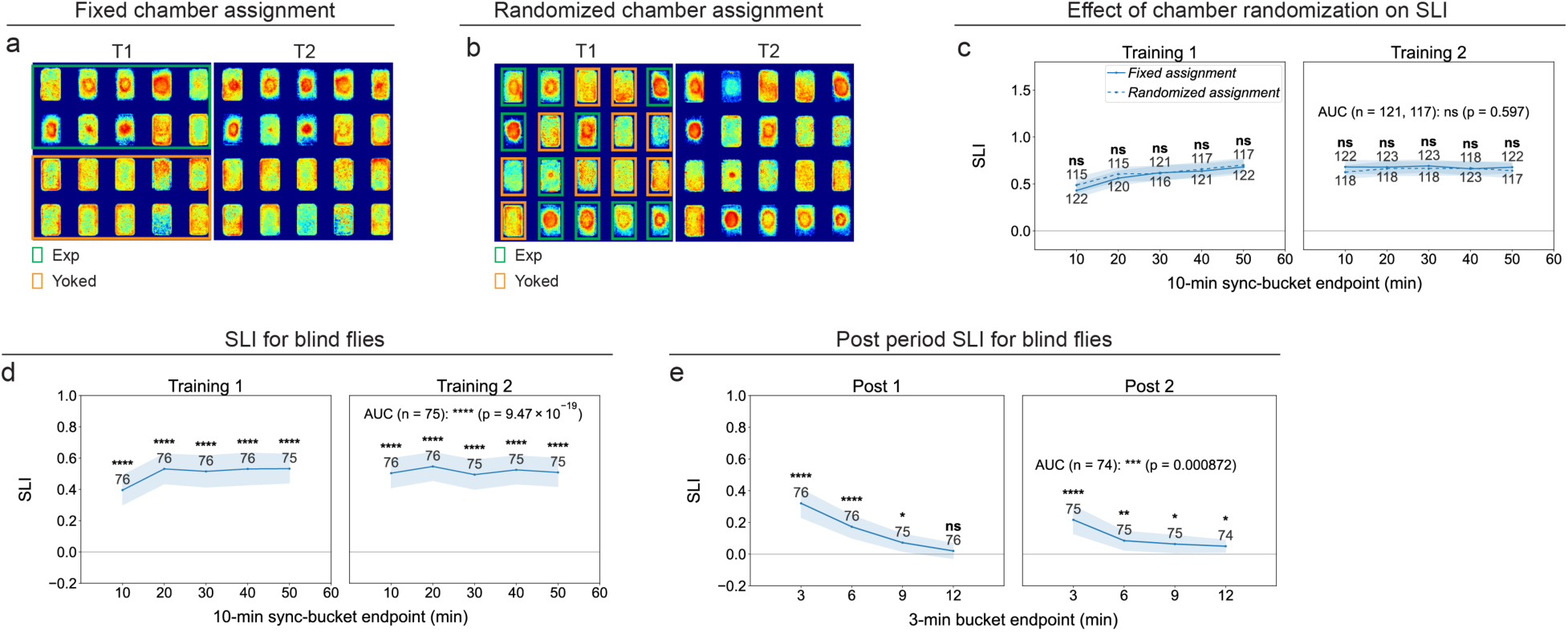
Chamber assignment and blindness do not affect flies’ learning. **a,** Positional heatmaps of *0273>CsChrimson* experimental flies and their paired yoked controls during Training 1 (T1) and Training 2 (T2) in the flat chamber with fixed chamber assignments. **b,** Positional heatmaps of *0273>CsChrimson* experimental flies and their paired yoked controls during T1 and T2 in the flat chamber with randomized chamber assignments. **c,** SLI of *0273>CsChrimson* flies with fixed versus randomized chamber assignments during Training sessions in the flat chamber. **d,** SLI of blind (*norpA; 0273>CsChrimson*) flies during Training sessions in the flat chamber. **e,** SLI of blind flies during Post 1 and Post 2 periods in the flat chamber. Unless otherwise stated, line plots show means and shaded 95% confidence intervals; numbers indicate sample sizes for each sync bucket. Throughout the paper, *: *p* < 0.05; **: *p* < 0.01; ***: *p* < 0.001; ****: *p* < 0.0001.

**Extended Data Fig. 3.**
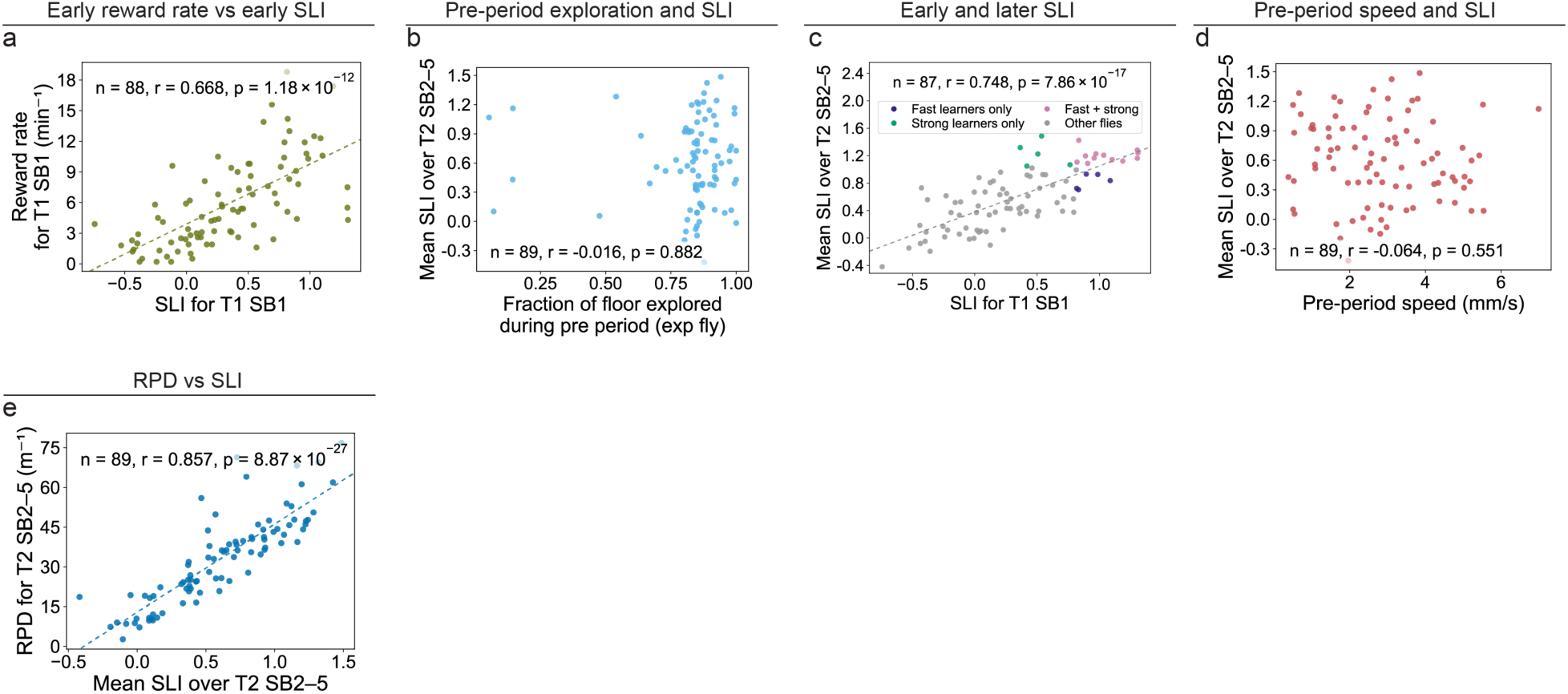
Correlations between SLI and other learning relevant parameters of *0273>CsChrimson* flies in the flat chamber. **a,** Correlation between SLI and reward rate of *0273>CsChrimson* flies during Training 1 (T1) SB1. **b,** Correlation between fraction of floor explored during the Pre period and mean SLI over Training 2 (T2) SB2–SB5 in *0273>CsChrimson* flies. **c,** Correlation between SLI in T1 SB1 and mean SLI over T2 SB2–SB5 in *0273>CsChrimson* flies. **d,** Correlation between mean locomotion speed during the Pre period and mean SLI over T2 SB2–SB5 in *0273>CsChrimson* flies. **e,** Correlation between mean SLI and rewards per distance traveled (RPD) over T2 SB2–SB5 in *0273>CsChrimson* flies. Pearson’s product-moment correlation was used in all panels.

**Extended Data Fig. 4.**
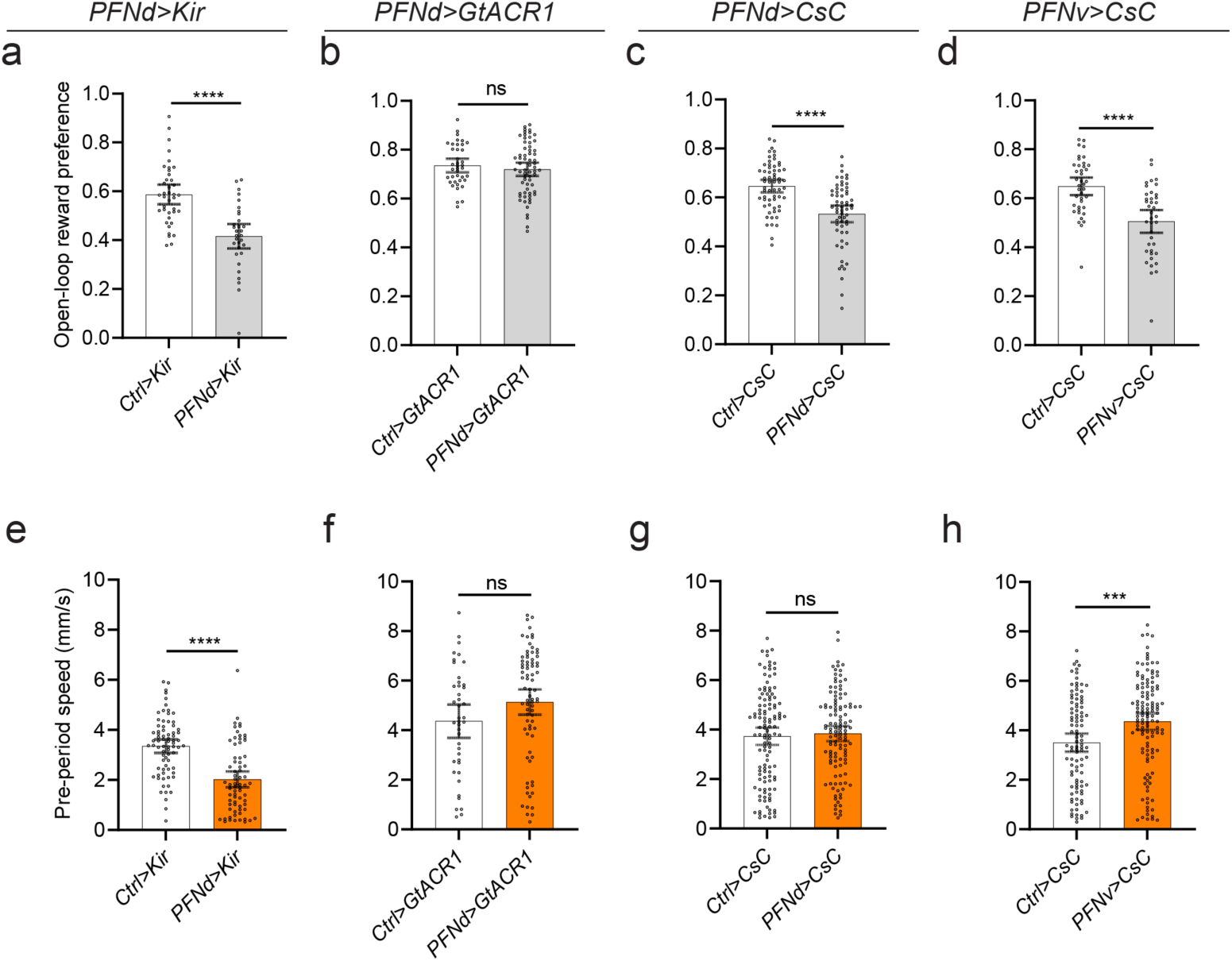
Manipulating PFNd and PFNv neurons does not consistently alter reward preference or innate locomotion speed. **a,** Open-loop reward preference of *Ctrl>Kir* (n = 39) versus *PFNd>Kir* (n = 31) flies. **b,** Open-loop reward preference of *Ctrl>GtACR1* (n = 39) versus *PFNd>GtACR1* (n = 64) flies. **c,** Open-loop reward preference of *Ctrl>CsChrimson* (n = 60) versus *PFNd>CsChrimson* (n = 60) flies. **d,** Open-loop reward preference of *Ctrl>CsChrimson* (n = 40) versus *PFNv>CsChrimson* (n = 39) flies. **e,** Mean locomotion speed of *Ctrl>Kir* (n = 74) versus *PFNd>Kir* (n = 70) flies during the Pre period. **f,** Mean locomotion speed of *Ctrl>GtACR1* (n = 42) versus *PFNd>GtACR1* (n = 77) flies. **g,** Mean locomotion speed of *Ctrl>CsChrimson* (n = 119) versus *PFNd>CsChrimson* (n = 118) flies. **h,** Mean locomotion speed of *Ctrl>CsChrimson* (n = 99) versus *PFNv>CsChrimson* (n = 124) flies. Error bars indicate 95% confidence intervals around the mean. Throughout the paper, *: *p* < 0.05; **: *p* < 0.01; ***: *p* < 0.001; ****: *p* < 0.0001.

**Extended Data Fig. 5.**
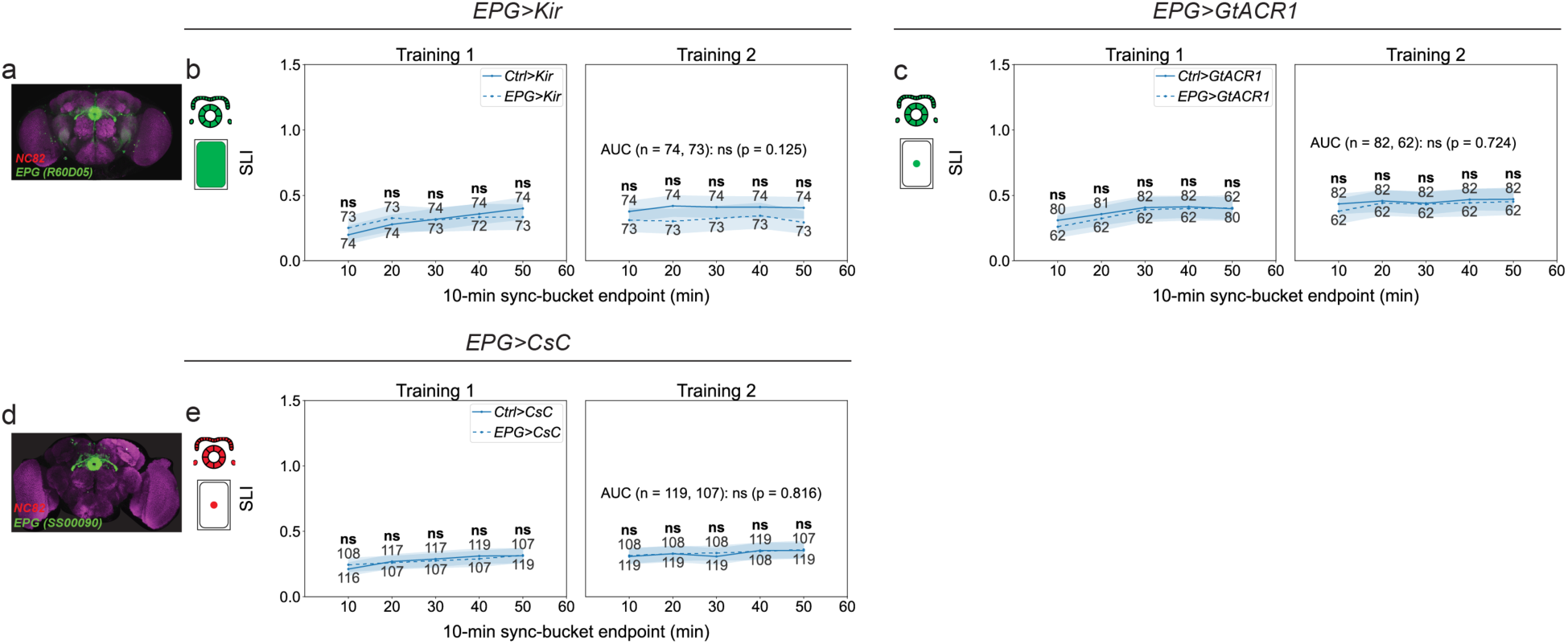
Manipulating EPG neurons involved in allocentric heading representation does not significantly affect spatial learning. **a,** Expression pattern of the *LexA* driver line labeling EPG neurons (*R60D05-LexA*). **b,** Left: Schematic of EPG neurons. Right: SLI of *Ctrl>Kir* versus *EPG>Kir* flies in the flat chamber. **c,** SLI of *Ctrl>GtACR1* versus *EPG>GtACR1* flies in the flat chamber. **d,** Expression pattern of the split-GAL4 driver labeling EPG neurons (*SS00090*). **e,** SLI of *Ctrl>CsChrimson* versus *EPG>CsChrimson* flies in the flat chamber. Unless otherwise stated, line plots show means and shaded 95% confidence intervals; numbers indicate sample sizes for each sync bucket. Throughout the paper, *: *p* < 0.05; **: *p* < 0.01; ***: *p* < 0.001; ****: *p* < 0.0001.

**Extended Data Fig. 6.**
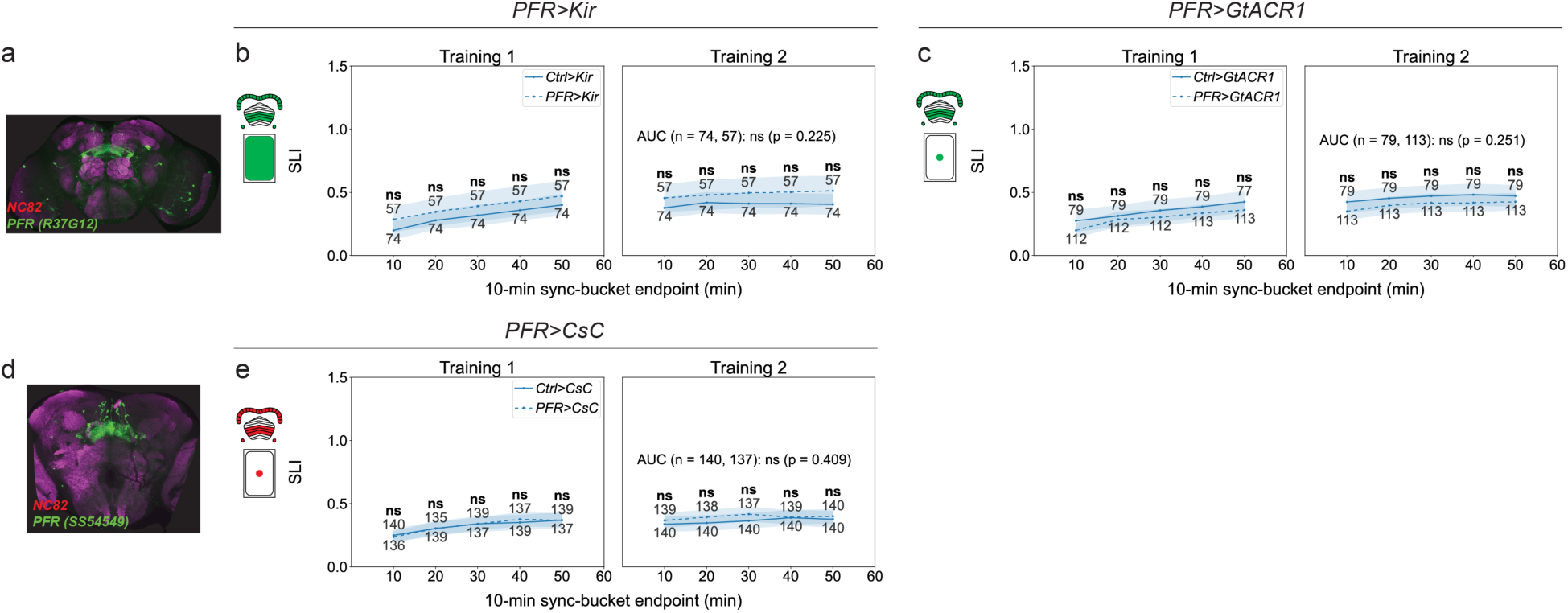
Manipulating PFR neurons involved in allocentric translational velocity representation does not significantly affect spatial learning. **a,** Expression pattern of the *LexA* driver line labeling PFR neurons (*R37G12-LexA*). **b,** Left: Schematic of PFR neurons. Right: SLI of *Ctrl>Kir* versus *PFR>Kir* flies in the flat chamber. **c,** SLI of *Ctrl>GtACR1* versus *PFR>GtACR1* flies in the flat chamber. **d,** Expression pattern of the split-GAL4 driver labeling PFR neurons (*SS54549*). **e,** SLI of *Ctrl>CsChrimson* versus *PFR>CsChrimson* flies in the flat chamber. Unless otherwise stated, line plots show means and shaded 95% confidence intervals; numbers indicate sample sizes for each sync bucket. Throughout the paper, *: *p* < 0.05; **: *p* < 0.01; ***: *p* < 0.001; ****: *p* < 0.0001.

**Extended Data Fig. 7.**
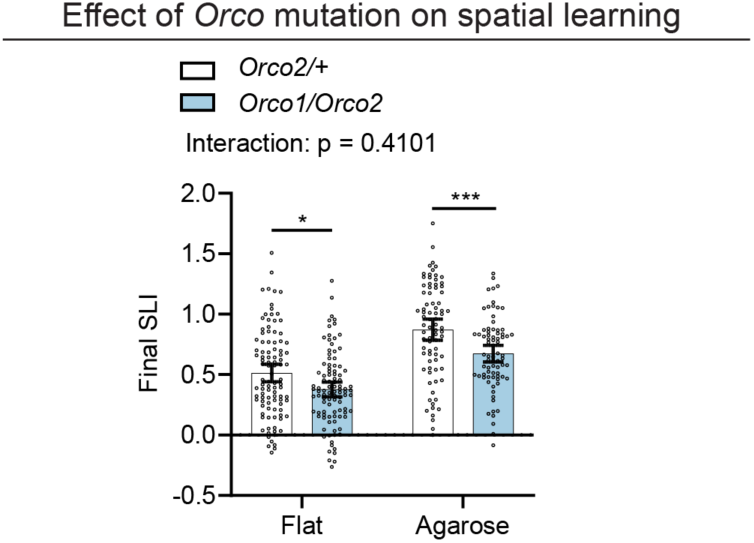
Orco is required for spatial learning in both flat and agarose chambers. **a,** Left two columns: Final SLI of *Orco2/+* (n = 99) and *Orco1/Orco2* (n = 98) *flies* in the flat chamber. Right two columns: Final SLI of *Orco2/+* (n = 79) and *Orco1/Orco2* (n = 76) flies in the agarose chamber. Two-way ANOVA was used to test the interaction between genotype and chamber type, followed by Šídák’s multiple comparisons test. Error bars indicate 95% confidence intervals around the mean. Throughout the paper, *: *p* < 0.05; **: *p* < 0.01; ***: *p* < 0.001; ****: *p* < 0.0001.

**Extended Data Fig. 8.**
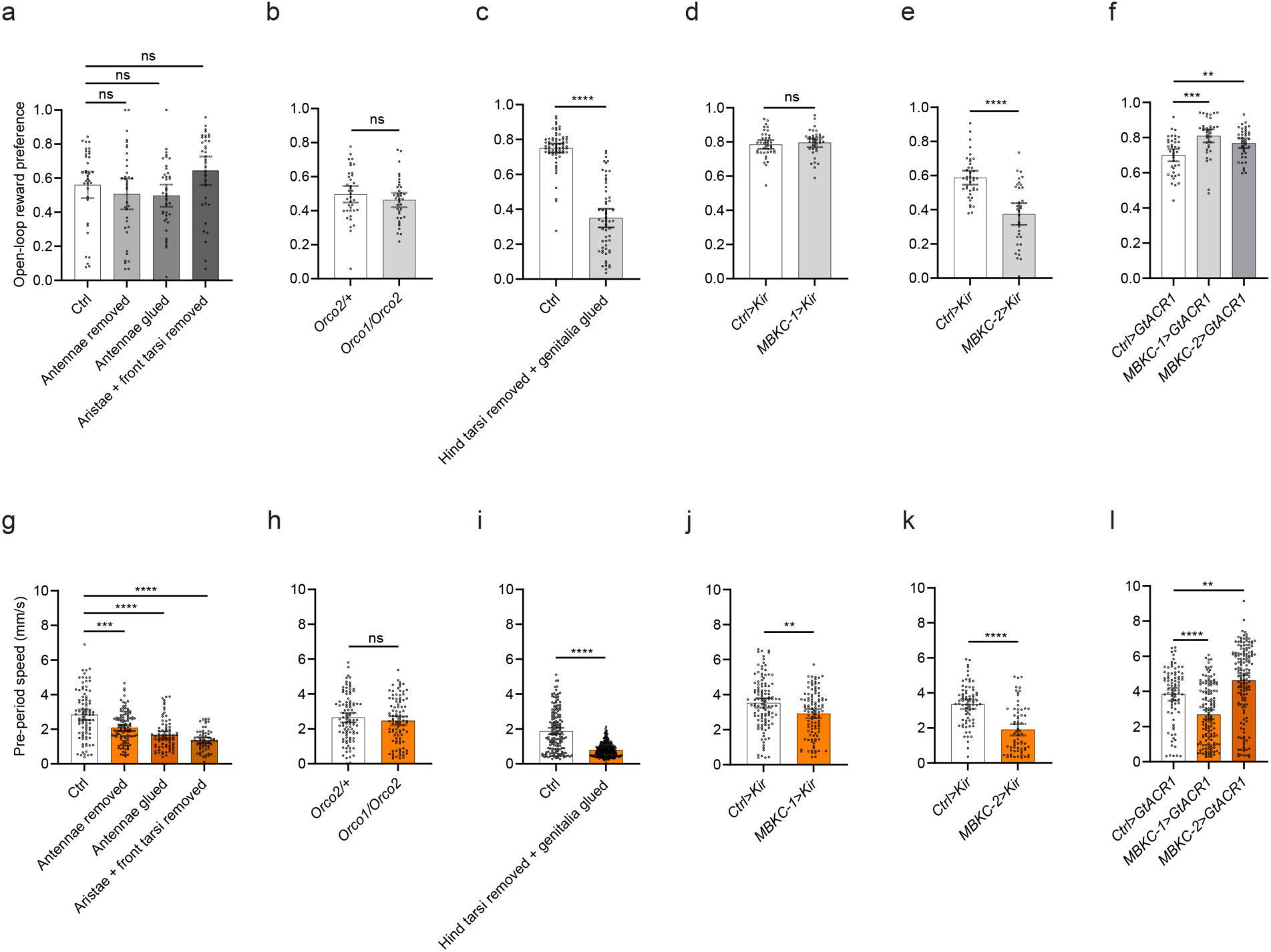
Manipulations affecting olfactory signaling and scent deposition do not consistently alter reward preference or innate locomotion speed. **a,** Open-loop reward preference of control flies (n = 36), antennae-removed flies (n = 36), antennae-glued flies (n = 39), and flies with aristae and front tarsi removed (n = 35). **b,** Open-loop reward preference of *Orco2/+* (n = 38) versus *Orco1/Orco2* (n = 37) flies. **c,** Open-loop reward preference of control flies (n = 68) and flies with hind tarsi removed and genitalia glued (n = 57). **d,** Open-loop reward preference of *Ctrl>Kir* (n = 38) versus *MBKC-1>Kir* (n = 37) flies. **e,** Open-loop reward preference of *Ctrl>Kir* (n = 39, same control group as for *PFNd>Kir*) versus *MBKC-2>Kir* (n = 34) flies. **f,** Open-loop reward preference of *Ctrl>GtACR1* (n = 39) versus *MBKC-1>GtACR1* (n = 36) and *MBKC-2>GtACR1* (n = 40) flies. **g,** Mean locomotion speed of control flies (n = 90), antennae-removed flies (n = 113), antennae-glued flies (n = 76), and flies with aristae and front tarsi removed (n = 63) during the Pre period. **h,** Mean locomotion speed of *Orco2/+* (n = 99) versus *Orco1/Orco2* (n = 98) flies during the Pre period. **i,** Mean locomotion speed of control flies (n = 159) and flies with hind tarsi removed and genitalia glued (n = 256) during the Pre period. **j,** Mean locomotion speed of *Ctrl>Kir* (n = 115) versus *MBKC-1>Kir* (n = 105) flies during the Pre period. **k,** Mean locomotion speed of *Ctrl>Kir* (n = 74, same control group as for *PFNd>Kir*) versus *MBKC-2>Kir* (n = 67) flies during the Pre period. **l,** Mean locomotion speed of *Ctrl>GtACR1* (n = 82) versus *MBKC-1>GtACR1* (n = 144) and *MBKC-2>GtACR1* (n = 153) flies during the Pre period. Welch’s ANOVA followed by Games–Howell post hoc comparisons was used in **a, f, g,** and **l**. Error bars indicate 95% confidence intervals around the mean. Throughout the paper, *: *p* < 0.05; **: *p* < 0.01; ***: *p* < 0.001; ****: *p* < 0.0001.

**Extended Data Fig. 9.**
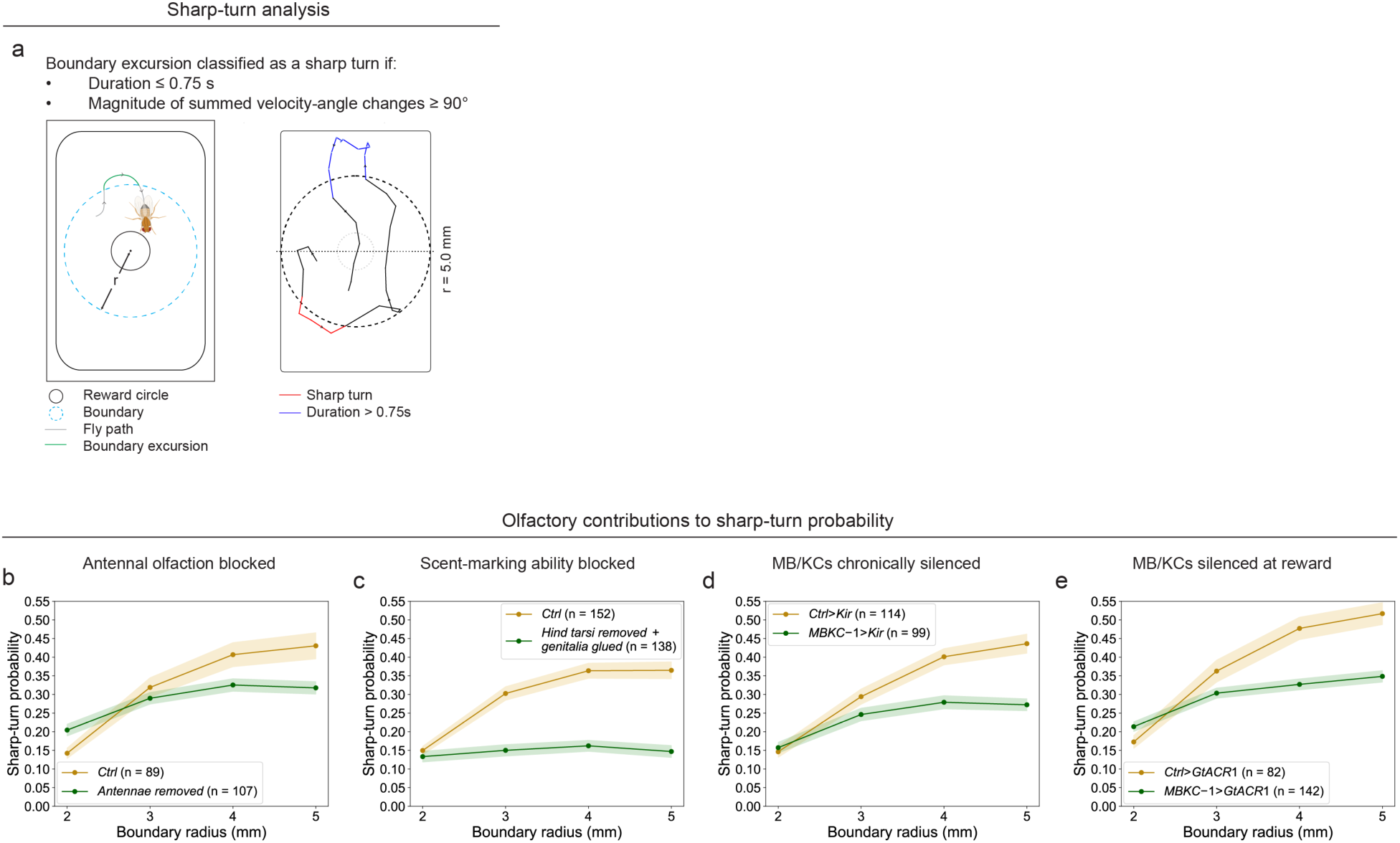
Effects of disrupting olfaction, scent deposition, and mushroom-body activity on sharp-turn probabilit. **a,** Schematic of the analysis used to identify sharp turns (≥90°) within 0.75 s after outward crossings of boundaries at different radii (2–5 mm) from the reward center (see **Methods**). **b,** Sharp-turn probability of control flies versus antennae-removed flies. **c,** Sharp-turn probability of control flies versus flies with hind tarsi removed and genitalia glued. **d,** Sharp-turn probability of *Ctrl>Kir* versus *MBKC-1>Kir* flies. **e,** Sharp-turn probability of *Ctrl>GtACR1* versus *MBKC-1>GtACR1* flies. Unless otherwise stated, line plots show means and shaded 95% confidence intervals. Throughout the paper, *: *p* < 0.05; **: *p* < 0.01; ***: *p* < 0.001; ****: *p* < 0.0001.

**Extended Data Fig. 10.**
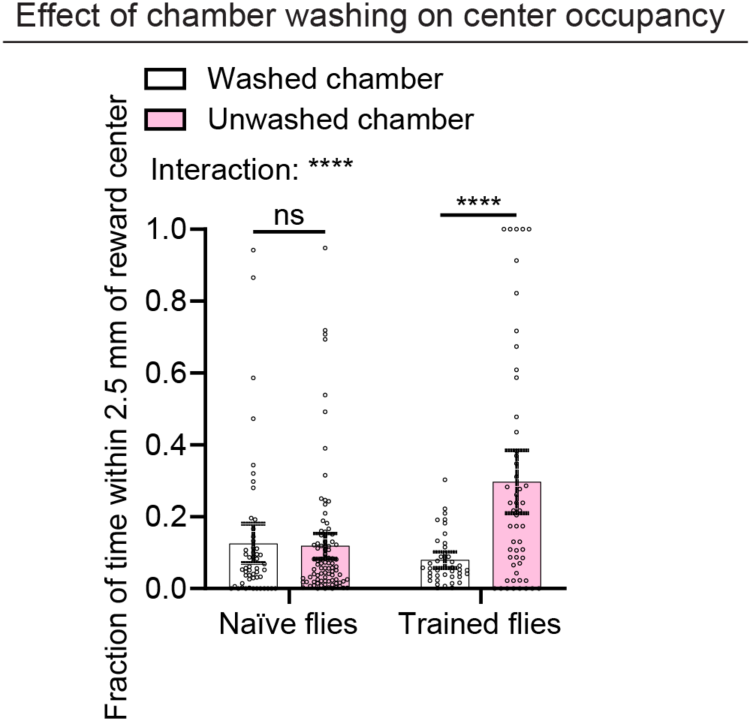
Training-dependent attraction to fly-deposited cues. **a,** Left two columns: Fraction of time naïve flies spent within 2.5 mm of the reward center in washed (n = 53) versus unwashed (n = 91) flat chambers. Right two columns: Fraction of time previously trained flies spent within 2.5 mm of the reward center in washed (n = 40) versus unwashed (n = 53) flat chambers. Unwashed chambers had been used for three Training sessions by another group of flies. Two-way ANOVA was used to test the interaction between training history and chamber washing, followed by Šídák’s multiple comparisons test. Error bars indicate 95% confidence intervals around the mean. Throughout the paper, *: *p* < 0.05; **: *p* < 0.01; ***: *p* < 0.001; ****: *p* < 0.0001.

**Extended Data Fig. 11.**
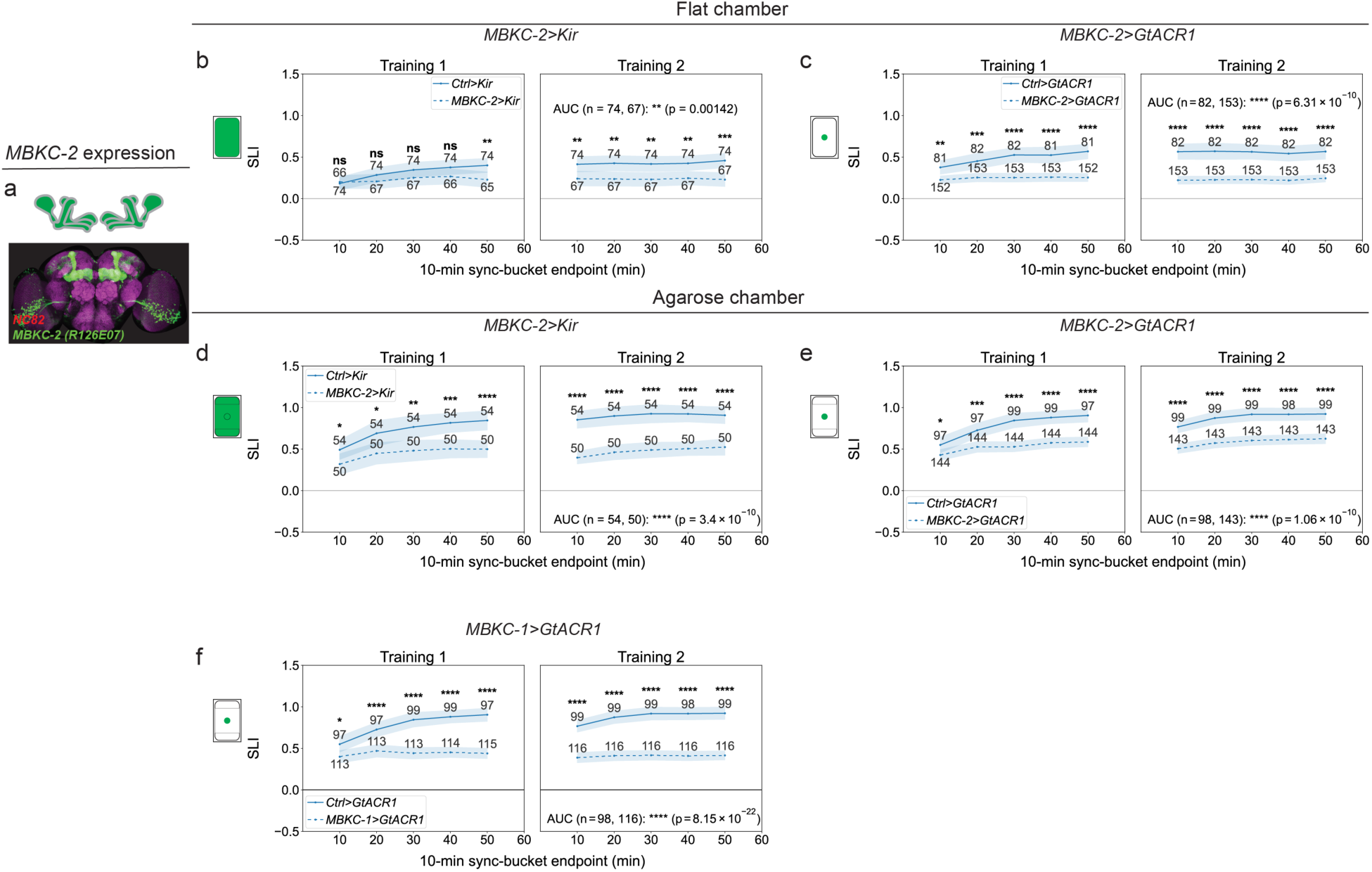
Silencing MBKCs with a second driver line also impairs spatial learning performance in both flat and agarose chambers. Expression pattern of the second *LexA* driver labeling MBKCs (*R26E07-LexA*). **b,** SLI of *Ctrl>Kir* versus *MBKC-2>Kir* flies in the flat chamber. **c,** SLI of *Ctrl>GtACR1* versus *MBKC-2>GtACR1* flies in the flat chamber. **d,** SLI of *Ctrl>Kir* versus *MBKC-2>Kir* flies in the agarose chamber. **e,** SLI of *Ctrl>GtACR1* versus *MBKC-2>GtACR1* flies in the agarose chamber. **f,** SLI of *Ctrl>GtACR1* (same control group as in **e**) versus *MBKC-1>GtACR1* flies in the agarose chamber. Unless otherwise stated, line plots show means and shaded 95% confidence intervals; numbers indicate sample sizes for each sync bucket. Throughout the paper, *: *p* < 0.05; **: *p* < 0.01; ***: *p* < 0.001; ****: *p* < 0.0001.

**Extended Data Fig. 12.**
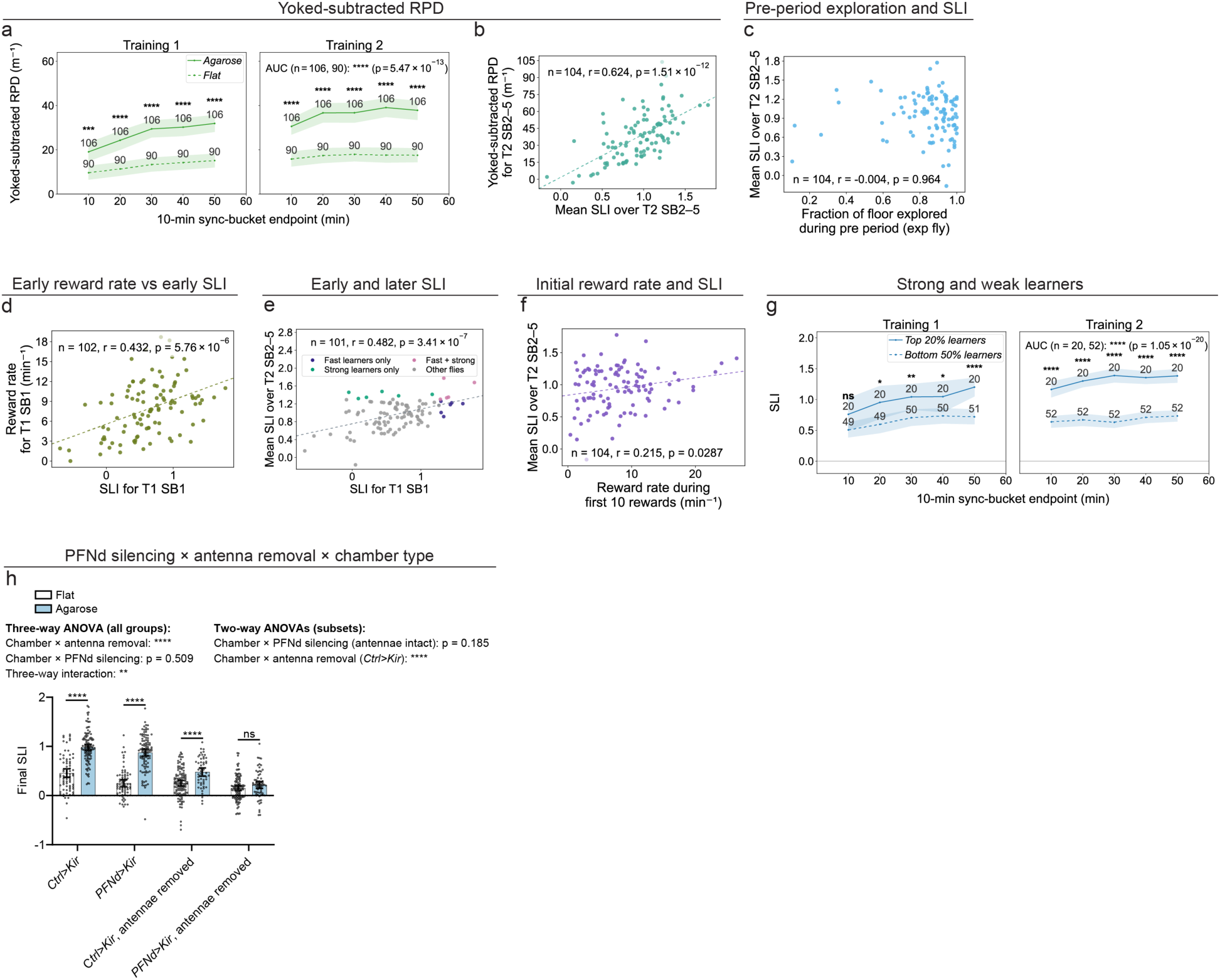
Additional analyses of spatial learning and its enhancement by agarose. **a,** Yoked-subtracted rewards per distance traveled (RPD) of *0273>CsChrimson* flies in the flat versus agarose chambers during Training sessions. **b,** Correlation between yoked-subtracted RPD and mean SLI over Training 2 (T2) SB2–SB5 in *0273>CsChrimson* flies in the agarose chamber. **c,** Correlation between fraction of floor explored during the Pre period and mean SLI over T2 SB2–SB5 in *0273>CsChrimson* flies in the agarose chamber. **d,** Correlation between SLI and reward rate in Training 1 (T1) SB1 in *0273>CsChrimson* flies in the agarose chamber. **e,** Correlation between SLI in T1 SB1 and mean SLI over T2 SB2–SB5 in *0273>CsChrimson* flies in the agarose chamber. **f,** Correlation between reward rate during the first 10 rewards and mean SLI over T2 SB2–SB5 in *0273>CsChrimson* flies in the agarose chamber. **g,** SLI of strong (top 20%) and weak (bottom 50%) learners among *0273>CsChrimson* flies during Training sessions in the agarose chamber. Learner classification was based on mean SLI over T2 SB2–SB5. **h,** First two columns: Final SLI of *Ctrl>Kir* flies in the flat (n = 74) and agarose (n = 110) chambers. Second two columns: Final SLI of *PFNd>Kir* flies in the flat (n = 70) and agarose (n = 100) chambers. Third two columns: Final SLI of antennae-removed *Ctrl>Kir* flies in the flat (n = 110) and agarose (n = 52) chambers. Last two columns: Final SLI of antennae-removed *PFNd>Kir* flies in the flat (n = 106) and agarose (n = 67) chambers. Three-way ANOVA was used to test interactions among chamber type, PFNd silencing, and antenna removal, followed by Šídák’s multiple comparisons test in **h**. Additional two-way ANOVAs were performed on subsets of the data. Pearson’s product-moment correlation was used in panels **b–f**. Unless otherwise stated, line plots show means and shaded 95% confidence intervals; numbers indicate sample sizes for each sync bucket. Throughout the paper, *: p < 0.05; **: p < 0.01; ***: p < 0.001; ****: p < 0.0001.

**Extended Data Fig. 13.**
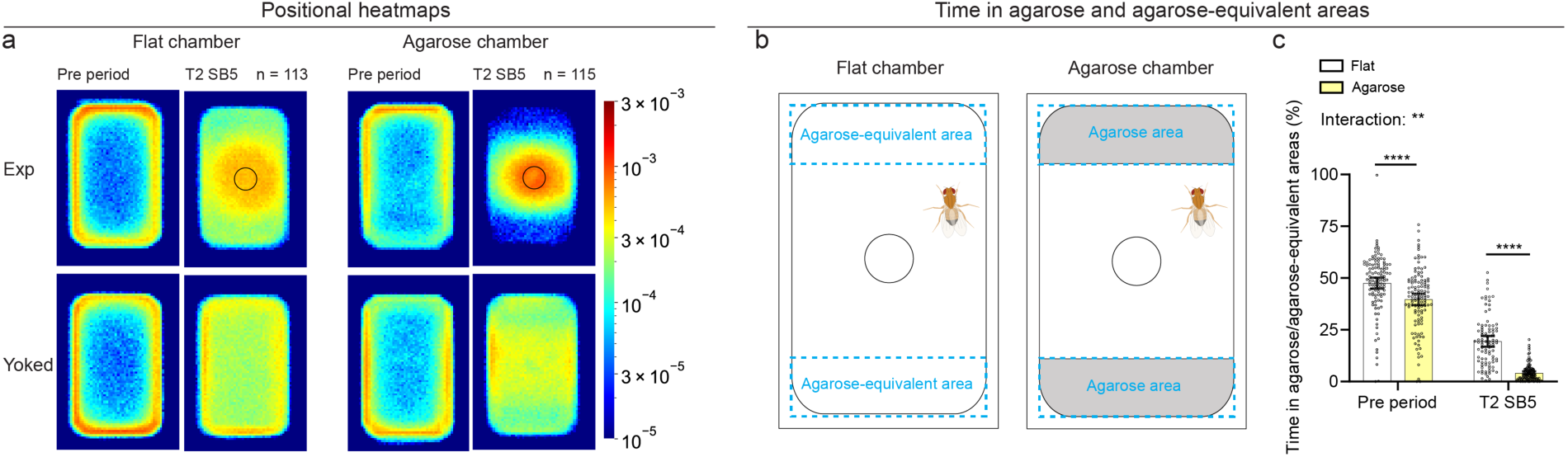
Learning-induced avoidance of agarose-occupied areas. **a,** Positional heatmaps of experimental flies and their paired yoked controls during the last 10 min of the Pre period and Training 2 (T2) SB5. Left: Flat chamber. Right: Agarose chamber. **b,** Schematic of agarose-equivalent areas in the flat chamber (left) and agarose-occupied areas in the agarose chamber (right). **c,** Left two columns: Percentage of time spent in agarose-equivalent areas in the flat chamber (n = 113) versus agarose-occupied areas in the agarose chamber (n = 115) during the last 10 min of the Pre period. The difference was statistically significant but small. Right two columns: Percentage of time spent in agarose-equivalent areas in the flat chamber (n = 89) versus agarose-occupied areas in the agarose chamber (n = 105) during T2 SB5. (Pre-period data included flies with valid tracking, whereas T2 SB5 data were restricted to flies with a defined SLI in that sync bucket.) Two-way ANOVA was used to test the interaction between chamber type and experimental period, followed by Šídák’s multiple comparisons test. Error bars indicate 95% confidence intervals around the mean. Throughout the paper, *: *p* < 0.05; **: *p* < 0.01; ***: *p* < 0.001; ****: *p* < 0.0001.

**Extended Data Fig. 14.**
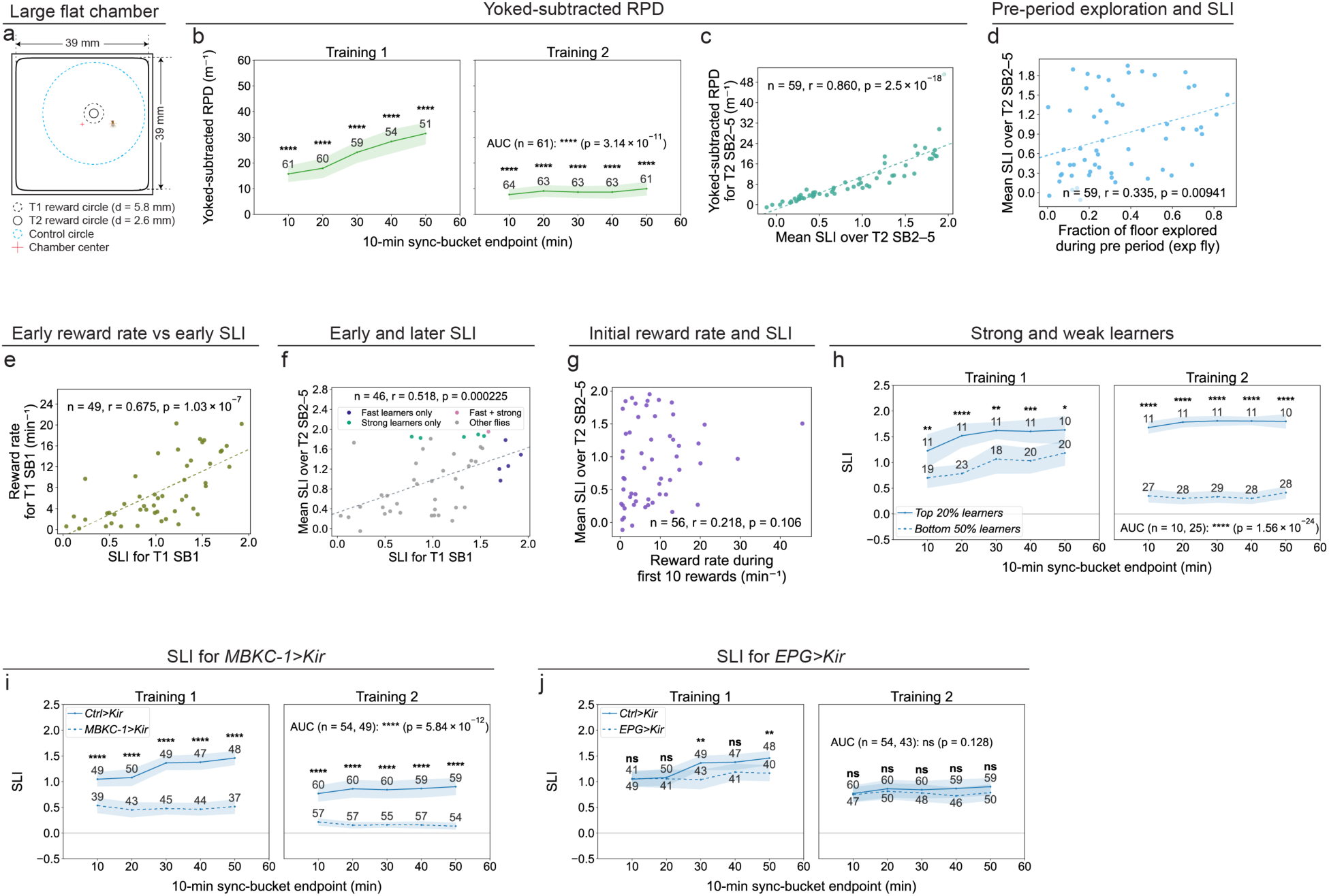
Additional analyses of spatial learning in the large flat chamber. **a,** Schematic of the large flat chamber. **b,** Yoked-subtracted rewards per distance traveled (RPD) of *0273>CsChrimson* flies during Training sessions in the large flat chamber. **c,** Correlation between yoked-subtracted RPD and mean SLI over Training 2 (T2) SB2–SB5 in *0273>CsChrimson* flies in the large flat chamber. **d,** Correlation between fraction of floor explored during the Pre period and mean SLI over T2 SB2–SB5 in *0273>CsChrimson* flies in the large flat chamber. **e,** Correlation between SLI and reward rate in Training 1 (T1) SB1 in *0273>CsChrimson* flies in the large flat chamber. **f,** Correlation between SLI in T1 SB1 and mean SLI over T2 SB2–SB5 in *0273>CsChrimson* flies in the large flat chamber. **g,** Correlation between reward rate during the first 10 rewards and mean SLI over T2 SB2–SB5 in *0273>CsChrimson* flies in the large flat chamber. **h,** SLI of strong (top 20%) and weak (bottom 50%) learners among *0273>CsChrimson* flies during Training sessions in the large flat chamber. Learner classification was based on mean SLI over T2 SB2–SB5. **i,** SLI of *Ctrl>Kir* versus *MBKC-1>Kir* flies in the large flat chamber. **j,** SLI of *Ctrl>Kir* versus *EPG>Kir* flies in the large flat chamber. Pearson’s product-moment correlation was used in **c–g**. Unless otherwise stated, line plots show means and shaded 95% confidence intervals; numbers indicate sample sizes for each sync bucket. Throughout the paper, *: *p* < 0.05; **: *p* < 0.01; ***: *p* < 0.001; ****: *p* < 0.0001.

**Extended Data Fig. 15.**
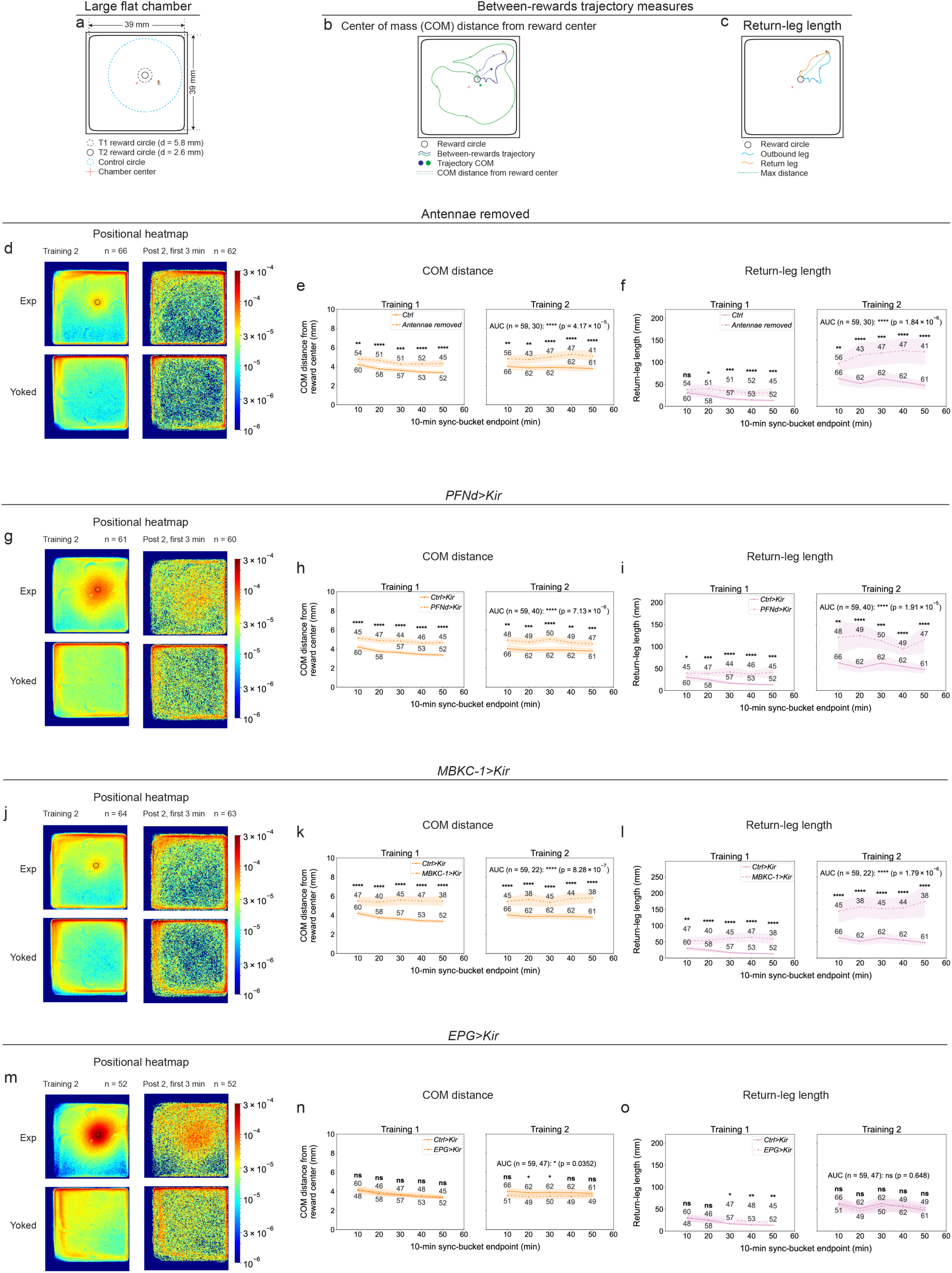
Antennae, PFNd neurons, and MBKCs, but not EPG neurons, are required for spatial learning in the large flat chamber. **a,** Schematic of the large flat chamber. **b,** Schematic of center-of-mass (COM) distance from the reward center (see **Methods**). **c,** Schematic of return-leg length for a between-rewards trajectory (see **Methods**). **d,** Positional heatmaps of experimental flies and their paired yoked controls during Training 2 (T2) and during the first 3 min of the Post 2 period for antennae-removed flies in the large flat chamber. **e,** COM distance from the reward center of control flies versus antennae-removed flies in the large flat chamber. **f,** Return-leg length of control flies versus antennae-removed flies in the large flat chamber. **g,** Positional heatmaps of experimental flies and their paired yoked controls during T2 and during the first 3 min of the Post 2 period for *PFNd>Kir* flies in the large flat chamber. **h,** COM distance from the reward center of *Ctrl>Kir* versus *PFNd>Kir* flies in the large flat chamber. **i,** Return-leg length of *Ctrl>Kir* versus *PFNd>Kir* flies in the large flat chamber. **j,** Positional heatmaps of experimental flies and their paired yoked controls during T2 and during the first 3 min of the Post 2 period for *MBKC-1>Kir* flies in the large flat chamber. **k,** COM distance from the reward center of *Ctrl>Kir* versus *MBKC-1>Kir* flies in the large flat chamber. **l,** Return-leg length of *Ctrl>Kir* versus *MBKC-1>Kir* flies in the large flat chamber. **m,** Positional heatmaps of experimental flies and their paired yoked controls during T2 and during the first 3 min of the Post 2 period for *EPG>Kir* flies in the large flat chamber. **n,** COM distance from the reward center of *Ctrl>Kir* versus *EPG>Kir* flies in the large flat chamber. **o,** Return-leg length of *Ctrl>Kir* versus *EPG>Kir* flies in the large flat chamber. Unless otherwise stated, line plots show means and shaded 95% confidence intervals; numbers indicate sample sizes for each sync bucket. Throughout the paper, *: *p* < 0.05; **: *p* < 0.01; ***: *p* < 0.001; ****: *p* < 0.0001.

**Extended Data Fig. 16.**
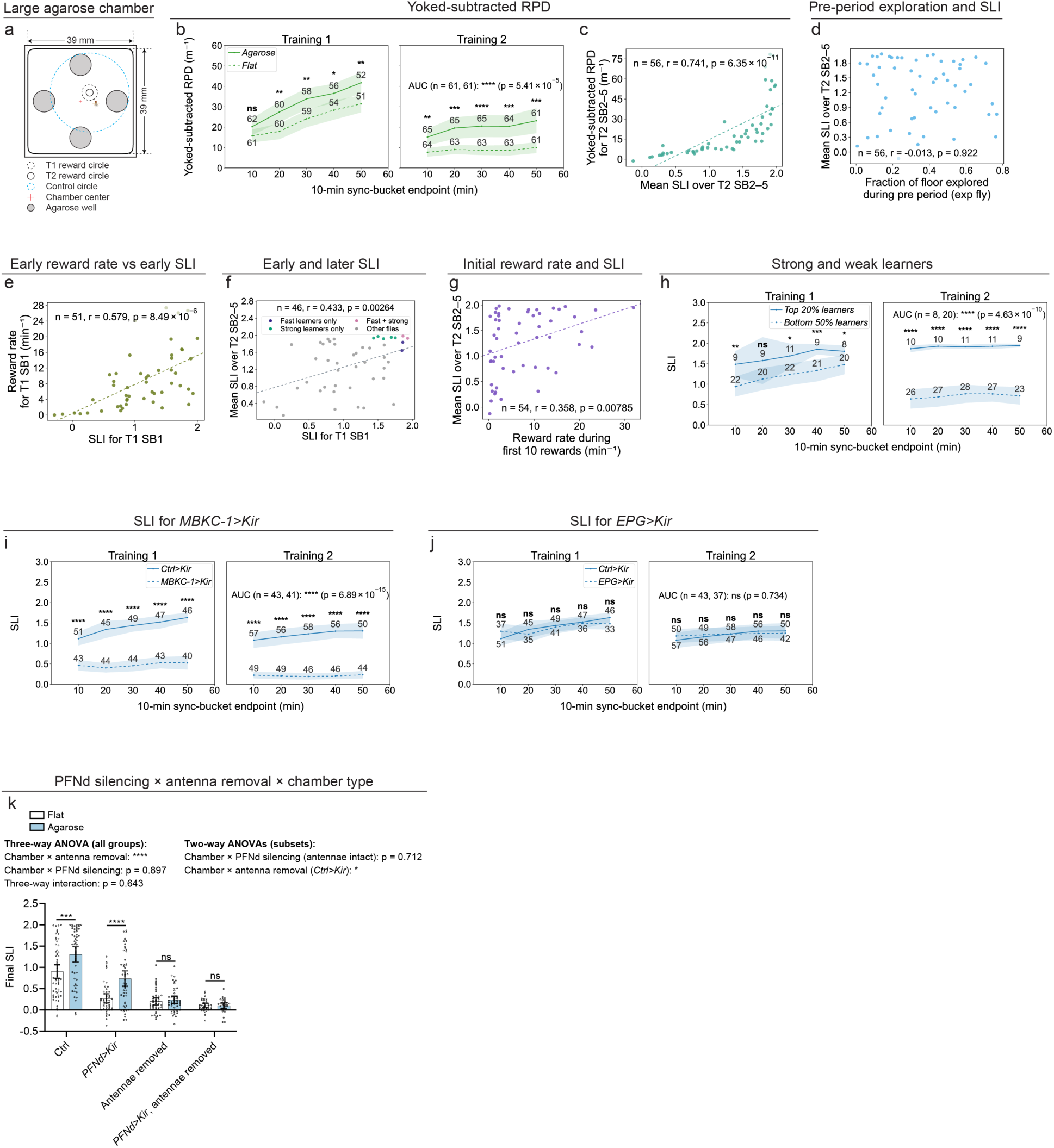
Additional analyses of spatial learning and its enhancement by agarose in large chambers. **a,** Schematic of the large agarose chamber. **b,** Yoked-subtracted rewards per distance traveled (RPD) of *0273>CsChrimson* flies in the large flat versus large agarose chambers during Training sessions. **c,** Correlation between yoked-subtracted RPD and mean SLI over Training 2 (T2) SB2–SB5 in *0273>CsChrimson* flies in the large agarose chamber. **d,** Correlation between fraction of floor explored during the Pre period and mean SLI over T2 SB2–SB5 in *0273>CsChrimson* flies in the large agarose chamber. **e,** Correlation between SLI and reward rate in Training 1 (T1) SB1 in *0273>CsChrimson* flies in the large agarose chamber. **f,** Correlation between SLI in T1 SB1 and mean SLI over T2 SB2–SB5 in *0273>CsChrimson* flies in the large agarose chamber. **g,** Correlation between reward rate during the first 10 rewards and mean SLI over T2 SB2–SB5 in *0273>CsChrimson* flies in the large agarose chamber. **h,** SLI of strong (top 20%) and weak (bottom 50%) learners among *0273>CsChrimson* flies during Training sessions in the large agarose chamber. Learner classification was based on mean SLI over T2 SB2–SB5. **i,** SLI of *Ctrl>Kir* versus *MBKC-1>Kir* flies in the large agarose chamber. **j,** SLI of *Ctrl>Kir* versus *EPG>Kir* flies in the large agarose chamber. **k,** First two columns: Final SLI of *Ctrl>Kir* flies in the large flat (n = 59) and large agarose (n = 50) chambers. Second two columns: Final SLI of *PFNd>Kir* flies in the large flat (n = 47) and large agarose (n = 50) chambers. Third two columns: Final SLI of antennae-removed *Ctrl>Kir* flies in the large flat (n = 45) and large agarose (n = 45) chambers. Last two columns: Final SLI of antennae-removed *PFNd>Kir* flies in the large flat (n = 33) and large agarose (n = 29) chambers. Pearson’s product-moment correlation was used in **c–g**. Three-way ANOVA was used to test interactions among chamber type, PFNd silencing, and antenna removal, followed by Šídák’s multiple comparisons test in **k**. Additional two-way ANOVAs were performed on subsets of the data. Unless otherwise stated, line plots show means and shaded 95% confidence intervals; numbers indicate sample sizes for each sync bucket. Throughout the paper, *: *p* < 0.05; **: *p* < 0.01; ***: *p* < 0.001; ****: *p* < 0.0001.

**Extended Data Fig. 17.**
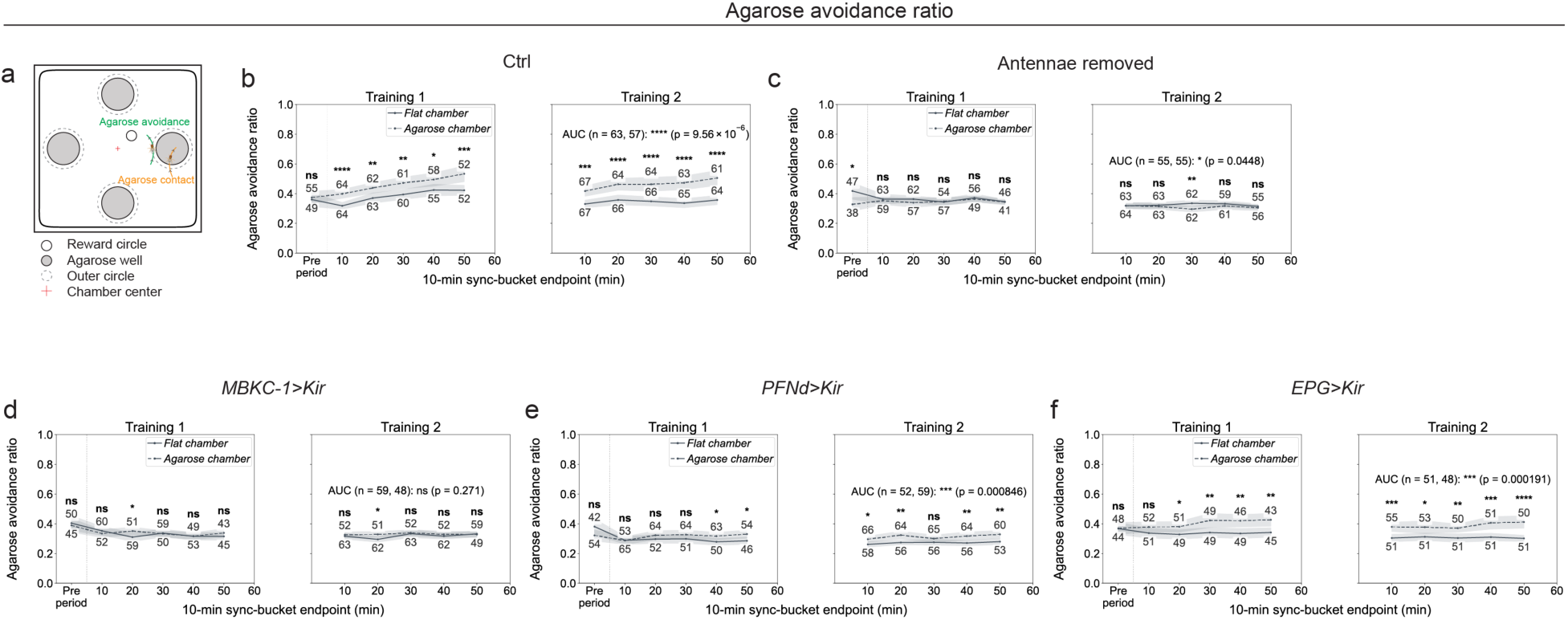
Analyses of agarose avoidance in the large agarose chamber. **a,** Schematic of the agarose avoidance ratio calculation (see **Methods**). **b,** Agarose avoidance ratio of *0273>CsChrimson* flies in the large agarose versus large flat chambers. **c,** Agarose avoidance ratio of antennae-removed flies in the large agarose versus large flat chambers. **d,** Agarose avoidance ratio of *MBKC-1>Kir* flies in the large agarose versus large flat chambers. **e,** Agarose avoidance ratio of *PFNd>Kir* flies in the large agarose versus large flat chambers. **f,** Agarose avoidance ratio of *EPG>Kir* flies in the large agarose versus large flat chambers. Unless otherwise stated, line plots show means and shaded 95% confidence intervals; numbers indicate sample sizes for each sync bucket. Throughout the paper, *: *p* < 0.05; **: *p* < 0.01; ***: *p* < 0.001; ****: *p* < 0.0001.

**Extended Data Fig. 18.**
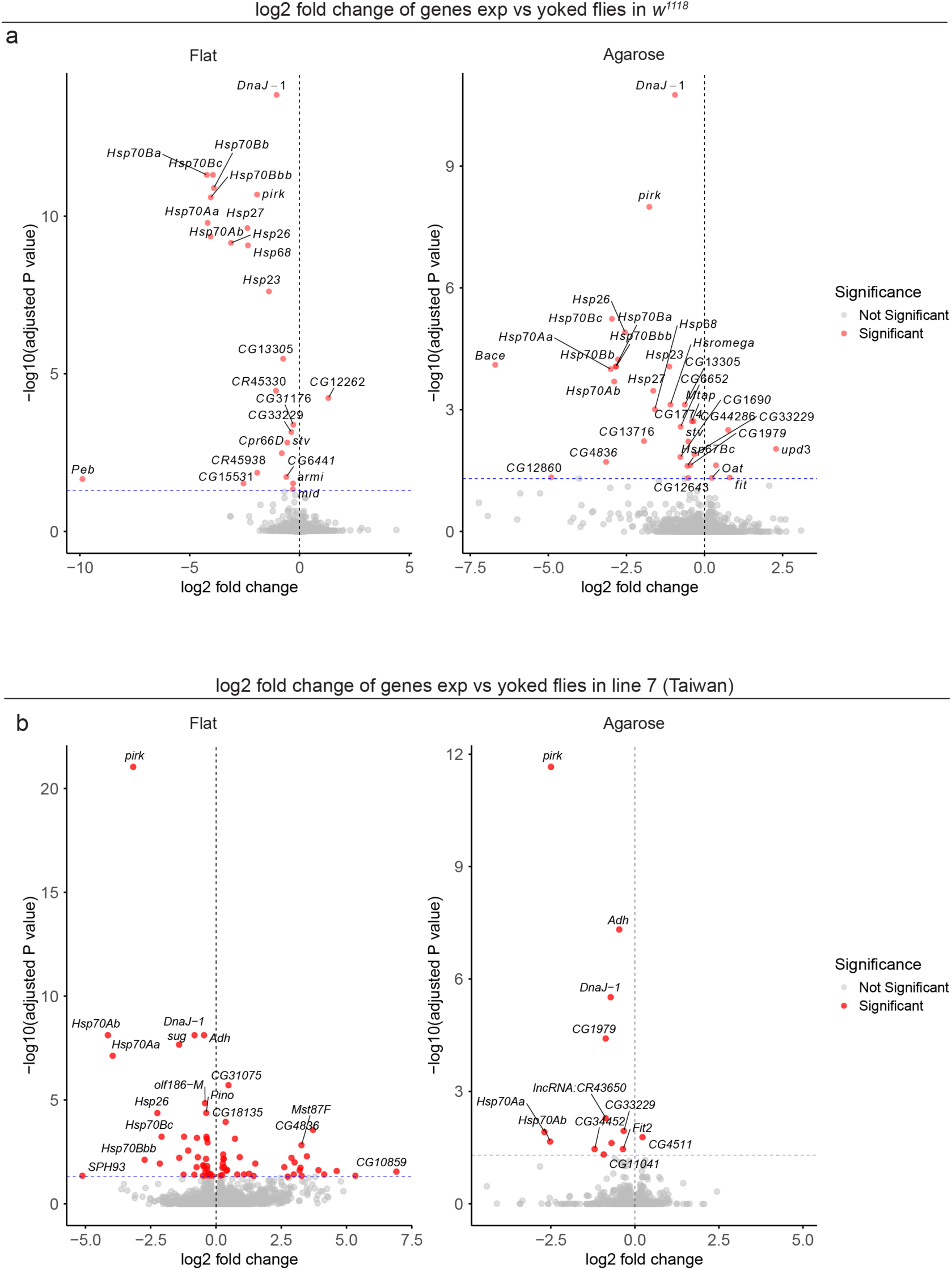
Volcano plots of differential gene expression between experimental flies and their paired yoked controls in the *w^1118^* and wild-caught Line 7 (Taiwan) genetic backgrounds. **a,** Volcano plots showing differential gene expression in the flat (left) and agarose (right) chambers in the *w^1118^*genetic background. The horizontal dotted line indicates an adjusted *p*-value threshold of 0.05, and the vertical dotted line indicates a log_2_ fold change of 0. **b,** Corresponding volcano plots for the wild-caught Line 7 (Taiwan) genetic background.

**Extended Data Fig. 19.**
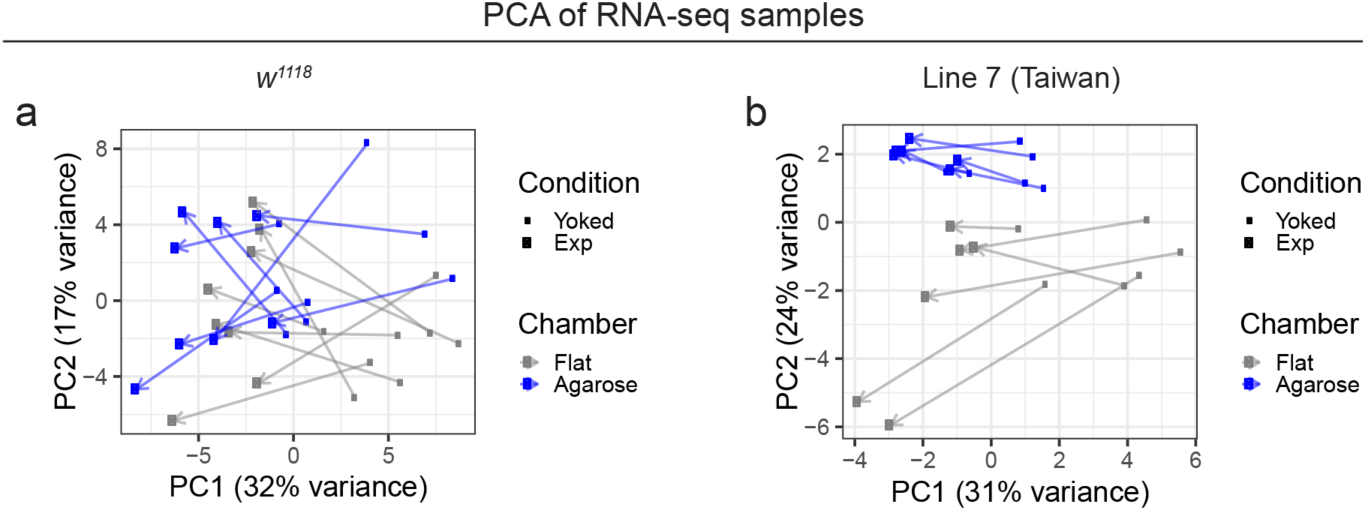
Principal component analysis of head transcriptomes from experimental and yoked flies in flat and agarose chambers. **a,** Principal component analysis (PCA) of head transcriptomes from experimental flies and their paired yoked controls in flat and agarose chambers immediately after Training 3 in the lab strain genetic background. **b,** Corresponding PCA of head transcriptomes in the wild-caught Line 7 (Taiwan) genetic background. For each background, the 1,000 genes with the smallest adjusted *p* values, selected from the combined differential expression analyses of the flat and agarose chambers, were used for PCA.

**Extended Data Fig. 20.**
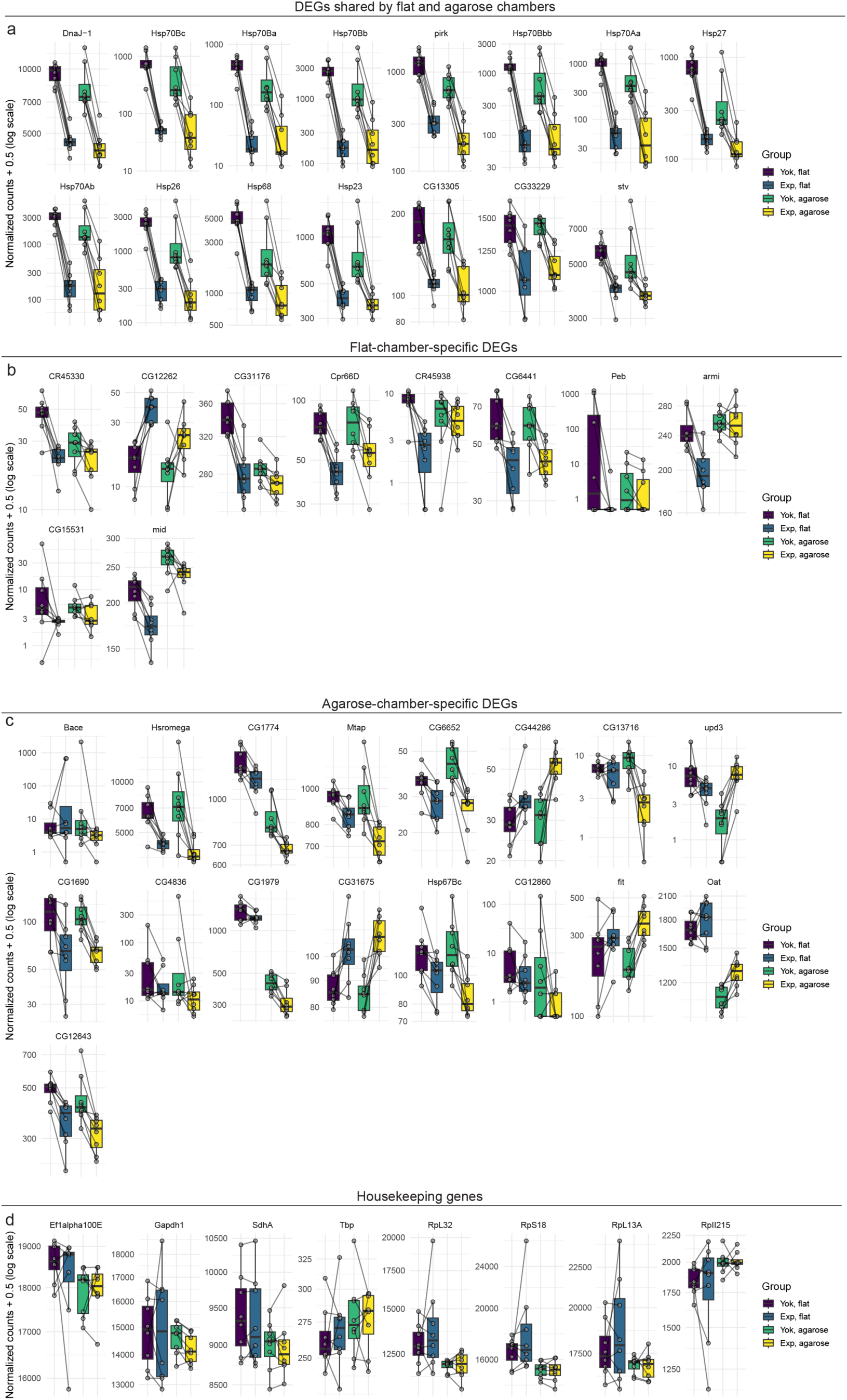
Normalized counts for identified DEGs and housekeeping genes in the *w^1118^*background. **a,** Normalized counts of 15 DEGs shared between flat and agarose chambers. **b,** Normalized counts of 10 flat-chamber-specific DEGs. **c,** Normalized counts of 17 agarose-chamber-specific DEGs. **d,** Normalized counts of eight housekeeping genes. Values are displayed on a logarithmic scale after adding 0.5 to the normalized counts.

**Extended Data Fig. 21.**
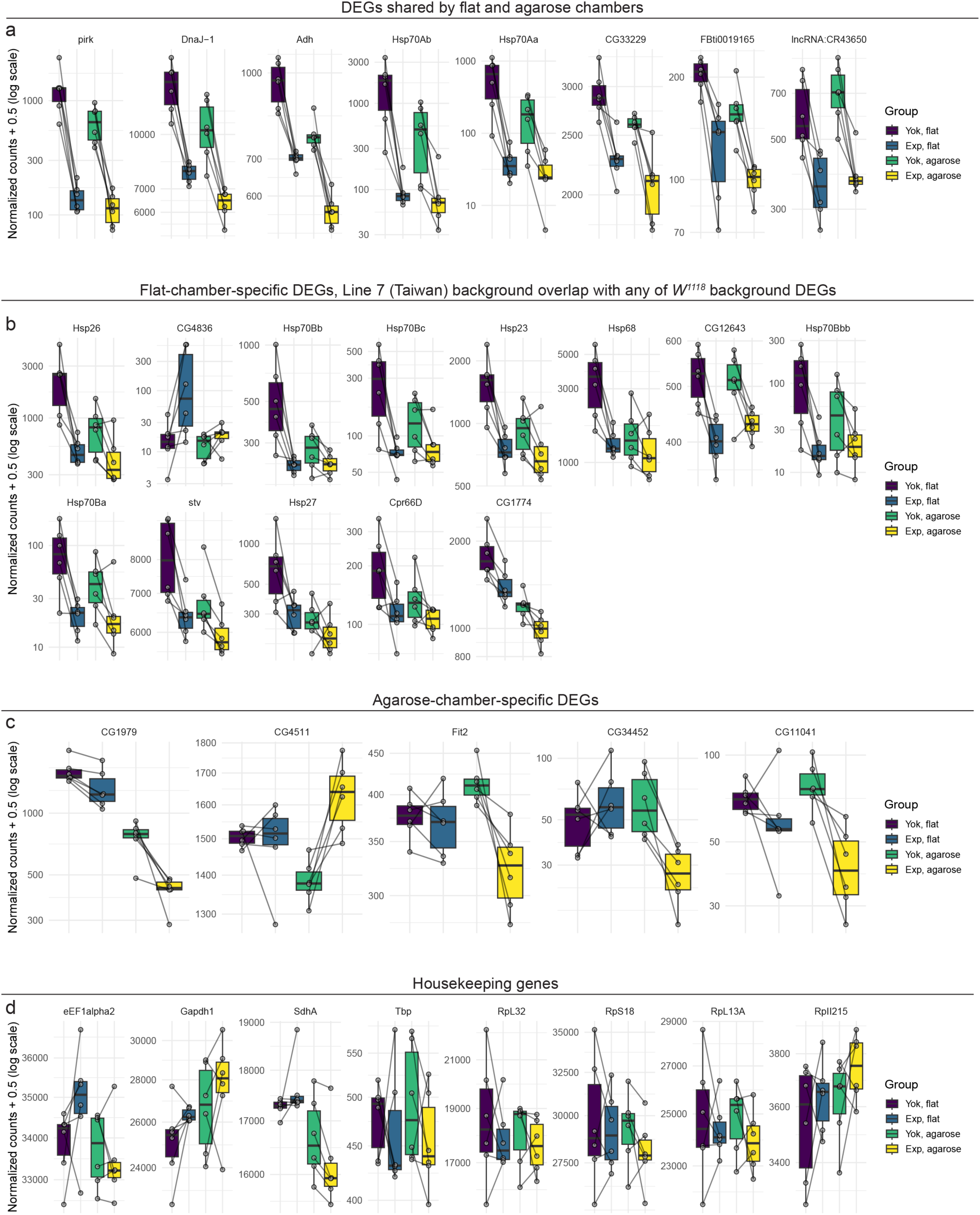
Normalized counts for identified DEGs and housekeeping genes in the wild-caught Line 7 (Taiwan) background. **a,** Normalized counts of eight DEGs shared between flat and agarose chambers. **b,** Normalized counts of 13 flat-chamber-specific DEGs also identified in the *w^1118^* background. **c,** Normalized counts of five agarose-chamber-specific DEGs. **d,** Normalized counts of eight housekeeping genes. Values are displayed on a logarithmic scale after adding 0.5 to the normalized counts.

**Extended Data Fig. 22.**
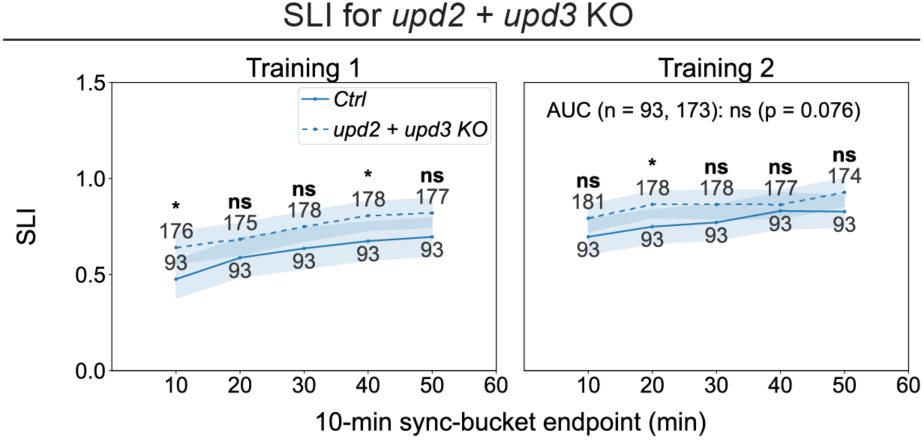
Double knockout of *upd2* and *upd3* does not significantly alter spatial learning performance. **a,** SLI of control flies versus *upd2 + upd3* knockout (KO) flies in the agarose chamber. Unless otherwise stated, line plots show means and shaded 95% confidence intervals; numbers indicate sample sizes for each sync bucket. Throughout the paper, *: *p* < 0.05; **: *p* < 0.01; ***: *p* < 0.001; ****: *p* < 0.0001.

**Extended Data Fig. 23.**
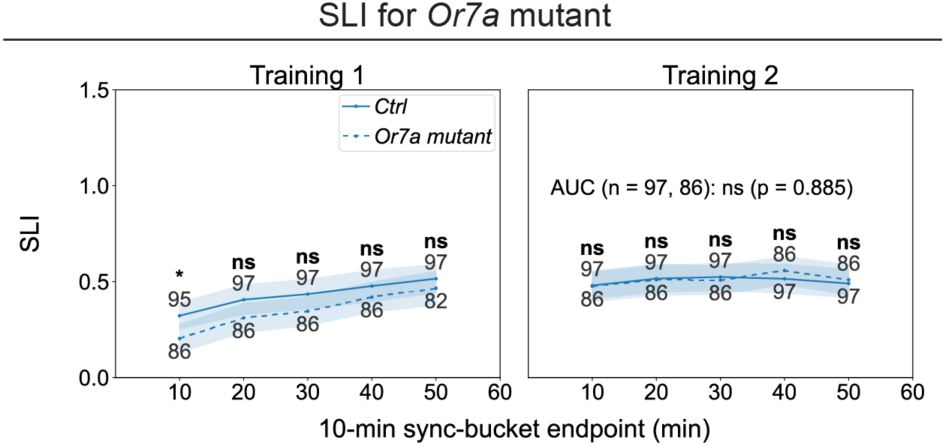
*Or7a* mutants do not show significant changes in spatial learning performance. **a,** SLI of control flies versus *Or7a* mutant flies in the flat chamber. Unless otherwise stated, line plots show means and shaded 95% confidence intervals; numbers indicate sample sizes for each sync bucket. Throughout the paper, *: *p* < 0.05; **: *p* < 0.01; ***: *p* < 0.001; ****: *p* < 0.0001.

## Supplementary Tables

**Supplementary Table 1. Reward rates across training sessions by genotype and experimental condition**

**Supplementary Table 2. Experimental–yoked differences in rewards per distance traveled by genotype and experimental condition**

**Supplementary Table 3. Sources of the stocks used in this study**

## Supplementary Videos

**Supplementary Video 1. High-throughput spatial learning task with *0273>CsChrimson* flies in the flat chamber**

Left half: Example video showing 10 experimental flies and their paired yoked control flies undergoing training in our high-throughput spatial learning task in the flat chamber. Right half: The real-time positional heatmap of corresponding flies during training. The video is at 2X play back speed.

**Supplementary Video 2. High-throughput spatial learning task with *0273>CsChrimson* flies in the agarose chamber**

Left half: Example video showing 10 experimental flies and their paired yoked control flies undergoing training in our high-throughput spatial learning task in the agarose chamber. Right half: The real-time positional heatmap of corresponding flies during training. The video is at 2X play back speed.

**Supplementary Video 3. Spatial learning task with *0273>CsChrimson* flies in the large agarose chamber**

Left half: Example video showing 2 experimental flies and their paired yoked control flies undergoing training in our spatial learning task in the large agarose chamber. Right half: The real-time positional heatmap of corresponding flies during training. The video is at 2X play back speed.

